# Metabolite therapy guided by liquid biopsy proteomics delays retinal neurodegeneration

**DOI:** 10.1101/764100

**Authors:** Katherine J. Wert, Gabriel Velez, Kanchustambham Vijayalakshmi, Vishnu Shankar, Jesse D. Sengillo, Richard N. Zare, Alexander G. Bassuk, Stephen H. Tsang, Vinit B. Mahajan

## Abstract

Neurodegenerative diseases are debilitating, incurable disorders caused by progressive neuronal cell death. Retinitis pigmentosa (RP) is a blinding neurodegenerative disease that results in retinal photoreceptor cell death and progresses to the loss of the entire neural retinal network. We previously found that proteomic analysis of the adjacent vitreous serves as way to indirectly biopsy the neural retina and identify changes in the retinal proteome. We therefore analyzed protein expression in liquid vitreous biopsies from autosomal recessive retinitis pigmentosa (arRP) patients with *PDE6A* mutations and arRP mice with *Pde6ɑ mutations.* Proteomic analysis of retina and vitreous samples identified molecular pathways affected at the onset of photoreceptor cell death. Based on affected molecular pathways, arRP mice were treated with a ketogenic diet or metabolites involved in fatty-acid synthesis, oxidative phosphorylation, and the tricarboxylic acid (TCA) cycle. Dietary supplementation of a single metabolite, ɑ-ketoglutarate, increased docosahexaeonic acid (DHA) levels, provided neuroprotection, and enhanced visual function in arRP mice. A ketogenic diet delayed photoreceptor cell loss, while vitamin B supplementation had a limited effect. Finally, desorption electrospray ionization mass spectrometry imaging (DESI-MSI) revealed restoration of key metabolites that correlated with our proteomic findings: pyrimidine and purine metabolism (uridine, dihydrouridine, and thymidine), glutamine and glutamate (glutamine/glutamate conversion), and succinic and aconitic acid (TCA cycle). This study demonstrates that replenishing TCA cycle metabolites via oral supplementation prolongs vision and provides a neuroprotective effect on the photoreceptor cells and inner retinal network.

**One Sentence Summary:** The study shows protein and metabolite pathways affected during neurodegeneration and that replenishing metabolites provides a neuroprotective effect on the retina.

## Introduction

Neurodegenerative diseases are debilitating, chronic disorders associated with neuronal cell loss in sensory, motor, or cognitive systems [1]. They are typically characterized by abnormal proteostasis, where the cell is unable to maintain regulation of protein folding, translation, and degradation pathways. These defects in proteostasis lead to processes of cell death such as apoptosis and disrupted signaling mechanisms within the cell from oxidative stress, glutamate-mediated cytotoxicity, neuroinflammation, mitochondrial dysfunction, and increased endoplasmic reticulum (ER) stress [2–7]. However, the early factors involved in neuronal cell death are still not understood. Retinitis pigmentosa (RP) consists of a group of inherited retinal neurodegenerative diseases with high genetic heterogeneity, which can be caused by mutations in more than 60 genes. RP affects approximately 1.5 million people worldwide, and approximately 10% of Americans carry a recessive mutant RP allele. Despite precise genetic characterization of RP patients, identification of causative mutations has had limited impact on therapy. To date, the only treatment for RP is supplementation with vitamin A (15,000 IU/day), but its clinical value is controversial, it does not cure disease, and vitamin A can have adverse effects for other retinal neurodegenerations such as Stargardt macular dystrophy [8]. There have been recent advances to treat RP patients using prosthetics or gene therapy approaches [9–12], but only end-stage RP or patients with the *RPE65* gene mutation, respectively, are currently eligible for such therapies. Thus, there is a critical need to understand the underlying mechanisms of photoreceptor death and provide a general therapy independent of specific gene mutations. Delaying photoreceptor cell degeneration, even without a complete cure, can significantly increase the quality of life for RP patients and prolong their ability to live independently.

There is accumulating evidence linking neurodegeneration to energy metabolism, particularly in age-related neurological disorders like Huntington’s, Alzheimer’s, and Parkinson’s disease [1]. Neuronal aging is often accompanied by metabolic and neurophysiologic changes that are associated with impaired function. Retinal photoreceptors are metabolically robust and require high rates of aerobic ATP synthesis. Defects in retinal cellular metabolism are associated with degenerative retinal diseases, such as rod dystrophy and RP. We have previously shown, using proteomics, that the representation of metabolic pathways and enzymes in the human retina varies by anatomic location (i.e. the peripheral, juxta-macular, and foveomacular regions) [13]. In addition to anatomical differences in metabolic pathway representation, energy consumption in photoreceptors is highly compartmentalized. It has been demonstrated that the mammalian inner retina metabolizes a majority (∼70%) of its glucose via the tricarboxylic acid (TCA) cycle and oxidative phosphorylation (OXPHOS). In contrast, the outer retina metabolizes ∼60% of its glucose through glycolysis [14, 15]. Thus, defects in aerobic metabolism could preferentially affect inner retinal function. Indeed, photoreceptor function is highly susceptible to glycolytic inhibition (via sodium iodoacetate injection) and mature ‘RP’ rats lacking a majority of their photoreceptors display an ∼50% reduction in glycolytic activity compared to normal rat retinas [16, 17]. In addition to meeting the high energy demands of photoreceptors, glucose metabolism plays other non-energetic roles in the retina [18, 19]. For example, glucose metabolism by the pentose phosphate pathway leads to the production of intracellular glutathione, which prevents neuronal cell death through redox inactivation of cytochrome *c*-mediated apoptosis pathways [20]. Glucose metabolism also feeds into fatty acid synthesis and lipid metabolism pathways, which are responsible for maintaining proper membrane lipid composition in neuronal cells [21]. Disruption of lipogenesis (by deleting fatty acid synthase) in the neural retina leads to abnormal synaptic structure and apoptosis, resulting in a phenotype resembling RP [22]. Thus, loss of these key metabolic pathways and enzymes would result in abnormal nutrient and metabolite availability, and impaired visual function.

It has been shown that the risk of neuronal cell death is constant, regardless of the underlying cause [23]. Therefore, pathways must exist that can be targeted to generally reduce or protect against neuronal cell death and these pathways could be valuable therapeutic targets. A potential therapeutic approach for RP is to reprogram metabolic pathways that are altered during rod dystrophy and RP progression. In one such approach, we found that knocking down sirtuin-6 (SIRT6) to reprogram retinal cells toward anabolism delayed photoreceptor cell degeneration in a mouse model of RP [24]. For this study, we applied a liquid biopsy proteomics approach to identify the molecular pathway targets affected during RP progression. Using proteomics, we have previously shown that molecular changes in the neural retina can be detected through proteomic analysis of human and mouse vitreous biopsies, since the retina deposits proteins into the adjacent vitreous extracellular matrix in retinal disease [25–29]. In this study, we collected vitreous biopsy samples from patients with autosomal recessive retinitis pigmentosa (arRP) carrying mutations in *PDE6A*, a gene encoding the catalytic ɑ-subunit of the rod-specific cGMP-dependent phosphodiesterase (PDE6), and from mice carrying a *Pde6ɑ* mutation. Based on proteomic analyses, we targeted diseased metabolic pathways to investigate treatments for RP patients and found key metabolites that prolong photoreceptor cell survival and visual function. We further compared the distribution of these metabolites using desorption electrospray ionization mass spectrometry imaging (DESI-MSI) between untreated *Pde6ɑ* mice and metabolite-treated mice.

## Methods

### Study approval

The study protocol was approved by the Institutional Review Board for Human Subjects Research (IRB) at Columbia University and Stanford University, was HIPAA compliant, and adhered to the tenets of the Declaration of Helsinki. All subjects underwent informed consent for study participation. All experiments were performed in accordance with the ARVO Statement for the Use of Animals in Ophthalmic and Visual Research and were all approved by the Animal Care and Use Committee at Columbia University and Stanford University.

### Mouse lines and husbandry

C57BL/6J-*Pde6α^nmf363/nmf363^*, with a D670G mutation, herein referred to as *Pde6α^D670G^* mice, were obtained from the Jackson Laboratory (Bar Harbor, ME, USA). *Pde6α^D670G^*are co-isogenic in the C57BL/6J (B6) background; therefore, age-matched B6 mice were used as experimental controls. Mice were bred and maintained at the facilities of Columbia University and Stanford University. Animals were kept on a light–dark cycle (12 hour–12 hour). Food and water were available *ad libitum* throughout the experiment, whether treated or untreated. The ketogenic diet was provided by BioServ (Ketogenic Diet, AIN-76A-Modified, High-Fat, Paste; Flemington, NJ) in place of standard chow. The ketogenic diet was stored refrigerated and changed every other day for the mice during the duration of the experiment. Vitamins B_2_ (riboflavin, Sigma Aldrich) and B_3_ (nicotinamide, Sigma Aldrich) were provided at a concentration of 2.5 g/L in the mouse drinking water. ɑ-KG (alpha-ketoglutaric acid, Sigma Aldrich) was provided at a concentration of 10 g/L in the mouse drinking water. Melatonin (Sigma Aldrich) was provided at a concentration of 80 mg/L in the mouse drinking water. Resveratrol (Sigma Aldrich) was provided at a concentration of 120 mg/L in the mouse drinking water. All water bottles were covered in foil to reduce light exposure and changed weekly during the duration of the experiment following approved institutional guidelines.

### Human patient imaging

Clinical examination and testing were performed as previously described [30]. Autofluorescence (AF) images were obtained using a Topcon TRC 50DX camera (Topcon, Pyramus, NJ, USA). Optical coherence tomography imaging was obtained from the spectral-domain Heidelberg HRA2 Spectralis, version 1.6.1 (Heidelberg Engineering, Inc, Vista, CA, USA). Genetic testing was performed as previously described [31].

### Mouse autofluorescence/infrared imaging

AF/IR fundus imaging was obtained with the Spectralis scanning laser confocal ophthalmoscope (OCT-SLO Spectralis 2; Heidelberg Engineering, Heidelberg, Germany) as previously described [32, 33]. Pupils were dilated using topical 2.5% phenylephrine hydrochloride and 1% tropicamide (Akorn, Inc., Lakeforest, IL, USA). Mice were anesthetized by intraperitoneal injection of 0.1 mL /10 g body weight of anesthesia [1 mL ketamine—100 mg/ mL (Ketaset III, Fort Dodge, IA, USA) and 0.1 mL xylazine—20 mg/ mL (Lloyd Laboratories, Shenandoah, IA, USA) in 8.9 mL PBS]. AF imaging was obtained at 488-nm absorption and 495-nm emission using a 55° lens. IR imaging was obtained at 790 nm absorption and 830 nm emission using a 55° lens. Images were taken of the central retina, with the optic nerve located in the center of the image.

### Electroretinography (ERG)

Mice were dark-adapted overnight, manipulations were conducted under dim red-light illumination, and recordings were made using Espion or Celeris ERG Diagnosys equipment (Diagnosys LLL, Littleton, MA, USA) as previously described [32, 33]. Briefly, adult B6 control mice were tested at the beginning of each session to ensure equal readouts from the electrodes for both eyes before testing the experimental mice. Pupils were dilated using topical 2.5% phenylephrine hydrochloride and 1% tropicamide (Akorn Inc., Lakeforest, IL, USA). Mice were anesthetized by intraperitoneal injection of 0.1 mL /10 g body weight of anesthesia [1 mL ketamine 100 mg/mL (Ketaset III, Fort Dodge, IA, USA) and 0.1 mL xylazine 20 mg/mL (Lloyd Laboratories, Shenandoah, IA, USA) in 8.9 mL PBS]. Body temperature was maintained at 37°C during the procedure. Both eyes were recorded simultaneously, and responses were averaged for each trial. Responses were taken from the Espion/Celeris readout in microvolts. Statistical significance was set at *p* < 0.05, and 1-way ANOVA followed by Tukey’s multiple comparison’s test was used to compare all treated, untreated, and wild-type groups.

### Histology

Mice were sacrificed, and the eyes enucleated as previously described [34–36]. Eyes were embedded in paraffin, sectioned, and stained with hematoxylin and eosin by Exaclibur Pathology, Inc. (Norman, OK), before being visualized by light microscopy (Leica DM 5000B, Leica Microsystems, Germany). Quantification of photoreceptor nuclei was conducted on several sections that contained the optic nerve, as follows: the distance between the optic nerve and the ciliary body was divided into three, approximately equal, quadrants on each side of the retina. Four columns of nuclei (how many cell nuclei thick) were counted within each single quadrant. These counts were then used to determine the average thickness of the ONL for each individual animal and within each quadrant of the retina, spanning from the ciliary body to the optic nerve head (ONH) and out to the ciliary body. Sectioning proceeded along the long axis of the segment, so that each section contained upper and lower retina as well as the posterior pole. Statistical analysis was performed on the average ONL thickness of each individual animal, taking into account all cell counts for all quadrants, and Student’s *t*-test was performed with statistical significance set at *p* < 0.05.

### Human vitreous sample collection

Pars plana vitrectomy for vitreous biopsy was performed as previously described [37]. We used a single-step transconjunctival 23-gauge trocar cannula system (Alcon Laboratories Inc, Fort Worth, TX). A light pipe and vitreous cutter were inserted into the mid vitreous and the cutter was activated for 30 seconds without infusion. An undiluted 1.0-cc sample of vitreous was then manually aspirated into a 3-cc syringe. Vitreous samples were immediately centrifuged in the operating room at 15,000 g for 5 minutes at room temperature to remove particulate matter, and samples were then stored at −80 °C.

### Mouse vitreous and retina sample collection

The vitreous and retina from 12 mouse eyes were eviscerated as described previously [26, 38, 39]. Briefly, scleral tissue posterior to the limbus was grasped with 0.22 forceps and a microsurgical blade was used to make a linear incision in the cornea from limbus to limbus. A fine curved needle holder was inserted behind the lens toward the posterior aspect of the globe. The needle holder was partially closed and pulled forward pushing the lens through the corneal incision while leaving the eye wall intact. The vitreous was partially adherent to the lens. The lens-vitreous tissue was then placed into a filtered centrifugation tube containing 20 microliters of protease inhibitor cocktail (Roche) dissolved in PBS. The fine curved needle holder was placed as far posterior to the globe as possible, near the optic nerve. The needle holder was partially closed, and pulled forward, pushing the retina forward through the corneal incision. The vitreous appeared as a translucent gel adherent to the retina. The retina-vitreous tissue was placed into the filtered centrifuge tube containing the lens-vitreous tissue. The filtered centrifuge tube was spun at 14,000 x g for 12 minutes and the eluent (vitreous) was collected. Retina was separated from the lens tissue remaining in the filtered centrifuge tube and collected in PBS containing protease inhibitor cocktail.

### Protein extraction and digestion

The received fluid samples were diluted in 2% SDS, 100 mM Tris-HCl (pH 7.6), 100 mM DTT to approximately 0.5 mL volume and heated at 95 °C for 10 min. Each sample was then briefly vortexed and sonicated for 10 seconds using a probe-tip sonicator (Omni International). The samples were then returned to incubate at 95 °C for an additional 10 min. Samples were then transferred to a 30 K Amicon MWCO device (Millipore) and centrifuged at 16,100 x g for 30 min. Then 400 µL of 8 M urea, 100 mM Tris-HCl (pH 7.6) was added to each device and centrifuged as before and the filtrate discarded. This step was repeated. Then 400 µL of 8M urea, 100 mM Tris-HCl (pH 7.6), 15 mM iodoacetamide was added to each device and incubated in the dark for 30 minutes. The samples were then centrifuged as before, and the filtrate discarded. Then 400 µL of 8 M urea, 100 mM Tris-HCl (pH 7.6) was added to each device and centrifuged as before and the filtrate discarded. This step was repeated. Then 400 µL of 2 M urea, 100 mM Tris-HCl (pH 7.6) was added to each device along with 2.5 µg trypsin. The devices incubated overnight on a heat block at 37 °C. The devices were then centrifuged, and the filtrate collected. Then 400 µL 0.5 M NaCl was added to each device and centrifuged as before. The filtrate was added to the previously collected filtrate.

### Peptide desalting and fractionation

Digested peptides were desalted using C18 stop-and-go extraction (STAGE) tips. Briefly, for each sample a C18 STAGE tip was activated with methanol, then conditioned with 75% acetonitrile, 0.5% acetic acid followed by 0.5% acetic acid. Samples were loaded onto the tips and desalted with 0.5% acetic acid. Peptides were eluted with 75% acetonitrile, 0.5% acetic acid and lyophilized in a SpeedVac (Thermo Savant) to dryness, approximately 2 hours. Peptides were fractionated using SAX STAGE tips. Briefly, for each sample a SAX STAGE tip was activated with methanol, then conditioned with Britton-Robinson buffer (BRB), pH 3.0 followed by BRB (pH 11.5). Peptides were loaded onto the tips and the flow-through collected followed by and five additional fractions by subsequent application of BRB at pH 8.0, 6.0, 5.0, 4.0 and 3.0. Each fraction was desalted using a C18 STAGE tip and lyophilized as described above.

### Liquid chromatography-tandem mass spectrometry (LC-MS/MS)

Each SAX fraction was analyzed by LC-MS/MS. LC was performed on an Agilent 1100 Nano-flow system. Mobile phase A was 94.5% MilliQ water, 5% acetonitrile, 0.5% acetic acid. Mobile phase B was 80% acetonitrile, 19.5% MilliQ water, 0.5% acetic acid. The 150 min LC gradient ran from 5% A to 35% B over 105 minutes, with the remaining time used for sample loading and column regeneration. Samples were loaded to a 2 cm x 100 µm I.D. trap-column positioned on an actuated valve (Rheodyne). The column was 13 cm x 100 µm I.D. fused silica with a pulled tip emitter. Both trap and analytical columns were packed with 3.5 µm C18 (Zorbax SB, Agilent). The LC was interfaced to a dual pressure linear ion trap mass spectrometer (LTQ Velos, Thermo Fisher) via nano-electrospray ionization. An electrospray voltage of 1.5 kV was applied to a pre-column tee. The mass spectrometer was programmed to acquire, by data-dependent acquisition, tandem mass spectra from the top 15 ions in the full scan from 400 - 1400 *m/z*. Dynamic exclusion was set to 30 s.

### LC-MS/MS data processing, library searching, and analysis

Mass spectrometer MS data files were converted to mzXML format and then to mgf format using msconvert in ProteoWizard and mzXML_to_mgf v4.4, rev 1, respectively. Peak list data were searched using two algorithms: OMSSA v2.1.9 and X!Tandem TPP v4.4, rev 1. The UniProt mouse protein sequence library was used in a target-decoy format (derived from UniProt as of 10/22/2012). The mgf files were searched using OMSSA with precursor settings of +/-2.0 Da and fragment tolerance of +/-0.8 Da. In X!Tandem, precursor settings were set to −1 Da and +3 Da and the fragment tolerance to 0.8 Da. The number of missed cleavages was set to 1; the fixed modifications were Carbamidomethyl (C); the variable modifications were Oxidation (M). XML output files were parsed using TPP v4.4, rev 1 in the programs PeptideProphet, iProphet, and ProteinProphet. Proteins were required to have 2 or more unique peptides with E-value scores of 0.01 or less. Relative quantitation was performed by spectral counting. Data were normalized based on total spectral counts (hits) per sample. Results were saved in Excel as .txt format and were uploaded into the Partek Genomics Suite 6.5 software package as described previously [25–29]. The data was normalized to log base 2 and compared using 1-way ANOVA analysis. All proteins with non-significant (p>0.05) changes were eliminated from the table. The significant values were mapped using the ‘cluster based on significant genes’ visualization function with the standardization option chosen. PANTHER Pathway Analysis was utilized to determine the most significant molecular pathways affected by the proteins present in each group [40]. Gene ontology (GO) analysis was also performed in PANTHER. Pie charts were created for the visualization of GO distributions within the list of proteins for each GO term category including biological process, molecular function, and cellular component.

### Desorption electrospray ionization mass spectrometry imaging (DESI-MSI)

Retinas were dissected and fresh frozen in optimal cutting temperature (OCT) media. 10-µm cryosections were collected (Cryostat Microsystem, Leica Biosystems) with at least three sections per SuperFrost Plus Slides (Fisher Scientific), with at least three slides collected per retina (at least 9 sections per retinal sample). Briefly, DESI-MSI was performed in the negative ion mode (−5 kV) from *m/z* 50–1000, using LTQ-Orbitrap XL mass spectrometer (Thermo Scientific) coupled to a home-built DESI-source and a two-dimensional (2D) motorized stage. The retinal tissues were raster scanned under impinging charged droplets using a 2D moving stage in horizontal rows separated by a 200-µm (spatial resolution) vertical step. The impinging primary charged droplets were generated from the electrospray nebulization of a histologically compatible solvent system, 1:1 (vol/vol) dimethylformamide/acetonitrile (DMF/ACN, flow rate 1 µL/min). The electrospray nebulization was assisted by sheath gas nitrogen (N_2_, 170 psi) and a high electric field of −5 kV. The spatial resolution of DESI-MSI is defined by the spray spot size of ∼200 µm. DESI-MSI of all tissue samples were carried out under identical experimental conditions, such as spray tip-to-surface distance ∼2 mm, spray incident angle of 55°, and spray-to-inlet distance ∼5 mm. MSI data acquisition was performed using XCalibur 2.2 software (Thermo Fisher Scientific Inc.). An additional freeware tool, MSConvert (ProteoWizard software, version 2.1x), was employed to convert the XCalibur 2.2 mass spectra files (.raw files) into .mzML format files; then imzMLConverter (version 1.3.0) was used to combine .mzML files into .imzML format file, readable by another software MSiReader (version v1.00). The 2D chemical maps of molecular ions were plotted using Biomap. To identify significantly differentially-expressed metabolites, XCalibur raw data files were converted to csv files for statistical analysis. Raw csv data was imported to the R language for further processing. To account for random effects influencing mass spectrometry data, the intensity of each metabolite was normalized by the total ion current for the corresponding pixel. Next, a nearest neighbor clustering method was used to collect the pixel intensities corresponding to the nearest molecular ion peak. To determine which metabolites could distinguish *Pde6α^D670G^* mice from ɑ-KG treated mice, we applied the SAM (Significance Analysis of Microarrays) technique using the samr package in R [41]. The metabolites were filtered according to a false discovery rate (FDR) cutoff of 5%. SAM identified 913 potentially significant metabolites (FDR < 0.05), of which 377 metabolites have a fold change greater or less than 1. Although SAM returned 377 metabolites with notable fold change, to help interpret the biological role of metabolites, we restricted our analysis to peaks for which tandem-MS and high mass resolution analyses were performed by using the LTQ-Orbitrap XL (Thermo Scientific).

### Metabolite/lipid identification

For lipid and metabolite identification, tandem-MS and high-mass resolution analyses were performed using the LTQ-Orbitrap XL (Thermo Scientific). The retinal tissue sections were dissolved in 500 μL of methanol: water (80:20) for metabolite extraction. The undissolved tissue was centrifuged down by 4,000 × g for 5 min. The resultant clear solution was sprayed into mass spectrometer for tandem-MS and high-mass resolution analyses, at a flow rate of 2 μL/min, sheath gas pressure (N_2_, 120 psi), and the spray voltage −5 kV. For the MS/MS studies, a normalized collision energy of 20–40% was applied. The metabolites and lipids were assigned based on high-mass resolution analysis, isotope distribution, and tandem-MS fragmentation patterns. In the case when several detectable fragmentation patterns were generated by isomeric parent ions, the most intense fragmentation peaks were used to assign the molecule identity listed in SI Appendix. The LipidMaps (www.lipidmaps.org/), MassBank (www.massbank.jp), Metlin (https://metlin.scripps.edu/), and the human metabolome database (HMDB) were used for the metabolite identification. The detected species were mostly deprotonated small metabolites related to the tricarboxylic acid (TCA) cycle, and deprotonated lipids including free fatty acids (FAs), fatty acid dimers, phosphatidic acids, and glycerophospholipids.

### Statistical analysis

Differences in experimental groups were determined by the Student’s t-test as appropriate, or by one-way ANOVA followed by Tukey’s post-hoc multiple comparison’s test (see individual methods section). p values < 0.05 were considered significant. Measurements were done blinded to experimental group. All error bars represent standard deviation unless otherwise listed.

## Results

### Proteomics of the human vitreous highlight changes occurring in the neural retina during RP disease progression

The proband presented to us at age 38 with end-stage arRP. Genetic testing revealed p.R102C/p.S303C mutations in the *PDE6A* gene (II:5; Fig. 1A). Mutations in *PDE6* contribute to a significant fraction of RP cases (7-9%; OMIM: 180071) [42]. In comparison to a normal control (Fig. 1B-D), fundus autofluorescence (AF) revealed high-density central AF rings (Fig. 1E-F) and optical coherence tomography (OCT) revealed thinning of the neuronal cell layers (Fig. 1G). Additionally, the proband’s affected brother (II:4; Fig. 1A) displayed a similar disease stage with hyper-autofluorescent rings on AF imaging (Fig. 1H-I). OCT imaging similarly revealed thinning of retinal cell layers and cystoid macular edema (Fig. 1J). The affected brothers both developed significant epiretinal membranes (ERMs) and underwent vitrectomy surgery. Liquid vitreous biopsies were collected at the time of surgery (Fig. 2A-B, **Table S1**). Epiretinal membrane vitreous samples from two patients without RP were used as comparative controls since the two arRP patients underwent vitrectomy surgery for ERM removal (Fig. 2A-B). Vitreous samples were analyzed using shotgun liquid chromatography-tandem mass spectrometry (LC-MS/MS) to determine proteomic content (**Table S2**).

**Fig. 1.**
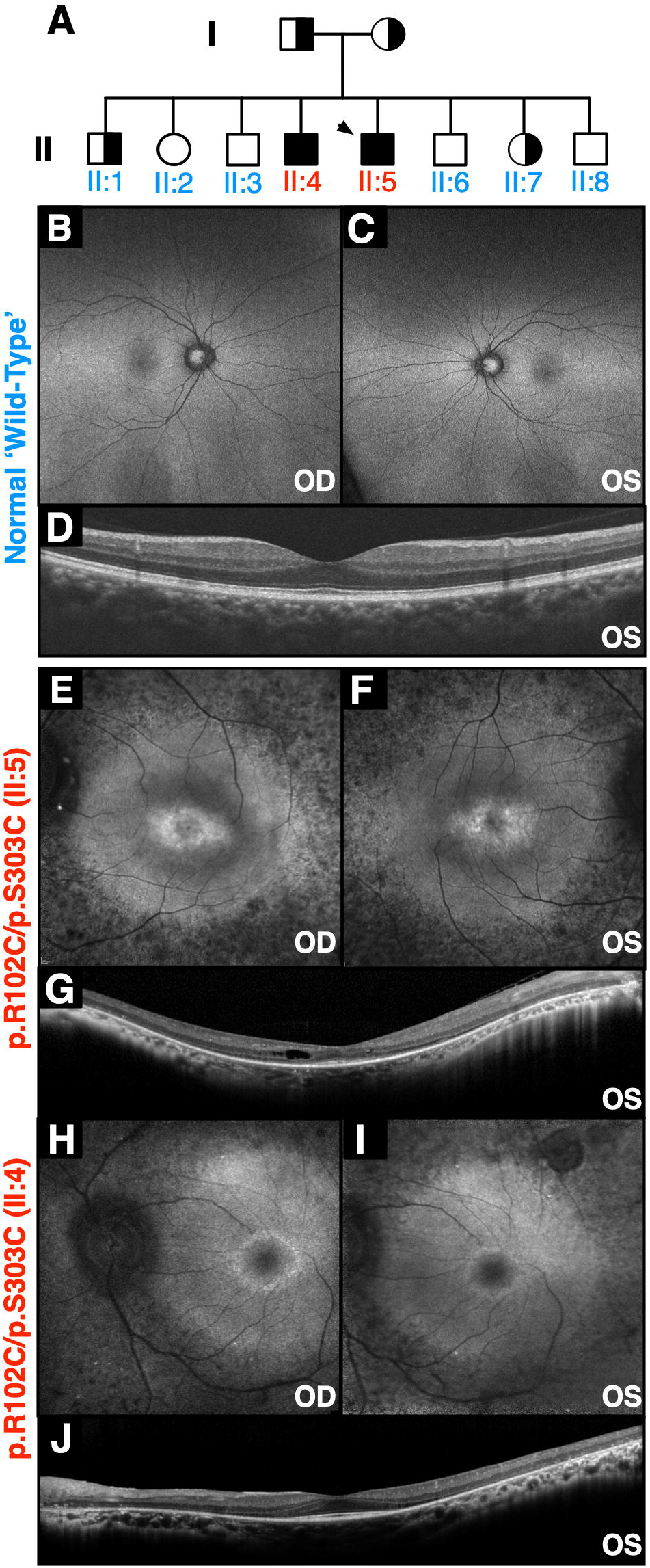
Two patients with autosomal recessive retinitis pigmentosa carrying mutations in *PDE6A*: (**A**) Pedigree of a family with autosomal recessive retinitis pigmentosa (arRP) caused by mutations in the PDE6A gene. The proband is denoted by the arrow. (**B-C**) Fundus autofluorescence (488nm) of a control human eye. (**D**) Spatially corresponding SD-OCT scans through the fovea of a control human eye. (**E-F**) Fundus autofluorescence (488nm) of the proband (II:5) revealed confluent areas of hypo-autofluorescence, suggesting RPE loss throughout the periphery, and a central ring of hyper-autofluorescence overlying the parafovea. (**G**) Spatially corresponding SD-OCT scans through the fovea confirmed marked peripheral outer retinal thinning and RPE loss. Despite macular edema, the fovea was relatively preserved, with a distinguishable ellipsoid zone. (**H-I**) Fundus autofluorescence of the affected brother (II:4) revealed similar confluent areas of hypo-autofluorescence and a central ring of hyper-autofluorescence overlying the parafovea. (**J**) Spatially corresponding SD-OCT scans showed similar marked peripheral outer retinal thinning and RPE loss.

**Fig. 2.**
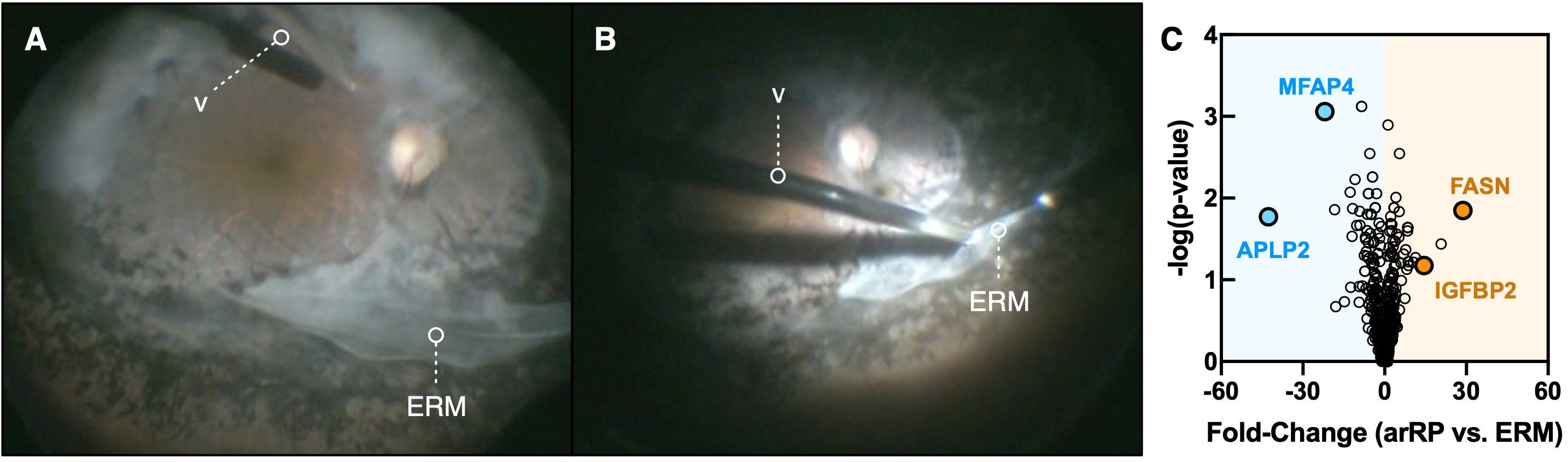
Proteomic analysis of *PDE6A* human vitreous. (**A-B**) Pars plana vitrectomy and epiretinal membrane removal procedure for an *PDE6A* (arRP) patient (II:5 in Fig. 1). A vitreous biopsy was collected from this patient and the affected brother. Protein content was analyzed by LC-MS/MS. The vitreous cutter is denoted by the letter ‘v.’ (**C**) Protein spectral counts were analyzed by 1-way ANOVA. Results are represented as a volcano plot. The horizontal axis (x-axis) displays the log2 fold-change value (arRP vs. ERM) and the vertical axis (y-axis) displays the noise-adjusted signal as the -log10 (p-value) from the 1-way ANOVA analysis. Increased levels of FASN and IGFBP2 are denoted by the orange circles. Blue circles indicate decreased levels of MFAP4 and APLP2 in arRP vitreous compared to controls.

It is rare to obtain arRP vitreous samples as the indication for vitrectomy surgery in these patients is exceptional. Despite the limitation of small sample size, descriptive semi-quantitative analysis can provide insight into underlying human disease mechanisms in a rare patient population, and there are no human arRP proteomic reports to date. We therefore performed label-free relative quantification of proteins identified in arRP vitreous relative to ERM, which was defined as the reference group. We detected 612 proteins in arRP vitreous by LC-MS/MS (**Table S3**). We compared protein expression between arRP and ERM vitreous: there were 110 elevated proteins (with a fold-change **≥** 2) in arRP vitreous compared to ERM controls (Fig. 2C). We observed a significant fraction (36%) of elevated, intracellular proteins in arRP vitreous despite the extracellular location and acellular composition of the vitreous. We previously reported that a significant fraction of proteins in the normal human vitreous are intracellular, particularly in the vitreous core, suggesting that surrounding tissues release proteins into the adjacent vitreous [27]. Among these elevated intracellular proteins were fatty acid synthase (FASN) and insulin-like growth factor-binding protein 2 (IGFBP2). FASN is involved in the synthesis of long-chain fatty acids from acetyl-CoA and NADPH. IGFBP2 has been shown to promote fibroblast differentiation and angiogenesis (Fig. 2C) [43, 44]. There were 135 depleted proteins (with a fold-change **≤** −2) in arRP vitreous compared to ERM controls. Among these proteins were microfibrillar-associated protein 4 (MFAP4) and amyloid beta precursor like 2 (APLP2; Fig. 2C). MFAP4 plays a role in maintaining integrin activation, which is important for maintaining RPE-cell adhesion to Bruch’s membrane [45]. Interestingly, APLP2 loss or dysfunction has been shown to result in abnormal amacrine and horizontal cell differentiation [45]. These results suggested that the observed proteomic changes in human arRP vitreous could reflect molecular changes in the neural retina. However, the human patient proteomes reflect samples collected at a very late stage of disease, long after significant photoreceptor death and may not reflect the critical cellular pathways altered at the onset of neurodegeneration. We therefore chose to gain further insight into the earlier disease stages using a preclinical model of arRP.

### Proteomic analysis of Pde6ɑ^D670G^ retina and vitreous identifies alterations in global protein expression that may precede cell loss and the RP clinical phenotype

We utilized the *Pde6ɑ^D670G^*mutant mouse to determine whether any of the cellular pathways found in our human patient proteomes were disrupted at the onset of rod dystrophy, before the clinical RP phenotype is detectable. The *Pde6ɑ^D670G^* phenotype closely resembles that of patients with arRP. These mice develop early onset severe retinal degeneration, characterized by photoreceptor death, retinal vessel attenuation, pigmented patches, ERG abnormalities, and white retinal spotting [33]. The mice display a loss of rod photoreceptor visual function by ERG analysis as early as one month of age and full loss of global visual function on ERG analysis by two months of age (Fig. 3A). In particular, *Pde6ɑ^D670G^*mice do not display histologically-detectable photoreceptor degeneration until after post-natal day 14 (P14; Fig. 3B), so retina and vitreous samples were collected from these mice at disease onset (P15), mid-stage (P30) and late-stage disease (P90; **Table S4**). Retina and vitreous samples for wild-type and *Pde6ɑ^D670G^*mice were trypsinized and underwent LC-MS/MS analysis (**Table S5-6**). Notably there were fewer unique proteins in the *Pde6ɑ^D670G^* mouse retina at 90 days (484 ± 63 individual proteins) than 15 days (1,135 ± 55 individual proteins), suggesting a decrease in protein expression as neurodegeneration progresses (**Table S7**). Conversely, we noticed there were more unique proteins in the *Pde6ɑ^D670G^* mouse vitreous at 15 days (364 ± 30 individual proteins) compared to wild-type vitreous (202 ± 9 individual proteins), suggesting an increase in vitreous protein expression at early stages of neuronal cell death (**Table S8**).

**Fig. 3.**
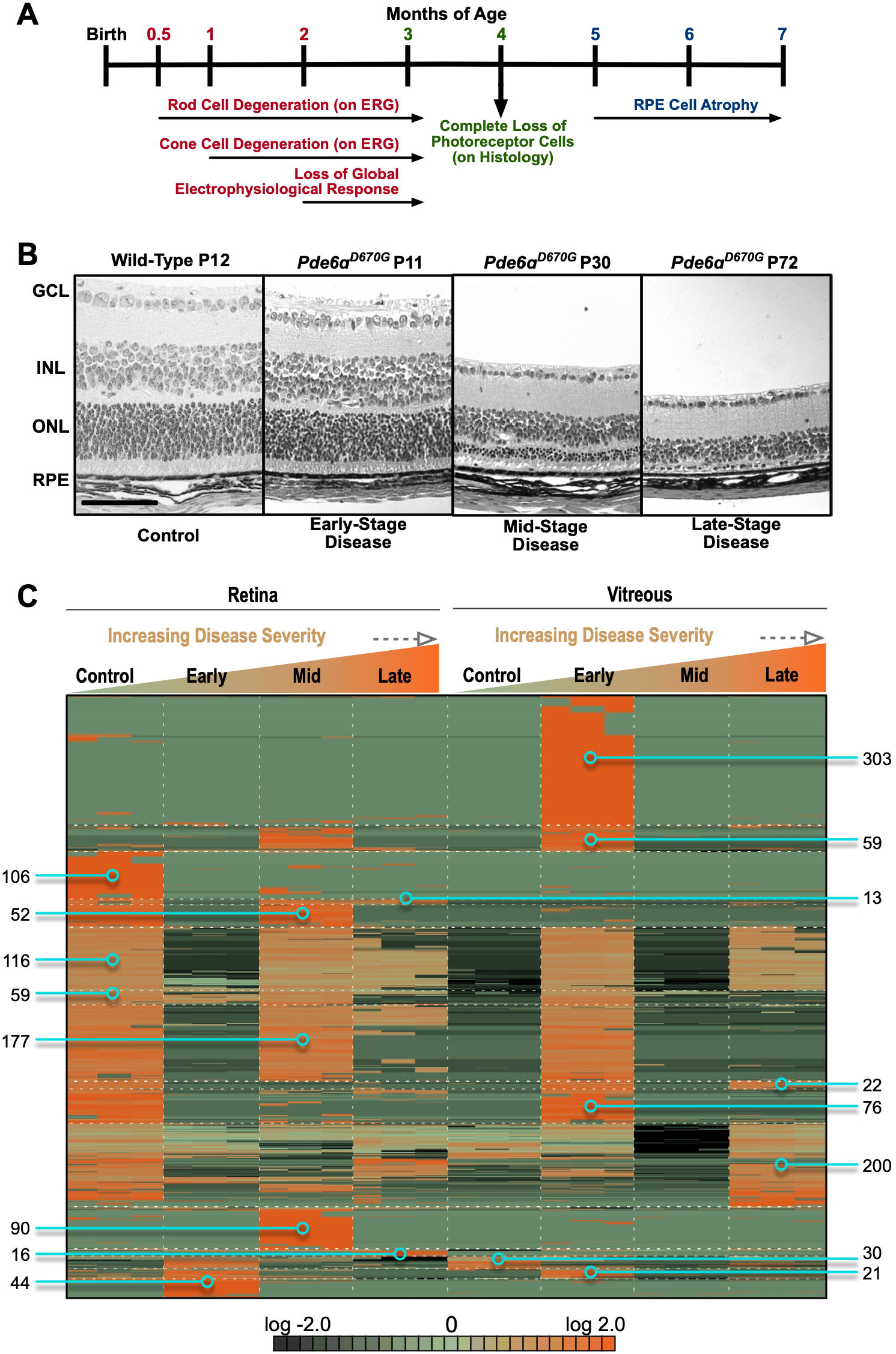
Proteomic analysis of *Pde6ɑ^D670G^*retina and vitreous at early-, mid-, and late-stage retinitis pigmentosa (RP). (**A**) Timeline of the clinical disease presentation in *Pde6ɑ^D670G^* mice. (**B**) Histological analysis of *Pde6ɑ^D670G^*mouse retinas at early-(P11), mid-(P30) and late-(P72) stage disease compared to a wild-type mouse at (P12) shows loss of the ONL photoreceptor cells over time. RPE, retinal pigment epithelium; ONL, outer nuclear layer; INL, inner nuclear layer; GCL, ganglion cell layer. Scale bar = 100 µm. (**C**) Protein spectral counts were analyzed 1-way ANOVA followed by hierarchical clustering. A total of 1,384 proteins were differentially-expressed in the *Pde6ɑ^D670G^* vitreous and retina samples compared to controls (p<0.05). Results are represented as a heatmap and display protein expression levels on a logarithmic scale. Orange indicates high expression while dark green/black indicates low or no expression.

Retina and vitreous protein levels (with two or more spectra) from *Pde6ɑ^D670G^*and wild-type mice were compared using 1-way ANOVA and heatmap clustering (**Table S9**). A total of 1,384 proteins were differentially-expressed in the *Pde6ɑ^D670G^* retina and vitreous samples compared to controls (p<0.05; Fig. 3C). There were 303 proteins highly-expressed in *Pde6ɑ^D670G^* vitreous at day 15 that were not present in the control retina or vitreous (Fig. 3C), suggesting that some proteins may have migrated into the vitreous from outside the eye. A majority of these proteins were intracellular and represented protein synthesis, gene expression, DNA replication, recombination, and repair pathways. Of these 303 proteins, there were 16 downstream effectors of the CD3 receptor: Coronin-1A (Coro1a), DEAD box protein 19A (Ddx19a), eukaryotic translation initiation factor 4E (Eif4e), GABA receptor-associated protein-like 2 (Gabarapl2), hemopexin (Hpx), E3 ubiquitin-protein ligase Huwe1 (Huwe1), Lon protease homolog (Lonp1), acyl-protein thioesterase 1 (Lypla1), metastasis-associated protein (Mta2), alpha-N-acetylgalactosaminidase (Naga), purine nucleoside phosphorylase (Pnp), RNA polymerase II subunit B1 (Polr2a), ubiquitin receptor (Rad23b), thiosulfate sulfurtransferase (Tsta2), thioredoxin reductase 1 (Txnrd1), and alanine-tRNA ligase (Aars). The CD3 co-receptor is expressed on T-cells and activates cytotoxic and helper T-cells [46]. The presence of downstream CD3 effectors in the *Pde6ɑ^D670G^*vitreous suggests that lymphocytes may be migrating into the eye at early stages of the disease. There were 360 proteins expressed in the control retina that were found to be elevated in the *Pde6ɑ^D670G^* vitreous at P15 (Fig. 3C). These proteins were not present in control vitreous, indicating that they had migrated from the degenerating retina into the vitreous (Fig. 3C).

To obtain a global view of the *Pde6ɑ^D670G^* retina and vitreous proteomes at different stages of degeneration, a gene ontology (GO) analysis of highly represented proteins in each group was performed (statistically-significantly elevated proteins in each group; p < 0.05). When comparing the four total retinal protein profiles, the GO summaries for the control retina were similar to the *Pde6ɑ^D670G^*retina at P30 and P90. The highest represented categories were cellular process, catalytic activity, and intracellular regions (**Fig. S1**). Interestingly, there was a decrease in the number of proteins with transporter activity in the *Pde6ɑ^D670G^*retina at early stage neuronal cell death, consistent with a loss of protein expression at this stage (**Fig. S1**). When comparing the four total vitreous protein profiles, the GO summaries were distinct. Vitreous proteins differed in the molecular function and cellular compartment representation at each disease stage. For example, there was an increase in the number of membrane-bound proteins and macromolecular complexes detected in the *Pde6ɑ^D670G^* vitreous at early stage neurodegeneration when compared to control vitreous (**Fig. S2**). Together, this indicated that the arRP retina and vitreous express distinct functional categories of proteins compared to wild-type mice at each stage of disease progression, and that these protein pathways may reflect early diagnostic markers or targets for therapeutics preventing further neuronal cell death and loss of the neural retinal network.

### Oxidative metabolic pathways are downregulated in the Pde6ɑ^D670G^ retina at the onset of neuronal cell death

We hypothesized that cellular pathways disrupted at the onset of rod dystrophy, before significant histological loss of the photoreceptor cells is detectable, might reflect key targets for therapeutic approaches. Therefore, we focused our further investigation on the *Pde6ɑ^D670G^* mouse proteomes collected at early-stage RP disease (P15). To examine differential protein expression at the onset of neuronal cell death, vitreous and retina protein levels (with two or more spectra) from *Pde6ɑ^D670G^* (P15) and wild-type mice were compared using 1-way ANOVA and hierarchical heatmap clustering. A total of 1,067 proteins were differentially-expressed in the *Pde6ɑ^D670G^* samples compared to controls (p<0.05; Fig. 4A**; Table S10**). Similar to the previous analysis comparing all *Pde6ɑ^D670G^* stages (Fig. 3C), there was a significant decrease in protein expression in *Pde6ɑ^D670G^*retina at early-stage neurodegeneration (P15). There was also a significant increase in vitreous protein expression at the onset of neuronal cell death in the *Pde6ɑ^D670G^*mice (779 upregulated proteins; p<0.05). The decrease in retinal protein expression and corresponding increase in vitreous protein expression suggested that the degenerating neural retina may leak proteins into the vitreous, similar to that seen in human patient samples [28, 47]. We then compared the list of proteins highly-abundant in control mouse retinas to that of the elevated proteins in *Pde6ɑ^D670G^* vitreous at the onset of neuronal cell death (P15). There were 446 proteins (when comparing protein expression profiles at P15 only) that were expressed in the control retina that were found to be elevated in the P15 *Pde6ɑ^D670G^* vitreous (Fig. 4A**; Table S11**). These proteins were not present in control vitreous. GO analysis categorized a significant fraction of these proteins to be intracellular, suggesting that they have migrated from the degenerating neural retina into the extracellular vitreous (**Fig. S3A**). The proteins with the most significant fold-change (FC) were ADP/ATP translocase 1 (Slc25a4; p-value = 1.68e-15), malate dehydrogenase 2 (Mdh2; p-value = 1.00e-19), and ATP synthase (Atp5a1; p-value = 2.06e-13). Categorization of these 446 proteins by their biological process revealed that a significant fraction are involved in energetic processes (**Fig. S3B**).

**Fig. 4.**
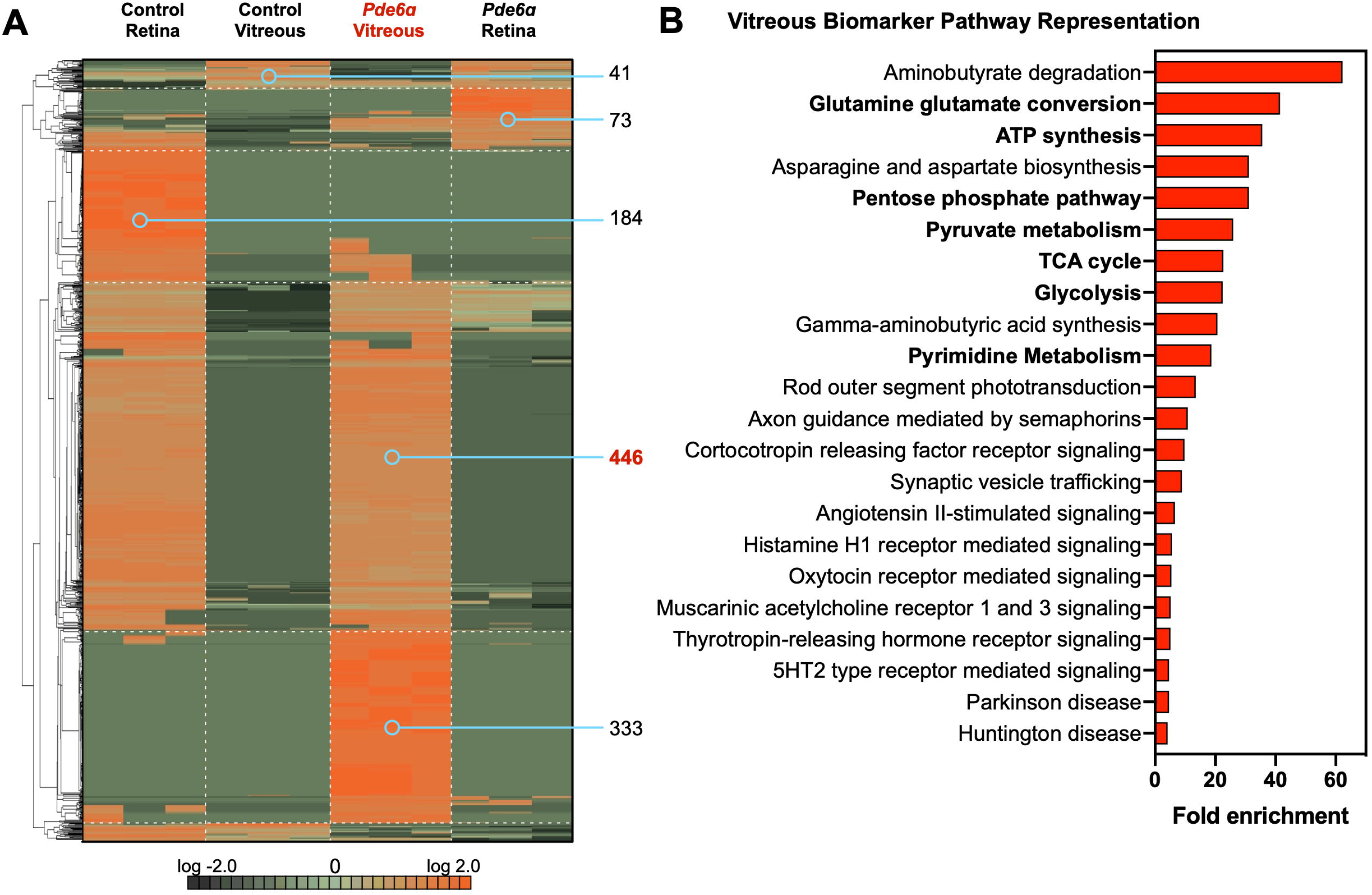
Proteomic analysis of *Pde6ɑ^D670G^*retina and vitreous identifies global protein expression changes. (**A**) Protein spectral counts were analyzed 1-way ANOVA followed by hierarchical clustering. A total of 1,067 proteins were differentially-expressed in the *Pde6ɑ^D670G^* vitreous and retina samples (when comparing protein expression profiles at P15 only) compared to controls (p<0.05). Results are represented as a heatmap and display protein expression levels on a logarithmic scale. Orange indicates high expression while dark green/black indicates low or no expression. There were 446 proteins that were downregulated in the *Pde6ɑ^D670G^* retina that were upregulated in the *Pde6ɑ^D670G^*vitreous. (**B**) Pathway representation of the 446 candidate biomarkers in the *Pde6ɑ^D670G^*vitreous. Pathways are ranked by their fold-enrichment obtained from the Mann-Whitney U Test. Metabolic pathways are bold-faced. Energetic pathways are denoted by bold-faced text.

We next analyzed the pathway representation of these downregulated retinal proteins. Pathway analysis revealed that these proteins represent metabolic (e.g. OXPHOS, TCA cycle), synaptic signaling, and visual transduction pathways (Fig. 4B**; Tables S12-15**). As anticipated, we found key proteins involved in the rod photoreceptor transduction pathway to be increased in the arRP vitreous at the onset of rod cell degeneration (P15): rhodopsin (Rho), guanine nucleotide-binding protein alpha and beta subunits (Gnat1 and Gnb1), phosducin (Pdc), cyclic nucleotide-gated channel beta 1 (Cngb1), rhodopsin kinase (Grk1), recoverin (Rcvrn), and S-arrestin (Sag). Taken together, these results suggest that these 446 proteins are candidate biomarkers for early arRP detection and possible targets for neuroprotective therapeutics. We identified proteins involved in glycolysis, TCA cycle, and OXPHOS: hexokinase (Hk2), aldolase A, (Aldoa), phosphofructokinase (Pfkl), enolase (Eno1), pyruvate kinase (Pkm), pyruvate dehydrogenase E1 component subunit (Pdha1), citrate synthase (Cs), aconitate hydratase (Aco2), isocitrate dehydrogenase (Idh1 and −3), ɑ-ketoglutarate dehydrogenase (Ogdh), succinate-CoA ligase beta subunit (Sucla2), succinate dehydrogenase flavoprotein subunit (Sdha), and malate dehydrogenase (Mdh1 and −2). We also identified proteins involved in fatty acid and amino acid synthesis pathways: fatty acid synthase (Fasn) and glutamic-oxaloacetic transaminase (Got1 and Got2). This observation suggested that retinal oxidative metabolism was affected in early retinal degeneration. Defects in oxidative metabolism have been previously shown to contribute to photoreceptor degeneration in RP patients [48]. For example, loss-of-function mutations in the *IDH3B* gene encoding the beta subunit of NAD-specific isocitrate dehydrogenase cause non-syndromic RP [49]. We similarly noted increased levels of fatty acid synthase in the vitreous of *Pde6ɑ^D670G^*mice at the onset of photoreceptor degeneration. Loss of FASN in the neural retina has been shown to result in progressive neurodegeneration resembling RP in mouse models [22]. Additionally, we have previously shown that metabolic and antioxidant stress proteins were regionally expressed within the human retina, in particular the OXPHOS and TCA cycle pathways, indicating that the retinal neural network is likely sensitive to changes within the metabolic pathways that occur during disease [13]. Although our human case represented a late-stage of disease, after loss of the rod neuronal cells, the validation and identification of a candidate biomarker with our early-stage *Pde6ɑ^D670G^* mouse proteome suggested potential target pathways for therapeutic approaches (Fig. 5).

**Fig. 5.**
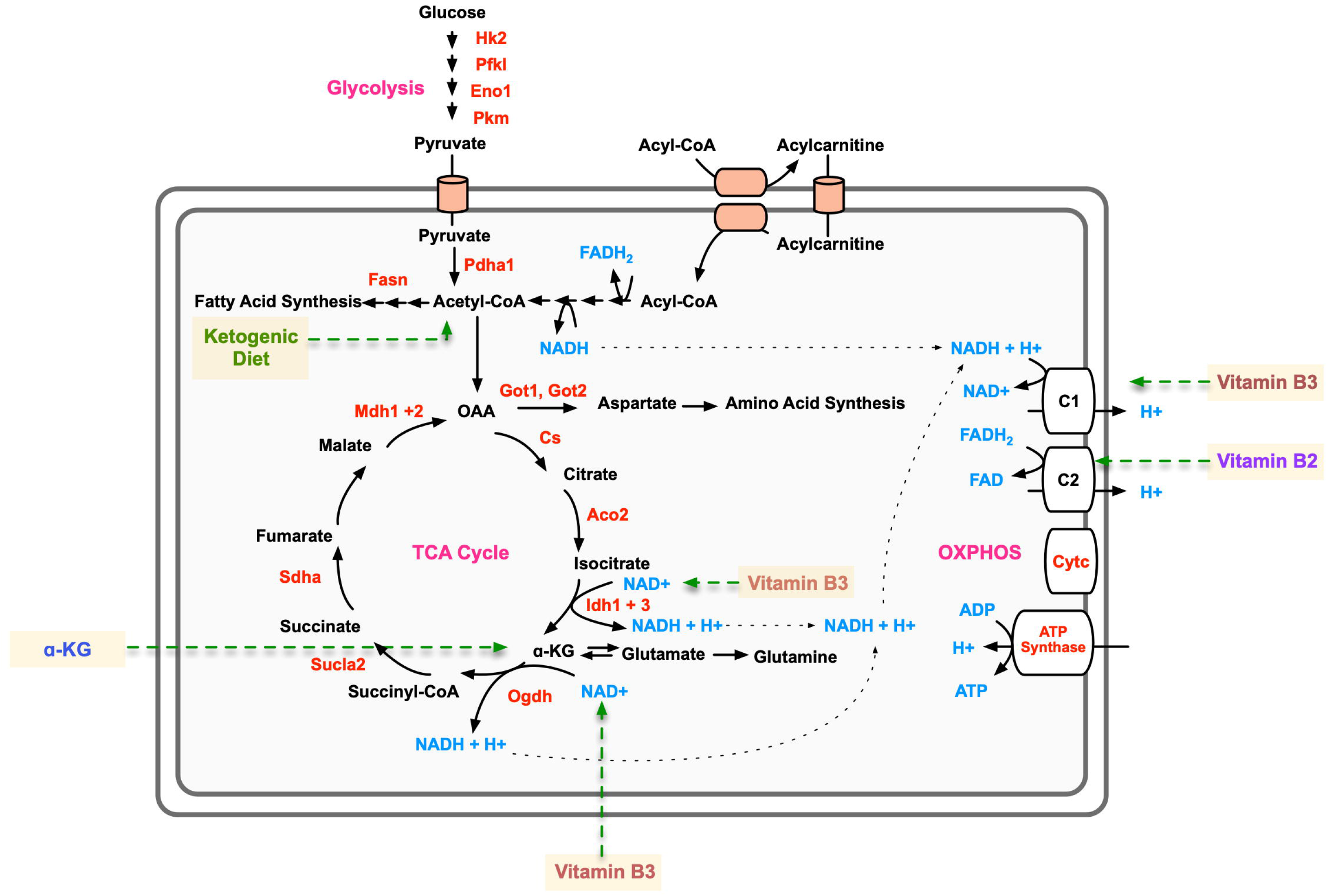
Proteomics-guided metabolite supplementation strategy for arRP. Metabolic pathway diagram highlighting identified significantly-elevated metabolic proteins in the *Pde6ɑ^D670G^* vitreous (red text). Metabolites are represented by blue text. Affected pathways include the TCA cycle, glycolysis, and oxidative phosphorylation (OXPHOS; magenta text). The metabolite/dietary therapies are highlighted in yellow.

### The ketogenic diet delays photoreceptor cell loss in arRP mice

We tested the arRP candidate biomarker, FASN, by targeting this critical protein with a dietary approach. The ketogenic diet has gained public popularity for health and weight loss benefits, as well as being an effective treatment option for young patients with epileptic seizures that do not respond to standard drug treatments [50]. The ketogenic diet provides acetyl-CoA that can feed into both the TCA cycle and the fatty acid synthesis pathways (Fig. 5). We placed arRP mice on a ketogenic diet (6:1:1 fat:protein:carbohydrate) beginning at post-natal day 21 (P21), mid-stage of photoreceptor degeneration. Histological analysis of the outer nuclear layer (ONL) thickness showed mild, but significant, photoreceptor cell survival in mice treated with the ketogenic diet compared to untreated control littermates at one month of age (Fig. 6A). We next examined visual function at one month of age. Representative ERG traces of arRP mice fed the ketogenic diet showed no difference in the rod photoreceptor cell response, but there was a mild rescue of the global visual response compared to littermates on standard chow diet (Fig. 6B). Quantification of ERG recordings from arRP mice treated with the ketogenic diet showed no significant difference in ERG responses compared to littermates on a standard chow diet, although there was a trend toward a- and b-wave visual rescue at the maximal scotopic 1.0 ERG setting (Fig. 6C-E). One limitation of the ketogenic diet was that the diet could not support nursing pups during early development, so it was not initiated until P21, at mid-stage of retinal disease when a rescue effect may be diminished. Therefore, we considered oral supplementation of metabolites that could be initiated earlier.

**Fig. 6.**
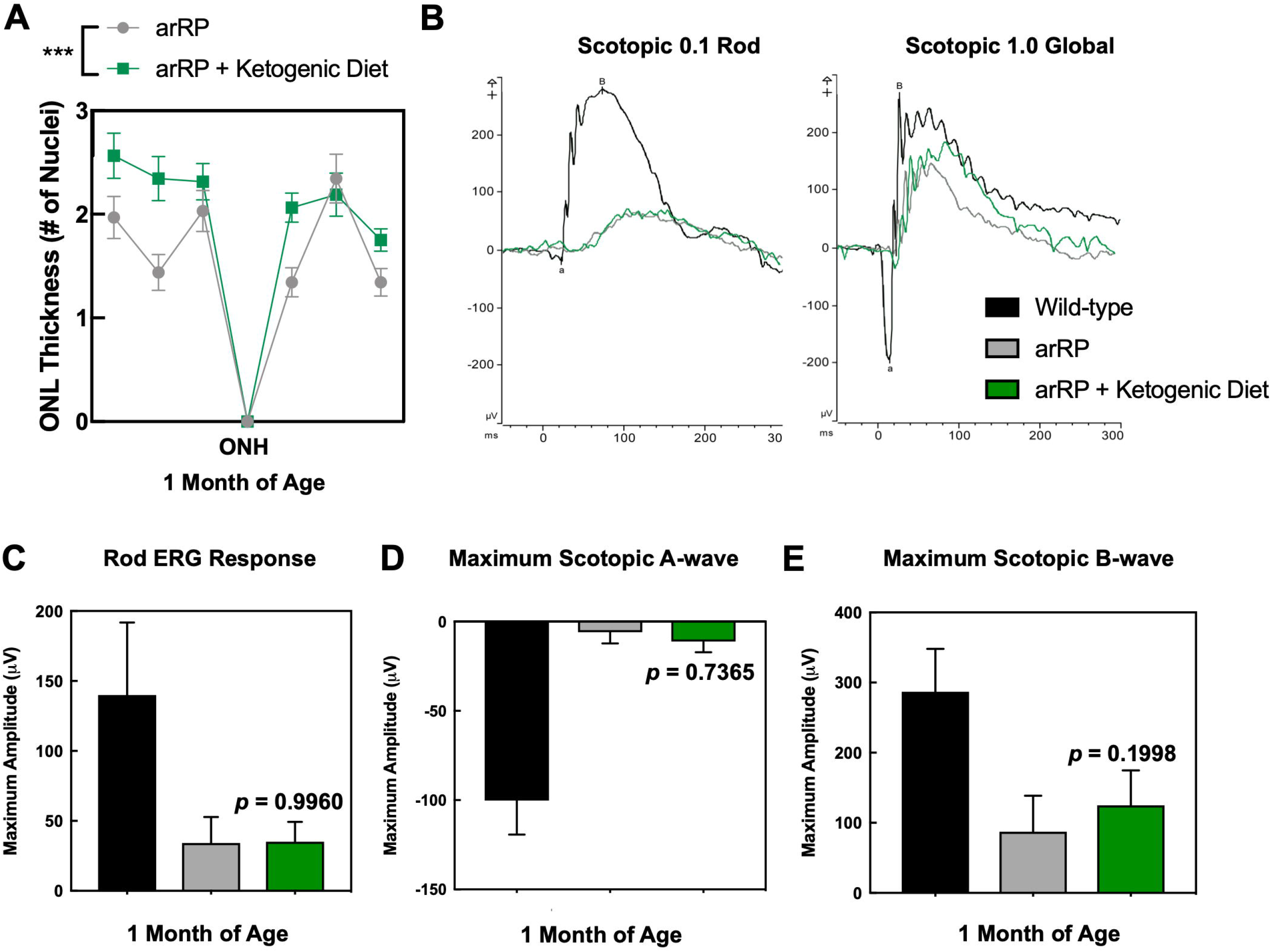
Treatment with the ketogenic diet rescues photoreceptor cell survival in the arRP preclinical mouse. (**A**) Histological analysis of the retinas of *Pde6ɑ^D670G^* mice on normal chow diet (gray) compared to *Pde6ɑ^D670G^*mice provided the ketogenic diet (green) shows a significant increase in the thickness of the outer nuclear layer (ONL) by number of photoreceptor cell nuclei in mice treated with the ketogenic diet at one month of age, one-week post-dietary treatment. Results are displayed as a morphometric quantification (spider graph) of ONL layer thickness superior (left) and inferior (right) to the optic nerve head (ONH). ***, *p* < 0.001. N = 8 eyes each group, with multiple ONL thickness counts per eye as described in Methods. Error bars = SEM. (**B**) Representative traces of the scotopic 0.1 dim-light rod-specific electroretinography (ERG; left) shows no significant difference in the rod photoreceptor cell response in *Pde6ɑ^D670G^* mice treated with the ketogenic diet (green) in comparison to untreated *Pde6ɑ^D670G^*mice (gray) at one month of age. Representative traces of the scotopic 1.0 global electroretinography (ERG; left) shows some rescue of the global visual response in *Pde6ɑ^D670G^*mice treated with the ketogenic diet (green) in comparison to wild-type controls (black) and untreated *Pde6ɑ^D670G^*mice (gray) at one month of age. Quantification of ERGs from a cohort of mice shows no significant visual rescue of (**C**) the rod photoreceptor cell response, (**D**) the maximum scotopic a-wave response, or (**E**) the maximum scotopic b-wave response in mice treated with the ketogenic diet (green) in comparison with wild-type controls (black) or untreated *Pde6ɑ^D670G^* mice (gray) at one month of age, although there was a trend toward an increased a- and b-wave maximal response. N = 5 WT, 6 arRP, 6 arRP + ketogenic diet mice.

### Oral supplementation with single metabolites prolongs neuronal cell survival and vision

The onset of retinal cell degeneration occurs rapidly in the arRP preclinical model, before weaning age; therefore, metabolites were chosen that were known to transfer into the mother’s milk and cross tissue barriers to be fed to the nursing pups [51, 52]. Thus, *Pde6ɑ^D670G^* mice were provided with ɑ-ketoglutarate (ɑ-KG), vitamin B_3_, vitamin B_2_, melatonin, or resveratrol in the drinking water beginning at post-natal day 0 (P0) and examined for functional visual rescue by ERG at one-month post-treatment (Fig. 5). Representative ERG traces showed visual rescue both in the rod photoreceptors, rod-cone photoreceptor a-wave response, and global inner retina scotopic ERG response in *Pde6ɑ^D670G^* mice treated with ɑ-KG (Fig. 7). For vitamin B_3_ and vitamin B_2_, some mice showed enhanced ERG visual responses compared to controls, but analysis of a larger cohort of mice did not show statistical significance for photoreceptor cell and inner retina function on ERG (Fig. 7B-D). Both melatonin and resveratrol did not show an effect on visual function after supplementation in the *Pde6ɑ^D670G^* arRP mice (Fig. 7B-D). Since ɑ-KG oral supplementation prolonged visual function in the *Pde6ɑ^D670G^* mice, we examined the retina by histology (Fig. 7E-F). Histological analysis confirmed a statistically significant rescue of photoreceptor cells (ONL) and their inner/outer segments after treatment with ɑ-KG (Fig. 7E-F). Thus, oral supplementation with a single metabolite, ɑ-KG, provided a significant neuroprotective effect on the rod photoreceptors and neural retinal network for at least one month of age, when treated before disease onset. Overall, the rescue effects on photoreceptor cell survival and visual function after treatment with the ketogenic diet or ɑ-KG supported our human and mouse proteomic data, suggesting that the neural retina is sensitive to changes in metabolism, and that targeting the TCA cycle may be a potential therapeutic approach for RP patients.

**Fig. 7.**
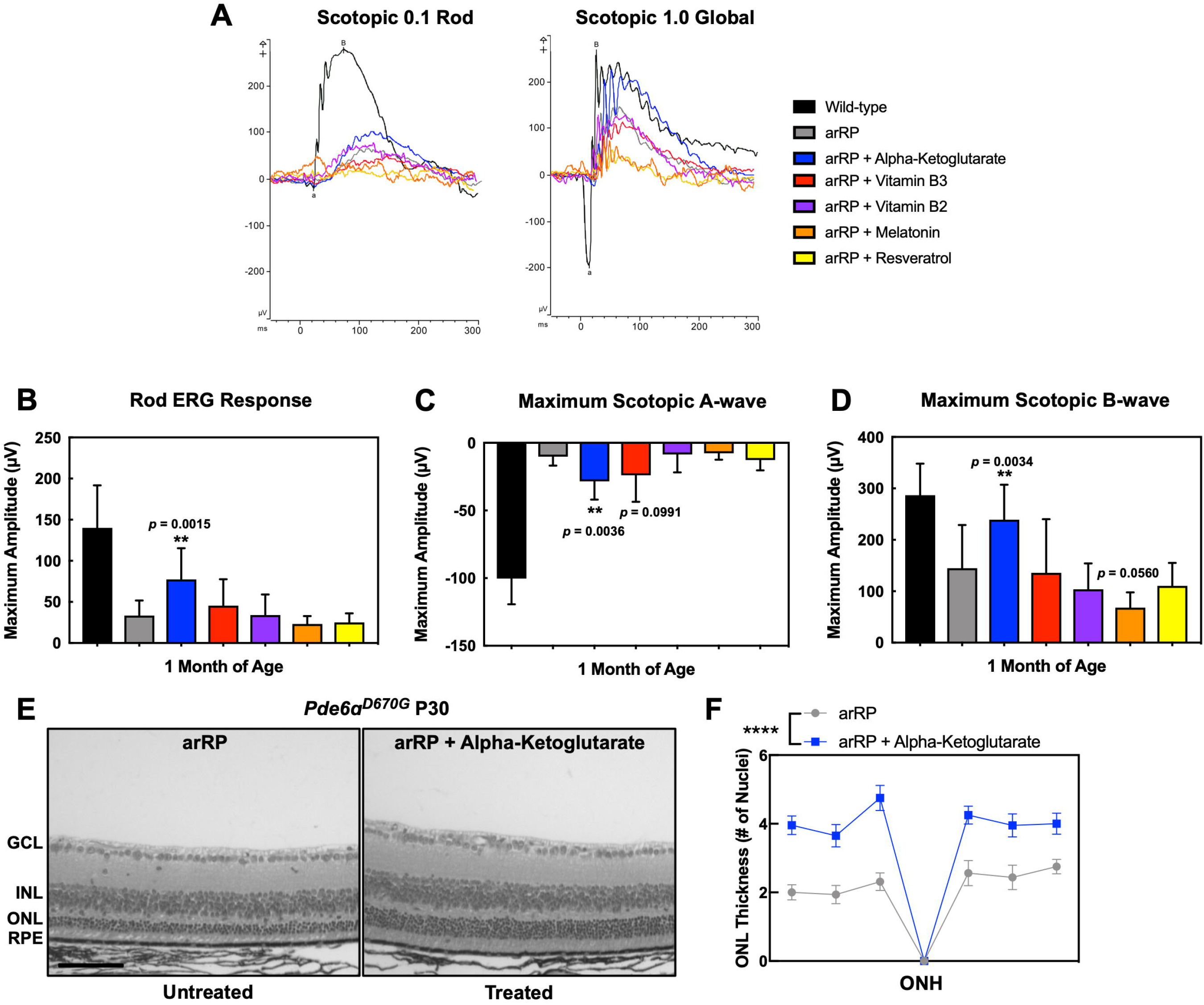
Oral supplementation of single metabolites rescues photoreceptor cell survival and visual function in the arRP preclinical mouse. (**A**) Representative traces of the scotopic 0.1 dim-light rod-specific electroretinography (ERG; left) and scotopic 1.0 global ERG (right) in *Pde6ɑ^D670G^*mice treated with ɑ-KG (blue), vitamin B_2_ (purple), vitamin B_3_ (red), melatonin (orange) or resveratrol (yellow) supplementation in comparison to untreated *Pde6ɑ^D670G^* mice (gray) and wild-type controls (black) at one month of age. Quantification of ERGs from a cohort of mice treated with ɑ-KG (blue), vitamin B_2_ (purple), vitamin B_3_ (red), melatonin (orange) or resveratrol (yellow) supplementation shows significant visual rescue of (**B**) the rod photoreceptor cell response, (**C**) the maximum scotopic a-wave response, and (**D**) the maximum scotopic b-wave response in mice treated with ɑ-KG supplementation (blue) in comparison with wild-type controls (black) or untreated *Pde6ɑ^D670G^* mice (gray) at one month of age. **, *p* < 0.01; ***, *p* < 0.001. N = 5 WT, 6 arRP, 11 arRP + ɑ-KG mice, 8 arRP + vitamin B_3_, 7 arRP + vitamin B_2_, 8 arRP + melatonin and 7 arRP + resveratrol. (**E-F**) Histological analysis of the retinas of untreated *Pde6ɑ^D670G^* mice (left) compared to treated *Pde6ɑ^D670G^*mice (right) shows a significant increase in the thickness of the outer nuclear layer (ONL) by number of photoreceptor cell nuclei in mice treated with ɑ-KG supplementation (blue) at one month of age. GCL, ganglion cell layer; INL, inner nuclear layer; RPE, retinal pigment epithelium. Scale bar = 100 µm. (**F**) Results are displayed as a morphometric quantification (spider graph) of ONL layer thickness superior (left) and inferior (right) to the optic nerve head (ONH). ****, *p* < 0.0001. N = 3 wild-type and 4 arRP + ɑ-KG eyes, with multiple ONL thickness counts per eye as described in Methods. Error bars = SEM.

### Metabolite/lipid imaging of retinal tissue sections

To investigate the lipid and metabolite profiles of *Pde6ɑ^D670G^* mice with and without ɑ-KG treatment, we selected 35 fresh frozen tissue samples and performed negative ion mode (−5 kV) desorption electrospray ionization mass spectrometry imaging (DESI-MSI; Fig. 8A). Retina tissue samples were harvested from wild-type (n =8), untreated *Pde6ɑ^D670G^*(n =8), and ɑ-KG treated *Pde6ɑ^D670G^* mice (n = 14) at one-month of age. Additionally, whole eye (n =5 for each group) samples were also harvested. For each case, 10-μm frozen sections were used for DESI-MSI. Each pixel (200 µm) in the tissue DESI-MS image is represented with the mass spectrum in the 50–1,000 *m/z* range. The molecular ions identified were small metabolites related to glycolysis, TCA cycle, fatty acids, and phospholipids (Fig. 8B-D). The relative intensity distribution of each individual metabolite in the wild type, arRP, and treated tissue of both retina and the whole eye were shown in 2D chemical heatmaps **(Fig. S4-S5).** Across 35 total tissue samples, 26,341 unique peaks were found in the 50 – 1,000 *m/z* range. Significance analysis of microarrays (SAM) identified 377 differentially-expressed metabolites between untreated *Pde6ɑ^D670G^* and ɑ-KG treated mice. Although SAM returned 377 metabolites with notable fold-change (absolute fold change > 1), to help interpret the biological role of metabolites, we restricted our analysis to peaks for which tandem-MS and high mass resolution analyses were performed by using the LTQ-Orbitrap XL (Thermo Scientific; **Fig. S6-S7; Table S16**). To check whether orally-supplemented metabolites had reached the retina, we first analyzed the changes in relative intensity for ɑ-KG (*m/z* 145.014) in our DESI-MSI data. Although DESI-MSI did detect ɑ-KG (**Fig. S8**) in some pixels, there was limited change between *Pde6ɑ^D670G^* and ɑ-KG treated groups. This is possible if the orally supplemented ɑ-KG is being consumed for the production of other metabolites in TCA and in glutamate-glutamine conversion pathways, which were ascertained next.

**Fig. 8.**
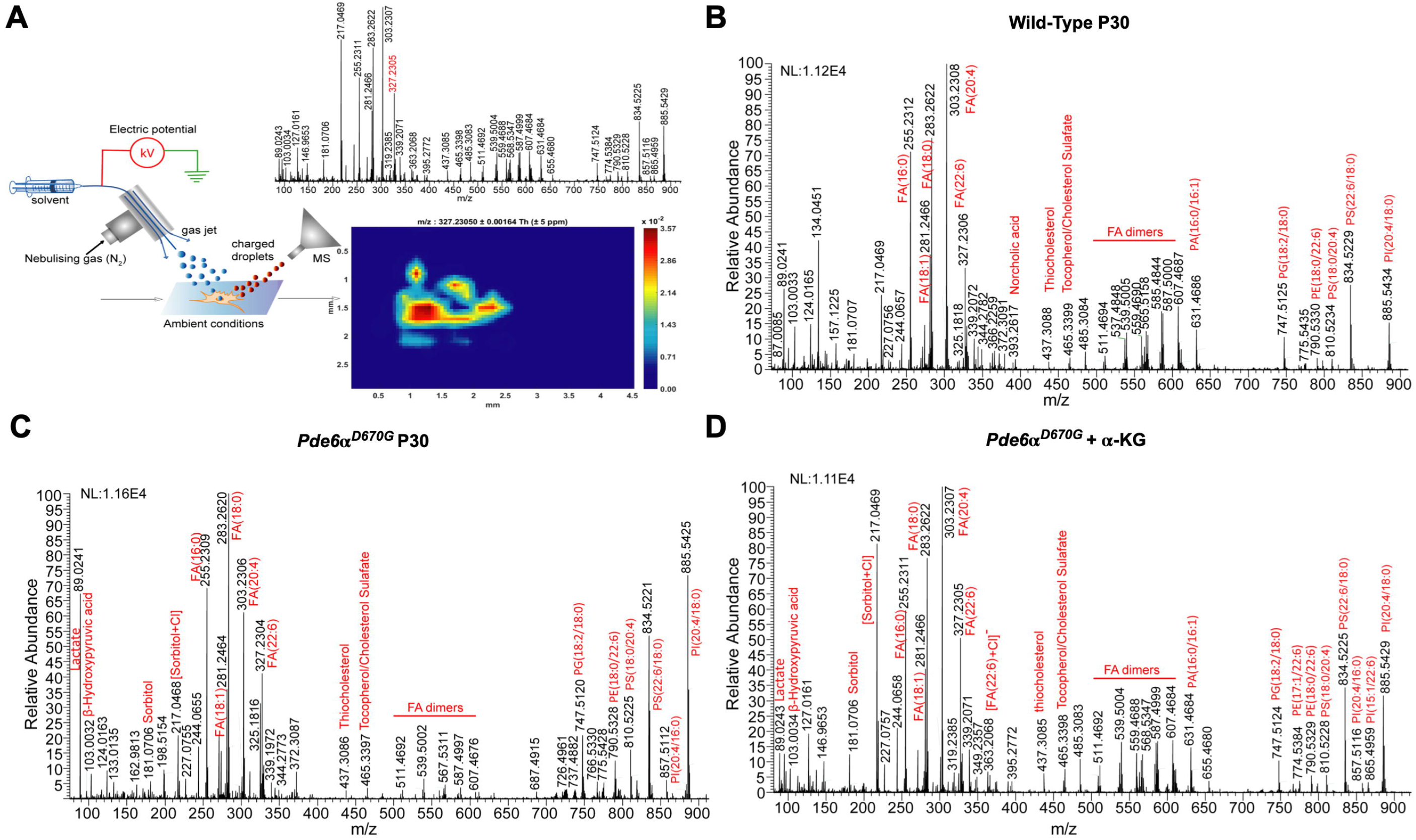
DESI-MSI of retinal tissue sections: (**A**) Schematic representation of the DESI-MSI in the range *m/z* 50-1000. The impinging primary charged droplets desorb the surface metabolites into the secondary charged droplets, followed by the ion transfer to the mass spectrometer. The 2D-chemical map of docosahexaenoic acid (DHA; peak listed in red font) is shown with pixel size 200 µm. Raw MS spectrum across the pixels in a row is shown for (**B**) wild type retina tissue (**C**) arRP retina tissue and (**D**) ɑ-KG treated retina tissue. Red labels denote peaks assigned to identified metabolites and lipids. NL, normalization level; FA, fatty acid; PA, phosphatidic acid; PE, phosphatidylethanolamine; PG, phosphatidylglycerol; PI, phosphatidylinositol; PS, phosphatidylserine.

Identification of differentially distributed metabolites from SAM revealed several altered pathways between untreated *Pde6ɑ^D670G^* and ɑ-KG treated groups that were consistent with our proteomic findings: pyrimidine and purine metabolism (uridine, dihydrouridine, and thymidine), docosahexaenoic and arachidonic acid (lipid metabolism), glutamine and glutamyl-alanine (glutamine/glutamate conversion and aminobutryate degradation), and ascorbic acid (oxidative metabolism). We found intermediates in the TCA cycle that were highly elevated in ɑ-KG treated mice compared to the untreated *Pde6ɑ^D670G^*mice. For example, α-KG is broken down and reduced to succinyl-CoA, which is then converted to succinic acid. From DESI-MSI imaging and SAM, we noted that succinic acid (*m/z* 117.01) levels are elevated in the treated state (fold change 1.3, FDR < 0.05) compared to the untreated group (Fig. 9A). This indicates that the consumption of the supplemental α-KG in the treatment group is reflected in higher levels of succinic acid. Additionally, aconitic acid (*m/z* 173.0) is an intermediate in the conversion of citrate to isocitrate in the TCA cycle, which can also be converted from ɑ-KG. Aconitic acid levels were elevated in the treated state (fold change 2.5, FDR < 0.05), compared to wild-type and untreated *Pde6ɑ^D670G^* levels (Fig. 9B). Since it is known that ɑ-KG can act as a substrate for amino acid synthesis, particularly glutamine, we investigated whether the consumption of ɑ-KG was reflected in glutamine synthesis pathways. Glutamine levels were elevated (fold change 1.87, FDR < 0.05) in treated tissues compared to untreated *Pde6ɑ^D670G^* mice (Fig. 9C). Glutamate levels were slightly increased in treated tissues compared to untreated *Pde6ɑ^D670G^* mice (Fig. 9D). Therefore, DESI-MSI provided some indication that supplemental ɑ-KG was being consumed to increase levels of TCA metabolites such as aconitic acid and glutamine levels, which likely play a role in the cell’s antioxidant response [53].

**Fig. 9.**
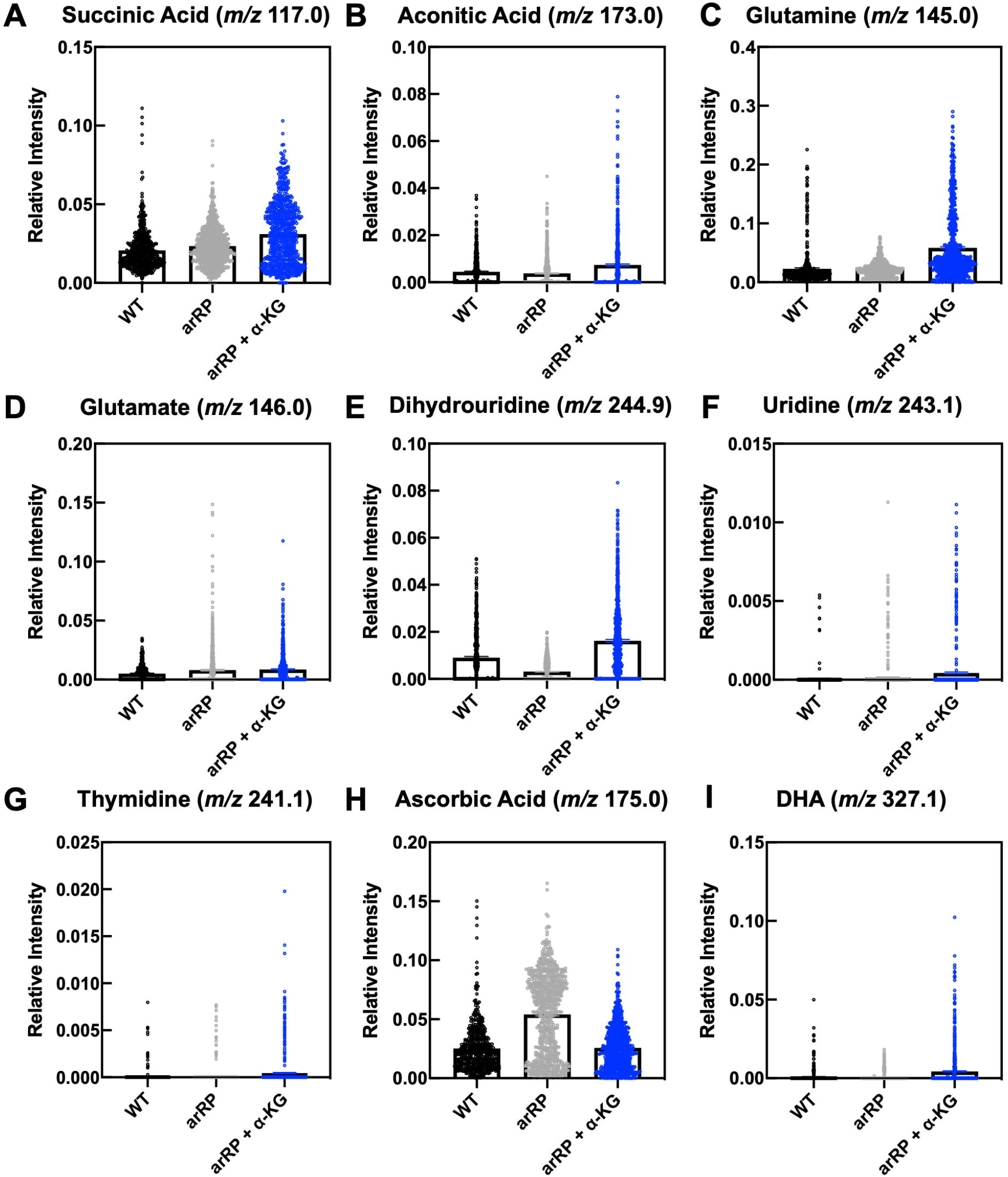
Distributions of retinal lipids and metabolite relative intensities across wild-type, arRP, and ɑ-KG treated mice from DESI-MSI data: Distribution of (**A**) succinic acid (*m/z* 117.01), (**B**) aconitic acid (*m/z* 173.0), (**C**) glutamine (*m/z* 145.0), (**D**) glutamate (*m/z* 146.0), (**E**) dihydrouridine (*m/z* 244.9), (**F**) uridine (*m/z* 243.1), (**G**) thymidine (*m/z* 241.1), (**H**) ascorbic acid, and (**I**) docosahexaenoic acid (DHA; *m/z* 327.1) across wild-type, arRP, and ɑ-KG treated mice. Results are displayed as scatter plots of the relative intensity of the corresponding metabolite in each treatment group. Boxes represent the mean ± SEM.

Consistent with the pathway representation of our vitreous proteomic biomarkers, DESI-MSI independently detected altered levels of metabolites related to pyrimidine metabolism. Notably, our results from SAM revealed that levels of dihydrouridine (*m/z* 244.98) were lower in *Pde6ɑ^D670G^* mice and restored following ɑ-KG treatment (fold change 18.1, FDR < 0.05; Fig. 9E) [54]. Similarly, levels of thymidine (*m/z* 241.17) and uridine (*m/z* 243.16) were elevated in ɑ-KG treated mice compared to *Pde6ɑ^D670G^*and wild-type mice, further suggesting that ɑ-KG treatment influenced pyrimidine and purine metabolism (Fig. 9F-G). This trend of altered nucleotide metabolism in arRP has been found in other murine metabolomics analyses [55]. Weiss, *et al.* had previously used broad-spectrum metabolomics to report links between altered levels of thymidine and other pyrimidine nucleotide metabolic products and RP in a different RP murine model [55]. Overall, our work, in conjunction with other previously published efforts, suggests that pyrimidine metabolism is altered in RP and may be related to mitochondrial DNA replication pathways [53, 54].

The proteomics data indicated the downregulation of oxidative pathways in *Pde6ɑ^D670G^*retina compared to wild-type retina. We therefore looked to determine if any of the identified metabolites from SAM reflected a change in oxidative pathways between the disease and treated groups. Ascorbic acid (*m/z* 175.06) was found at significantly higher intensities in untreated disease samples compared to the treated group (fold change 1.9, FDR < 0.05; Fig. 9H). It is well-known that RP is characterized by rod photoreceptor cell death followed by slower cone death, where cone death is associated with oxidative damage [56]. Our DESI-MSI results revealed that the untreated disease samples had a higher distribution of ascorbic acid, which is a known antioxidant. This higher intensity may reflect the tissue’s attempt to combat the oxidative damage that is characteristic of RP pathogenesis. This hypothesis is further supported by the decrease in ascorbic acid in the treated group compared to untreated *Pde6ɑ^D670G^*mice. Additionally, there were metabolites that did not change significantly in the disease state but were elevated by ɑ-KG treatment. For example, docosahexaeonic acid (DHA; *m/z* 327.12) levels were elevated (fold change 1.9, FDR < 0.05) in the treated mice compared to untreated mice (Fig. 9I). DHA accounts for a significant fraction of omega-3 fats in the retina, and studies have found links between reduction in DHA with altered retinal function, including in RP [56, 57]. This result is consistent with the numerous suggested protective roles of DHA in the retina, including anti-inflammatory, anti-angiogenesis, and anti-neurotoxic mechanisms [58].

## Discussion

Liquid biopsies and the analysis of fluid compartments near diseased tissues represents an innovative tool in precision health to overcome the current limitations of tissue biopsies. This advancement is exemplified in the personalized treatment of cancer, where circulating tumor-derived material (e.g. circulating tumor DNA/RNA, extracellular vesicles, and cells) are detected in the blood [59, 60]. These cellular biomarkers of the tumor microenvironment aid clinicians to better determine the appropriate therapeutic approach and predict prognosis. Neurodegenerative diseases, however, currently lack early detection methods for diagnosis and screening. In the case of retinal degenerative diseases, we have previously shown that liquid biopsies of the adjacent vitreous fluid can capture intracellular events in the neural retinal network that can guide diagnosis and treatment [28, 47, 61]. Advancements in this approach could allow for serial sampling and longitudinal monitoring of disease progress without the need to biopsy the neurosensory retina [27, 29]. Our proteomic analysis of end-stage human arRP vitreous highlighted molecular pathways that might be affected during the progression of neuronal cell death and retinal remodeling. In order to gain further insight into the earlier disease stages, we utilized a preclinical murine model of arRP.

We examined the retina and vitreous proteomes from *Pde6ɑ^D670G^* mice at early, mid, and late stages of neurodegeneration. In the early stage, we observed rod phototransduction proteins were depleted in the retina and elevated within the vitreous before significant loss of the photoreceptor cell nuclei is detectable by histological analysis. This suggests that molecular biomarkers are likely to appear in the human vitreous much earlier than the clinical biomarkers currently used for diagnosis. In human patients, liquid biopsies could similarly be used as an early diagnostic to confirm the onset of photoreceptor degeneration, even without diagnosis of a specific genetic mutation by genetic testing. Additionally, we found that the vitreous proteomes differed in the arRP mouse model at early-, mid-, and late-stage of disease. It may be possible to use this liquid biopsy approach to estimate the molecular stage of disease based on the proteomic content of the vitreous. Comparison of proteomes before and after therapeutic approaches can also be studied using liquid biopsy and may have important implications in gene therapy studies. Further, our DESI-MSI data complements our understanding of key altered pathways detected by our proteomics and pathway analysis approach. Specifically, DESI-MSI provided a precise approach to investigate how changes at the protein level are reflected in the retinal metabolome of arRP mice following ɑ-KG treatment. Using DESI-MSI and SAM, we identified several significantly altered metabolites between disease and treated groups, across multiple pathways, including TCA, glutamine synthesis, pyrimidine metabolism, oxidative pathways, and fatty acid metabolism. It is likely that DESI-MSI in tandem with studying the vitreous proteome can help identify potential biomarkers that help elucidate effects of metabolite therapy.

At the early timepoint, the *Pde6ɑ^D670G^* mice lose proteins critical for oxidative phosphorylation and aerobic metabolism, likely depressing protein turnover and leading to the retinal cell death of the rod photoreceptors. Although the number of patient biopsies was limited and the disease stage was late, there were overlapping proteins in the vitreous proteome of the *Pde6ɑ^D670G^* mice and humans. The loss of proteins involved in critical metabolic pathways from the retina was supported by our previous proteomics of the human retina that highlighted the sensitivity of the neural retinal network to changes within the metabolic pathways [13]. Interestingly, previous research studies have shown that metabolic rewiring is likely to be both a cause and a consequence of photoreceptor degeneration [62], and that the disruption of normal energy metabolism plays a key role in photoreceptor degeneration in the *rd10* RP mouse model [55]. We also found that there were significant alterations in purine and pyrimidine nucleotide metabolism pathways at disease onset in our arRP mouse model, similar to the results in the *rd10* RP mouse study (**Table S12**). Taken together, the proteomic profiles suggest targeting the affected metabolic pathways by replenishing them though dietary supplementation.

The most effective therapy tested was ɑ-KG, a key component of the TCA cycle that acts on various downstream metabolic pathways within the cell and determines the overall rate of the energetic process [55] as well as enhancing reductive carboxylation and supporting redox homeostasis in photoreceptors [63, 64]. In addition to its metabolic roles, ɑ-KG can be converted to glutamine, and subsequently glutathione GSH, thereby facilitating the synthesis of potent antioxidants in the cell [53]. These antioxidant properties may have beneficial effects in the context of RP therapy. Additional studies have shown that ɑ-KG supplementation can extend the lifespan of *C. elegans* through the inhibition of ATP synthase and TOR [65] as well as increase circulating plasma levels of insulin and growth hormones in humans [66]. In the mouse, ɑ-KG derivatives have been delivered as a treatment for hypoxic conditions without any notations of adverse side effects [67–69]. We found that oral supplementation with ɑ-KG alone provided significant visual rescue of the rods, cones, and inner retina visual responses through at least one month of age. Additional studies are needed to determine the lowest dosage of ɑ-KG and its durability. Although it seems likely that the metabolic effect acts directly on retinal cells, further studies are needed to test whether some rescue effect may be due to the metabolites acting indirectly via other organs in the body. In this study, mice treated with ɑ-KG displayed a smaller body stature than controls, and therefore for translation into humans the delivery of ɑ-KG may be best optimized as an intraocular therapeutic.

The ketogenic diet has shown to have neuroprotective effects in the context of several neurodegenerative diseases, such as Alzheimer’s, and Parkinson’s disease [70]. The neuroprotective effect of the ketogenic diet has been hypothesized to be a result of the biochemical changes resulting from glycolytic inhibition, increased acetyl-CoA production, and formation of ketone bodies in the cell [71]. These biochemical changes have proven effective in targeting cancer cells, which primarily use glycolysis as their main source of ATP [72]. Photoreceptors are similarly sensitive to glycolytic inhibition but can attenuate the impact of glycolytic inhibition by using alternative sources of ATP (e.g. fatty acid beta-oxidation) [17, 73]. The high rate of aerobic ATP synthesis in photoreceptors can lead to the production of harmful reactive oxygen species (ROS), which are implicated in the progression of retinal degenerative diseases [48]. We have previously shown that different anatomical regions of the human retina are susceptible to oxidative stress, highlighting the importance of antioxidant availability in the retina [13]. Ketone bodies increase mitochondrial respiration via increased ATP production and reduce ROS formation by increasing NADH oxidation, thereby improving the efficiency of the mitochondrial respiratory chain complex [70]. Targeting the TCA cycle and fatty acid synthase pathway by acetyl-CoA production via the ketogenic diet led to increased global retinal visual function compared to controls by one month of age. As the ketogenic diet could not be provided until weaning age, the effect was limited based on beginning the dietary treatment after approximately half of the neuronal photoreceptor cells have already been lost. Nevertheless, significant photoreceptor cell survival was detectable by histological analysis after treatment with the ketogenic diet. This suggests that earlier delivery of metabolites targeting the TCA cycle may lead to prolonged visual rescue in the *Pde6ɑ^D670G^* mouse model and neuroprotection of the photoreceptor cells and inner retinal network.

Other metabolites tested in our arRP preclinical mouse model were less effective. Nicotinamide adenine dinucleotide (NAD), available from vitamin B_3_ supplementation, is a critical co-enzyme for aerobic metabolism and OXPHOS with no known dosage-related toxicity. In particular, retinal levels of NAD decrease with age and may leave neuronal cells susceptible to insults and disease [74]. Furthermore, vitamin B_3_ supplementation has shown positive effects toward enhancing energy metabolism and proper functioning of the central nervous system [75–77] as well as acting as a protective agent against glaucoma and age-related neurodegeneration [74]. Flavin mononucleotide (FMN) and flavin adenine dinucleotide (FAD), which can be delivered via vitamin B_2_ supplementation, are critical for enzymatic reactions within the electron transport chain and the TCA cycle. Vitamin B_2_ acts as an antioxidant by promoting regeneration of glutathione and is indispensable for cellular growth and supplementation with vitamin B_2_ has shown possible beneficial effects for prevention of hyperhomocysteinemia, cataracts and migraine headaches [78]. We found that targeting OXPHOS by oral supplementation with vitamins B_2_ or B_3_ provided an electrophysiological rescue effect in some, but not all, neuronal cells of treated mice compared to their respective controls. Further studies are needed to test combination therapy with our effective treatment groups (ɑ-KG, the ketogenic diet, and/or vitamins B_2_ and B_3_) that may have a synergistic effect in rescuing the neuronal cell loss and more closely reproduce a strategy to use in human arRP patients. Longer-acting structural analogs of these metabolites may also provide a longer lasting effect.

We also tested metabolites that act on the sirtuin pathway and may indirectly affect cellular metabolism. Melatonin is produced within the retina and is known to stimulate sirtuin activity in mitochondria, which deacetylates many of the enzymes involved within the TCA cycle. Previous studies have shown that supplementation with melatonin has no known toxicity affects when taken during pregnancy in both humans and mice [79–81]. Additionally, resveratrol is a plant-derived polyphenol that has been widely studied for its potential benefits as a protective agent against cancers, mutagenesis, and a variety of other biological functions [81]. It also acts to stimulate the sirtuin pathway, and this stimulation as well as its antioxidative actions are thought to provide a neuroprotective effect in both rat and mouse studies [81]. Supplementation with resveratrol has no known toxicity in humans even at high dosages, and the only affect noted when taken during pregnancy in mice was a slight increase in the body weight of the pups after birth [82, 83]. However, in our study, oral supplementation with either melatonin or resveratrol did not provide a neuroprotective effect on the photoreceptor cells in our preclinical model of arRP. This suggests that directly targeting the TCA cycle may be the most beneficial approach for future RP therapy.

The proteomic and DESI-MSI approach in this study provided key pathways for targeting and testing therapeutics, and delivery of metabolites within these critical pathways in the arRP preclinical mouse model showed efficacious neuroprotective effects on the inner retina. The results from these studies lay the groundwork for future experiments to address how the TCA cycle and aerobic metabolism influence the photoreceptors and their signaling to the inner retinal network, and thus improve upon neuroprotective approaches targeting photoreceptor cell metabolism. Overall, the key metabolic pathways (OXPHOS and TCA cycle) detected in our proteomics screen are critical targets for therapeutics for RP, independent of specific genetic mutations and may be applicable to other human neurodegenerative disorders.

## Acknowledgements

We thank Jing Yang, Daniel A. Machlab, Teja Chemudupati, and Louis K. Chang for technical assistance. We thank Bioproximity for assistance with mass spectrometry data collection. We thank Patsy Nishina for mouse model resources. We thank James Hurley and Jianhai Du at the University of Washington for advice and discussions.

## Author contributions

Dr. Mahajan had full access to all the data in the study and takes responsibility for the integrity of the data and the accuracy of the data analysis. Study concept and design: KJW, SHT, AGB and VBM. Acquisition of data: KJW, GV, KV, VS, JDS, VBM. Analysis and interpretation of data: KJW, GV, KV, VS, SHT, AGB, VBM. Drafting of the manuscript: KJW, GV, VBM. Critical revision of the manuscript for important intellectual content: KJW, GV, VS, RNZ, SHT, AGB, VBM. Statistical analysis: KJW, GV, VS. Obtained funding: VBM. Administrative, technical, and material support: VBM. Study supervision: VBM.

## Data and materials availability

All data associated with this study are present in the paper or the Supplementary Materials. The mass spectrometry proteomics data have been deposited to Mendeley Data with the dataset identifiers (DOI): 10.17632/x9mkhv743y.2 and 10.17632/vn5c8zh5wd.2.

---

**Fig. S1. Gene ontology (GO) distributions of retina samples.** Identified retinal proteins from the wild-type and *Pde6ɑ^D670G^* mice at P15, P28, and P90. Gene ontology analysis categorized each protein group by biological process, molecular function, and cellular compartment.

**Fig. S2. Gene ontology (GO) distributions of vitreous samples.** Identified vitreous proteins from the wild-type and *Pde6ɑ^D670G^* mice at P15, P28, and P90. Gene ontology analysis categorized each protein group by biological process, molecular function, and cellular compartment.

**Fig. S3. Gene ontology (GO) distributions of candidate vitreous biomarkers.** Gene ontology (GO) categorization of the 446 protein biomarkers by **(A)** cellular compartment and **(B)** biological process.

**Fig. S4. DESI-MSI chemical maps of retina.** The 2D-chemical maps of individual metabolites are shown for **(A)** wild-type **(B)** arRP and **(C)** ɑ-KG treated mice. PA, phosphatidic acid; PE, phosphatidylethanolamine; PG, phosphatidylglycerol; PI, phosphatidylinositol; PS, phosphatidylserine; DHA, docosahexaenoic acid, CI, collision induced.

**Fig. S5. DESI-MSI chemical maps of whole eye.** The respective raw MS spectrum across all the pixels in a row and the 2D-chemical maps of individual metabolites is shown for **(A)** wild-type **(B)** arRP and **(C)** ɑ-KG treated mice. NL, normalization level.

**Fig. S6. Tandem-MS analysis of extracted metabolites from the retinal tissue:** Collision induced dissociation analysis (CID), is shown for small metabolites, fatty acids, and lipids to identify molecular species based on fragmentation pattern: **(A)** Taurine; **(B)** Glutamate; **(C)** Malate; **(D)** Norepinephrine; **(E)** N-acetyl aspartate; **(F)** Sorbitol; **(G)** [Sorbitol + collision-induced (CI)] adduct; **(H)** Malate; **(I)** Glutaconic acid; **(J)** Thymidine; **(K)** Uridine; **(L)** Dihydro-uridine; **(M)** Deoxy-uridine; **(N)** Taurine (dimer); **(O)** Arachidonic acid; **(P)** Glutathione; **(Q)** Phytanic acid. **(R)** Docosahexaenoic acid (DHA); **(S)** Phosphatidylethanolamine (PE, 18:0/22:6). **(T)** Phosphatidylserine (PS, 22:6/18:0). **(U)** Phosphatidylinositol (PI, 20:4/16:0). **(V)** Phosphatidylinositol (PI, 20:4/18:0). **(W)** Phosphatidylglycerol (PG, 18:2/18:0). **(X)** Phosphatidic acid (PA, 20:4/19:0). **(Y)** 5-LGlutamyl-L-alanine. **(Z)** Phosphatidylserine (PS, 18:0/20:4). NL, normalization level.

**Fig. S7. DESI-MS analysis of extracted metabolites from the retinal tissue:** Electrospray ionization is performed on the retina tissue extraction in solvents in the mass range *m/z* 50-1000. Red labels denote peaks assigned to identified metabolites and lipids. NL, normalization level.

**Fig. S8. Distribution of ɑ-KG across wild-type, arRP, and ɑ-KG treated mice:** Levels of ɑ-KG (*m/z* 145.01) are not significantly different between arRP and treated mice (fold change 1.26, false discovery rate < 0.05). The supplemental ɑ-KG in treated state is consumed and reflected in the increased intensity of other metabolites in TCA and glutamate-glutamine conversion pathways.

## List of Supplementary Materials

### Supplementary Figures

**Fig. S1.** Gene ontology of the *Pde6ɑ^D670G^* retina at different stages of degeneration.

**Fig. S2.** Gene ontology of the *Pde6ɑ^D670G^* vitreous at different stages of degeneration.

**Fig. S3.** Gene ontology distributions of candidate vitreous biomarkers.

**Fig. S4.** DESI-MSI chemical maps of retina from wild-type, arRP, and ɑ-KG treated mice.

**Fig. S5.** DESI-MSI chemical maps of whole eye from wild-type, arRP, and ɑ-KG treated mice.

**Fig. S6.** Tandem-MS analysis of extracted metabolites from the retinal tissue.

**Fig. S7**. DESI-MS analysis of extracted metabolites from the retinal tissue.

**Fig. S8**. Distribution of ɑ-KG across wild-type, arRP, and ɑ-KG treated mice.

### Supplementary Tables

**Table S1.** Vitreous biopsies from arRP patients and unaffected controls.

**Table S2.** Proteomic content of human vitreous samples by LC-MS/MS analysis.

**Table S3**. Relative quantification of proteins identified in human vitreous samples.

**Table S4.** Proteomic content of *Pde6ɑ^D670G^* retina and vitreous by LC-MS/MS analysis.

**Table S5.** Distinct peptides identified in *Pde6ɑ^D670G^* retina and vitreous samples.

**Table S6.** Protein groups identified in *Pde6ɑ^D670G^* retina and vitreous samples.

**Table S7.** Relative quantification and comparison of proteins identified in *Pde6ɑ^D670G^* retina samples.

**Table S8.** Relative quantification and comparison of proteins identified in *Pde6ɑ^D670G^*vitreous samples.

**Table S9.** Differentially-expressed *Pde6ɑ^D670G^* retina and vitreous proteins in all disease stages.

**Table S10.** Differentially-expressed *Pde6ɑ^D670G^* retina and vitreous proteins in early disease stages.

**Table S11**. Candidate biomarkers from *Pde6ɑ^D670G^* vitreous samples.

**Table S12.** Pathway representation of down-regulated proteins in the *Pde6ɑ^D670G^*retina.

**Table S13.** Down-regulated proteins in the *Pde6ɑ^D670G^*retina involved in oxidative phosphorylation.

**Table S14.** Down-regulated proteins in the *Pde6ɑ^D670G^* retina involved in the tricarboxylic acid (TCA) cycle.

**Table S15.** Down-regulated proteins in the *Pde6ɑ^D670G^* retina involved in rod outer segment phototransduction.

**Table S16.** Tandem-MS analysis of extracted metabolites from the retinal tissue.

## References

1. Procaccini, C., et al., Role of metabolism in neurodegenerative disorders. Metabolism, 2016. 65(9): p. 1376–90.

2. Lehtonen, S., et al., Dysfunction of Cellular Proteostasis in Parkinson’s Disease. Front Neurosci, 2019. 13: p. 457.

3. Strohm, L. and C. Behrends, Glia-specific autophagy dysfunction in ALS. Semin Cell Dev Biol, 2019.

4. Olivares-Banuelos, T.N., D. Chi-Castaneda, and A. Ortega, Glutamate transporters: Gene expression regulation and signaling properties. Neuropharmacology, 2019.

5. Shukla, M., et al., The role of melatonin in targeting cell signaling pathways in neurodegeneration. Ann N Y Acad Sci, 2019. 1443(1): p. 75–96.

6. Ambrogini, P., et al., Excitotoxicity, neuroinflammation and oxidant stress as molecular bases of epileptogenesis and epilepsy-derived neurodegeneration: The role of vitamin E. Biochim Biophys Acta Mol Basis Dis, 2019. 1865(6): p. 1098–1112.

7. Chi, H., H.Y. Chang, and T.K. Sang, Neuronal Cell Death Mechanisms in Major Neurodegenerative Diseases. Int J Mol Sci, 2018. 19(10).

8. Tsang, S.H. and T. Sharma, Stargardt Disease. Adv Exp Med Biol, 2018. 1085: p. 139–151.

9. Luo, Y.H. and L. da Cruz, The Argus((R)) II Retinal Prosthesis System. Prog Retin Eye Res, 2016. 50: p. 89–107.

10. Cideciyan, A.V., et al., Pseudo-fovea formation after gene therapy for RPE65-LCA. Invest Ophthalmol Vis Sci, 2014. 56(1): p. 526–37.

11. Maguire, A.M., et al., Safety and efficacy of gene transfer for Leber’s congenital amaurosis. N Engl J Med, 2008. 358(21): p. 2240–8.

12. Ashtari, M., et al., The human visual cortex responds to gene therapy-mediated recovery of retinal function. J Clin Invest, 2011. 121(6): p. 2160–8.

13. Velez, G., et al., Proteomic analysis of the human retina reveals region-specific susceptibilities to metabolic- and oxidative stress-related diseases. PLoS One, 2018. 13(2): p. e0193250.

14. Wang, L., P. Tornquist, and A. Bill, Glucose metabolism in pig outer retina in light and darkness. Acta Physiol Scand, 1997. 160(1): p. 75–81.

15. Wang, L., P. Tornquist, and A. Bill, Glucose metabolism of the inner retina in pigs in darkness and light. Acta Physiol Scand, 1997. 160(1): p. 71–4.

16. Graymore, C., Possible Significance of the Isoenzymes of Lactic Dehydrogenase in the Retina of the Rat. Nature, 1964. 201: p. 615–6.

17. Noell, W.K., The effect of iodoacetate on the vertebrate retina. J Cell Comp Physiol, 1951. 37(2): p. 283–307.

18. Chertov, A.O., et al., Roles of glucose in photoreceptor survival. J Biol Chem, 2011. 286(40): p. 34700–11.

19. Narayan, D.S., et al., Glucose metabolism in mammalian photoreceptor inner and outer segments. Clin Exp Ophthalmol, 2017. 45(7): p. 730–741.

20. Vaughn, A.E. and M. Deshmukh, Glucose metabolism inhibits apoptosis in neurons and cancer cells by redox inactivation of cytochrome c. Nat Cell Biol, 2008. 10(12): p. 1477–83.

21. Robertson, J.P., A. Faulkner, and R.G. Vernon, Regulation of glycolysis and fatty acid synthesis from glucose in sheep adipose tissue. Biochem J, 1982. 206(3): p. 577–86.

22. Rajagopal, R., et al., Retinal de novo lipogenesis coordinates neurotrophic signaling to maintain vision. JCI Insight, 2018. 3(1).

23. Joyal, J.S., M.L. Gantner, and L.E.H. Smith, Retinal energy demands control vascular supply of the retina in development and disease: The role of neuronal lipid and glucose metabolism. Prog Retin Eye Res, 2018. 64: p. 131–156.

24. Zhang, L., et al., Reprogramming metabolism by targeting sirtuin 6 attenuates retinal degeneration. J Clin Invest, 2016. 126(12): p. 4659–4673.

25. Skeie, J.M. and V.B. Mahajan, Proteomic landscape of the human choroid-retinal pigment epithelial complex. JAMA Ophthalmol, 2014. 132(11): p. 1271–81.

26. Skeie, J.M. and V.B. Mahajan, Proteomic interactions in the mouse vitreous-retina complex. PLoS One, 2013. 8(11): p. e82140.

27. Skeie, J.M., C.N. Roybal, and V.B. Mahajan, Proteomic insight into the molecular function of the vitreous. PLoS One, 2015. 10(5): p. e0127567.

28. Velez, G., et al., Precision Medicine: Personalized Proteomics for the Diagnosis and Treatment of Idiopathic Inflammatory Disease. JAMA Ophthalmol, 2016. 134(4): p. 444–8.

29. Mahajan, V.B. and J.M. Skeie, Translational vitreous proteomics. Proteomics Clin Appl, 2014. 8(3-4): p. 204–8.

30. Rowell, H.A., A.G. Bassuk, and V.B. Mahajan, Monozygotic twins with CAPN5 autosomal dominant neovascular inflammatory vitreoretinopathy. Clin Ophthalmol, 2012. 6: p. 2037–44.

31. Mahajan, V.B., et al., Calpain-5 mutations cause autoimmune uveitis, retinal neovascularization, and photoreceptor degeneration. PLoS Genet, 2012. 8(10): p. e1003001.

32. Wert, K.J., et al., Gene therapy provides long-term visual function in a pre-clinical model of retinitis pigmentosa. Hum Mol Genet, 2013. 22(3): p. 558–67.

33. Wert, K.J., J. Sancho-Pelluz, and S.H. Tsang, Mid-stage intervention achieves similar efficacy as conventional early-stage treatment using gene therapy in a pre-clinical model of retinitis pigmentosa. Hum Mol Genet, 2014. 23(2): p. 514–23.

34. Tsang, S.H., et al., Role for the target enzyme in deactivation of photoreceptor G protein in vivo. Science, 1998. 282(5386): p. 117–21.

35. Wert, K.J., et al., CAPN5 mutation in hereditary uveitis: the R243L mutation increases calpain catalytic activity and triggers intraocular inflammation in a mouse model. Hum Mol Genet, 2015. 24(16): p. 4584–98.

36. Wert, K.J., et al., Functional validation of a human CAPN5 exome variant by lentiviral transduction into mouse retina. Hum Mol Genet, 2014. 23(10): p. 2665–77.

37. Skeie, J.M., et al., Proteomic analysis of vitreous biopsy techniques. Retina, 2012. 32(10): p. 2141–9.

38. Mahajan, V.B., et al., Mouse eye enucleation for remote high-throughput phenotyping. J Vis Exp, 2011(57).

39. Skeie, J.M., S.H. Tsang, and V.B. Mahajan, Evisceration of mouse vitreous and retina for proteomic analyses. J Vis Exp, 2011(50).

40. Mi, H., et al., PANTHER version 14: more genomes, a new PANTHER GO-slim and improvements in enrichment analysis tools. Nucleic Acids Res, 2019. 47(D1): p. D419–D426.

41. Tusher, V.G., R. Tibshirani, and G. Chu, Significance analysis of microarrays applied to the ionizing radiation response. Proc Natl Acad Sci U S A, 2001. 98(9): p. 5116–21.

42. Tsang, S.H., et al., A novel mutation and phenotypes in phosphodiesterase 6 deficiency. Am J Ophthalmol, 2008. 146(5): p. 780–8.

43. Azar, W.J., et al., IGFBP-2 enhances VEGF gene promoter activity and consequent promotion of angiogenesis by neuroblastoma cells. Endocrinology, 2011. 152(9): p. 3332–42.

44. Park, S.H., K.W. Kim, and J.C. Kim, The Role of Insulin-Like Growth Factor Binding Protein 2 (IGFBP2) in the Regulation of Corneal Fibroblast Differentiation. Invest Ophthalmol Vis Sci, 2015. 56(12): p. 7293–302.

45. Afshari, F.T., et al., Integrin activation or alpha 9 expression allows retinal pigmented epithelial cell adhesion on Bruch’s membrane in wet age-related macular degeneration. Brain, 2010. 133(Pt 2): p. 448–64.

46. Malissen, B. and A.M. Schmitt-Verhulst, Transmembrane signalling through the T-cell-receptor-CD3 complex. Curr Opin Immunol, 1993. 5(3): p. 324–33.

47. Velez, G., et al., Personalized Proteomics for Precision Health: Identifying Biomarkers of Vitreoretinal Disease. Transl Vis Sci Technol, 2018. 7(5): p. 12.

48. Punzo, C., W. Xiong, and C.L. Cepko, Loss of daylight vision in retinal degeneration: are oxidative stress and metabolic dysregulation to blame? J Biol Chem, 2012. 287(3): p. 1642–8.

49. Hartong, D.T., et al., Insights from retinitis pigmentosa into the roles of isocitrate dehydrogenases in the Krebs cycle. Nat Genet, 2008. 40(10): p. 1230–4.

50. D’Andrea Meira, I., et al., Ketogenic Diet and Epilepsy: What We Know So Far. Front Neurosci, 2019. 13: p. 5.

51. Smilowitz, J.T., et al., The human milk metabolome reveals diverse oligosaccharide profiles. J Nutr, 2013. 143(11): p. 1709–18.

52. Wishart, D.S., et al., HMDB 4.0: the human metabolome database for 2018. Nucleic Acids Res, 2018. 46(D1): p. D608–D617.

53. Liu, S., L. He, and K. Yao, The Antioxidative Function of Alpha-Ketoglutarate and Its Applications. Biomed Res Int, 2018. 2018: p. 3408467.

54. Scaglia, F. and L.J. Wong, Human mitochondrial transfer RNAs: role of pathogenic mutation in disease. Muscle Nerve, 2008. 37(2): p. 150–71.

55. Weiss, E.R., et al., Broad spectrum metabolomics for detection of abnormal metabolic pathways in a mouse model for retinitis pigmentosa. Exp Eye Res, 2019. 184: p. 135–145.

56. Komeima, K., et al., Antioxidants reduce cone cell death in a model of retinitis pigmentosa. Proc Natl Acad Sci U S A, 2006. 103(30): p. 11300–5.

57. Hoffman, D.R., et al., Impaired synthesis of DHA in patients with X-linked retinitis pigmentosa. J Lipid Res, 2001. 42(9): p. 1395–401.

58. Querques, G., R. Forte, and E.H. Souied, Retina and omega-3. J Nutr Metab, 2011. 2011: p. 748361.

59. Alimirzaie, S., M. Bagherzadeh, and M.R. Akbari, Liquid biopsy in breast cancer: A comprehensive review. Clin Genet, 2019. 95(6): p. 643–660.

60. De Rubis, G., S. Rajeev Krishnan, and M. Bebawy, Liquid Biopsies in Cancer Diagnosis, Monitoring, and Prognosis. Trends Pharmacol Sci, 2019. 40(3): p. 172–186.

61. Velez, G., et al., Therapeutic Drug Repositioning Using Personalized Proteomics of Liquid Biopsies. JCI Insight, 2017. In Press.

62. Du, J., et al., How Excessive cGMP Impacts Metabolic Proteins in Retinas at the Onset of Degeneration. Adv Exp Med Biol, 2018. 1074: p. 289–295.

63. Jiang, L., et al., Reductive carboxylation supports redox homeostasis during anchorage-independent growth. Nature, 2016. 532(7598): p. 255–8.

64. Mullen, A.R., et al., Reductive carboxylation supports growth in tumour cells with defective mitochondria. Nature, 2011. 481(7381): p. 385–8.

65. Chin, R.M., et al., The metabolite alpha-ketoglutarate extends lifespan by inhibiting ATP synthase and TOR. Nature, 2014. 510(7505): p. 397–401.

66. Wu, N., et al., Alpha-Ketoglutarate: Physiological Functions and Applications. Biomol Ther (Seoul), 2016. 24(1): p. 1–8.

67. Hou, P., et al., Intermediary metabolite precursor dimethyl-2-ketoglutarate stabilizes hypoxia-inducible factor-1alpha by inhibiting prolyl-4-hydroxylase PHD2. PLoS One, 2014. 9(11): p. e113865.

68. MacKenzie, E.D., et al., Cell-permeating alpha-ketoglutarate derivatives alleviate pseudohypoxia in succinate dehydrogenase-deficient cells. Mol Cell Biol, 2007. 27(9): p. 3282–9.

69. Tennant, D.A., et al., Reactivating HIF prolyl hydroxylases under hypoxia results in metabolic catastrophe and cell death. Oncogene, 2009. 28(45): p. 4009–21.

70. Wlodarek, D., Role of Ketogenic Diets in Neurodegenerative Diseases (Alzheimer’s Disease and Parkinson’s Disease). Nutrients, 2019. 11(1).

71. Puchalska, P. and P.A. Crawford, Multi-dimensional Roles of Ketone Bodies in Fuel Metabolism, Signaling, and Therapeutics. Cell Metab, 2017. 25(2): p. 262–284.

72. Weber, D.D., S. Aminazdeh-Gohari, and B. Kofler, Ketogenic diet in cancer therapy. Aging (Albany NY), 2018. 10(2): p. 164–165.

73. Joyal, J.S., et al., Retinal lipid and glucose metabolism dictates angiogenesis through the lipid sensor Ffar1. Nat Med, 2016. 22(4): p. 439–45.

74. Williams, P.A., et al., Vitamin B3 modulates mitochondrial vulnerability and prevents glaucoma in aged mice. Science, 2017. 355(6326): p. 756–760.

75. Peechakara, B.V. and M. Gupta, Vitamin B3, in StatPearls. 2019: Treasure Island (FL).

76. Djadjo, S. and T. Bajaj, Niacin (Nicotinic Acid), in StatPearls. 2019: Treasure Island (FL).

77. Mills, K.F., et al., Long-Term Administration of Nicotinamide Mononucleotide Mitigates Age-Associated Physiological Decline in Mice. Cell Metab, 2016. 24(6): p. 795–806.

78. Peechakara, B.V. and M. Gupta, Vitamin B2 (Riboflavin), in StatPearls. 2019: Treasure Island (FL).

79. Mayo, J.C., et al., Melatonin and sirtuins: A “not-so unexpected” relationship. J Pineal Res, 2017. 62(2).

80. Guan, S., et al., Effects of Melatonin on Early Pregnancy in Mouse: Involving the Regulation of StAR, Cyp11a1, and Ihh Expression. Int J Mol Sci, 2017. 18(8).

81. Ramis, M.R., et al., Caloric restriction, resveratrol and melatonin: Role of SIRT1 and implications for aging and related-diseases. Mech Ageing Dev, 2015. 146–148: p. 28–41.

82. Poudel, R., et al., Effects of resveratrol in pregnancy using murine models with reduced blood supply to the uterus. PLoS One, 2013. 8(5): p. e64401.

83. Zheng, S., et al., Maternal resveratrol consumption and its programming effects on metabolic health in offspring mechanisms and potential implications. Biosci Rep, 2018. 38(2).

