## Supplementary figures and images for "Metabolite therapy guided by liquid biopsy proteomics delays retinal neurodegeneration"

### Supplemental Figure 1

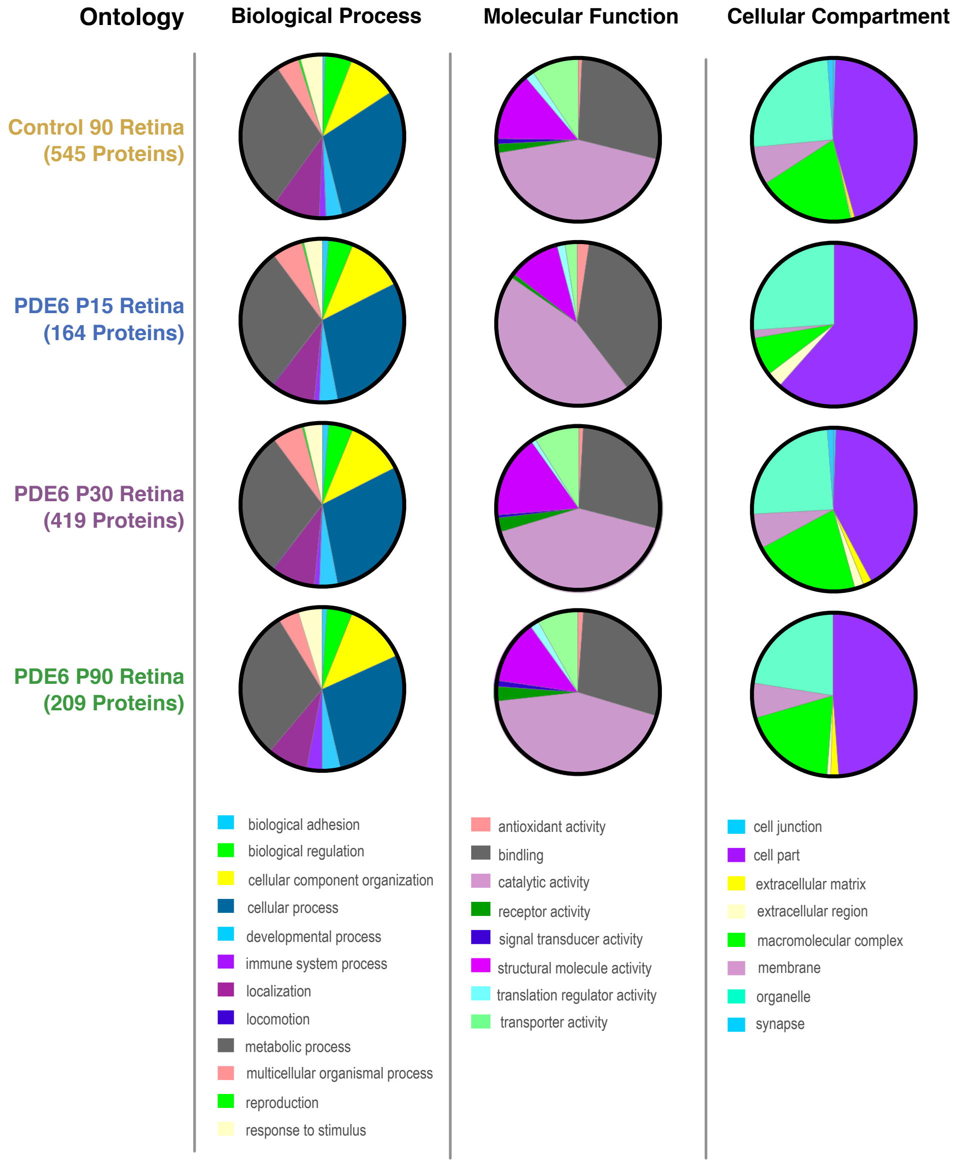

### Supplemental Figure 2

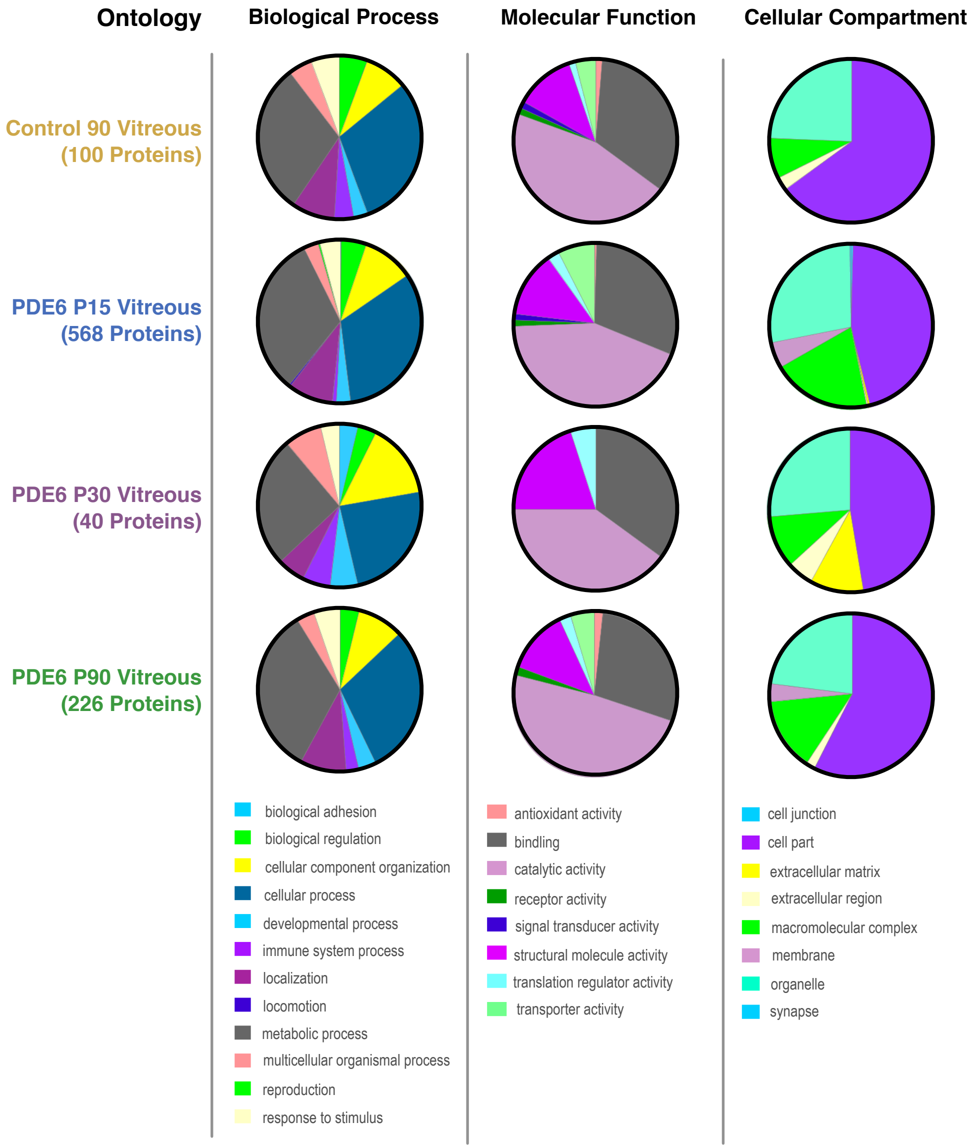

### Supplemental Figure 3

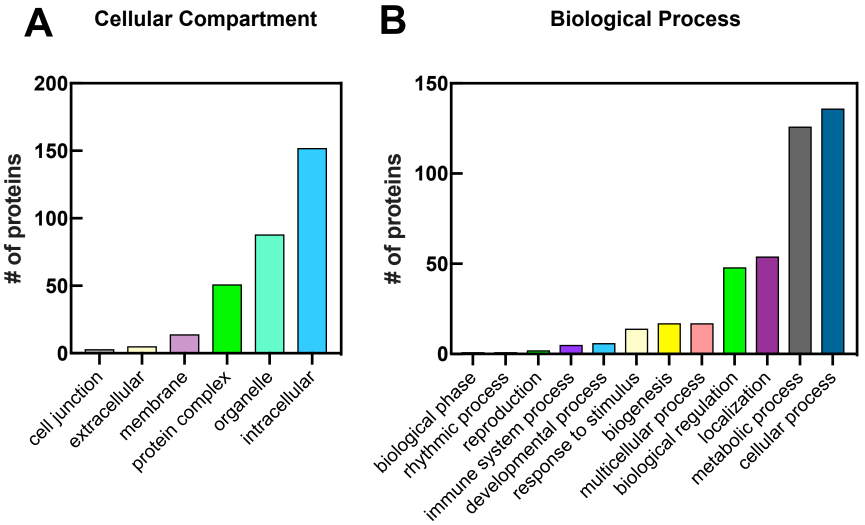

### Supplemental Figure 4

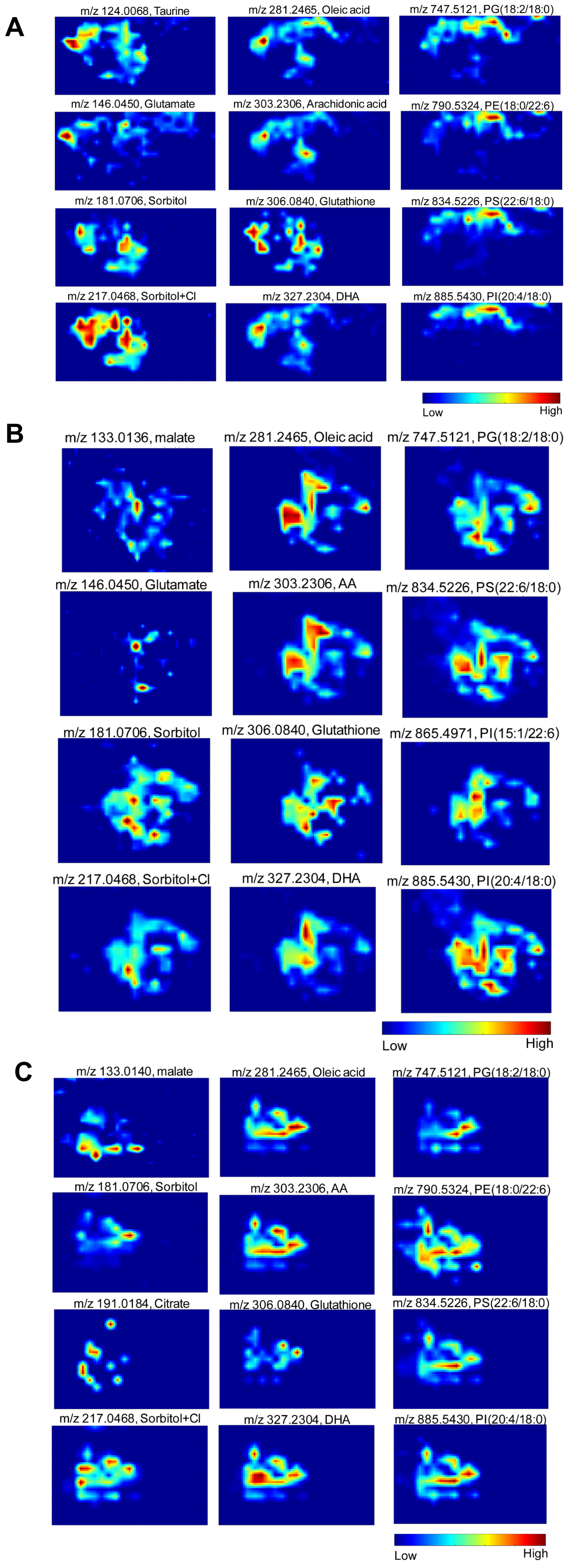

### Supplemental Figure 5

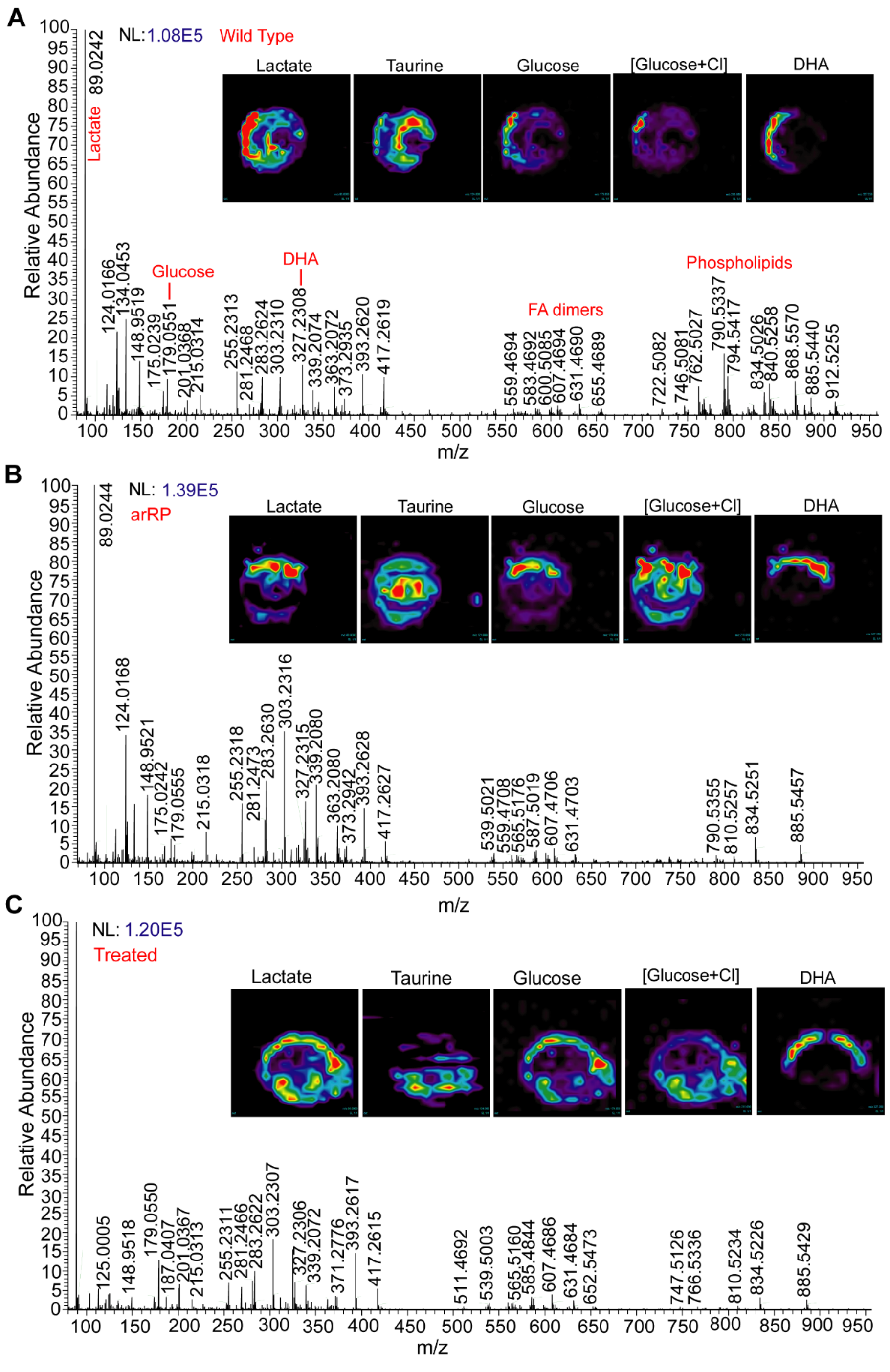

### Supplemental Figure 6

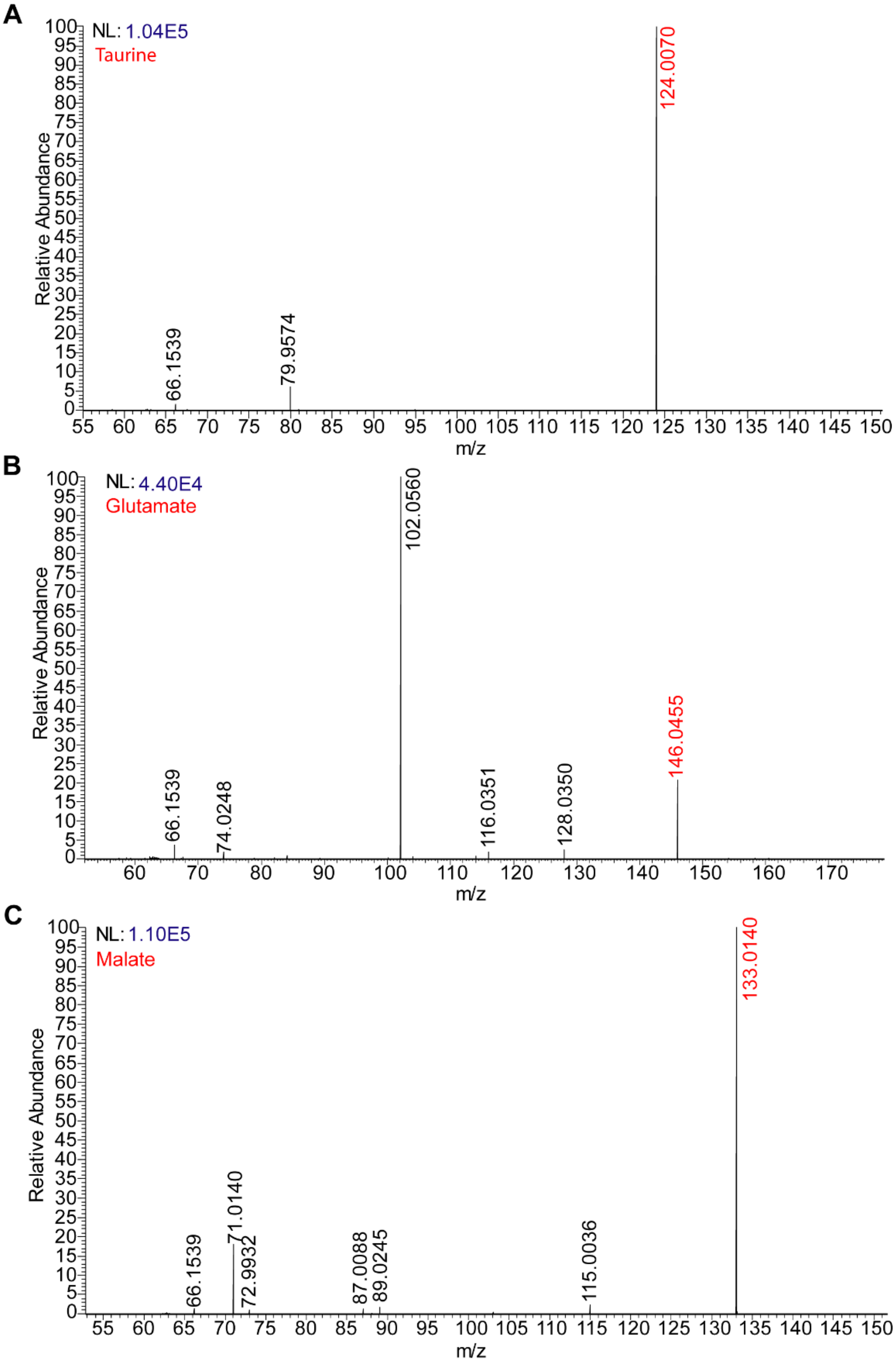

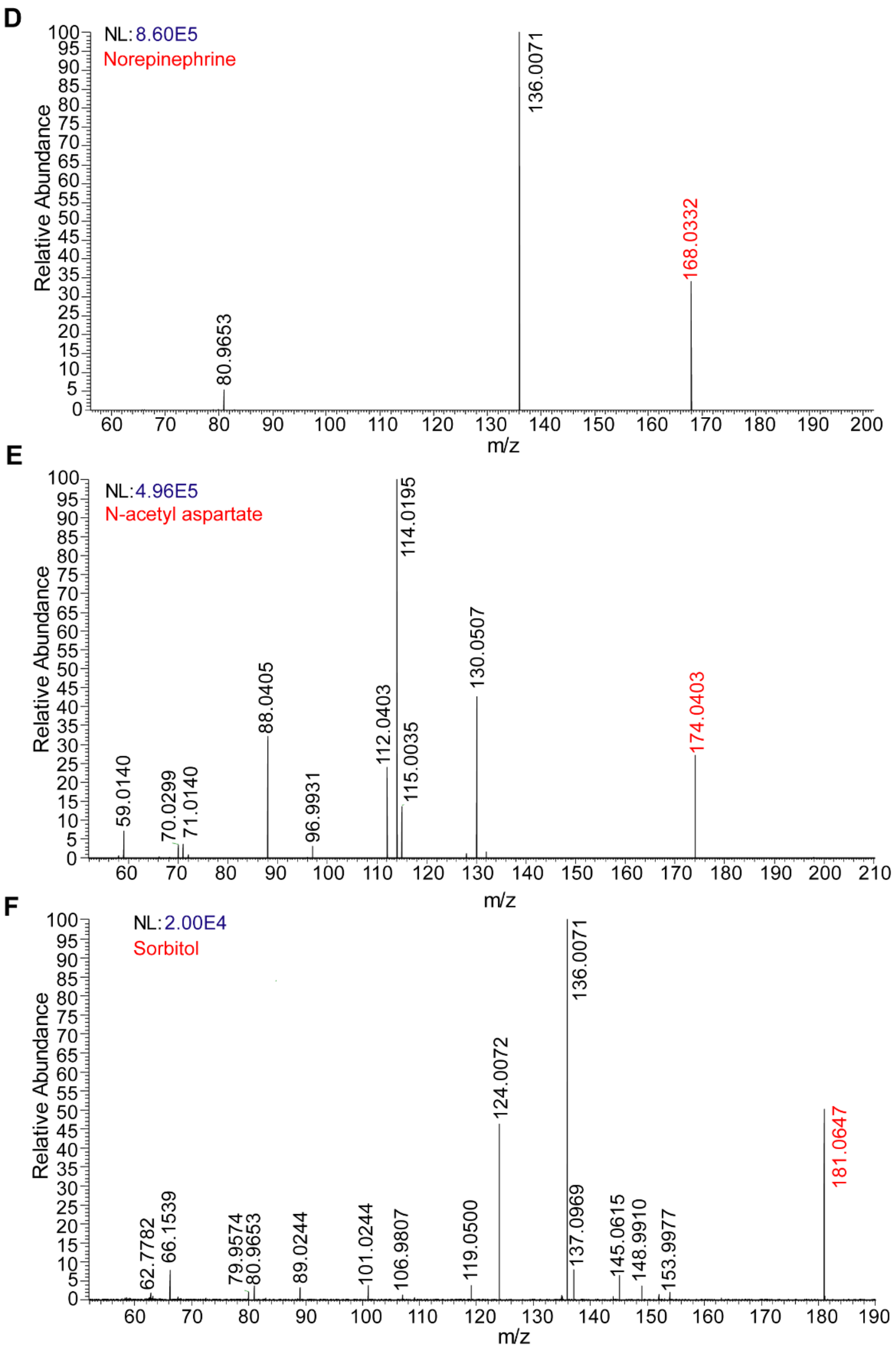

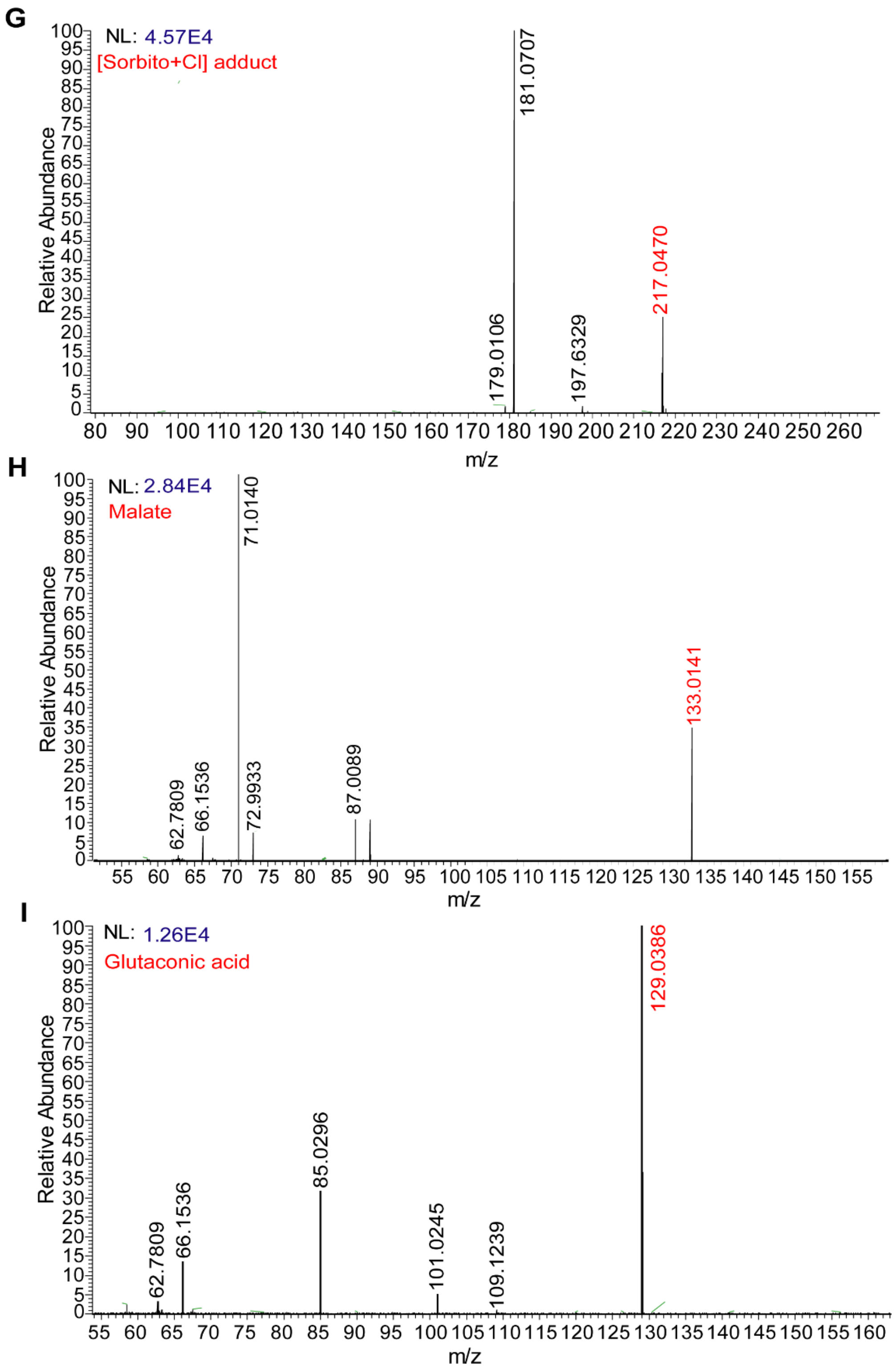

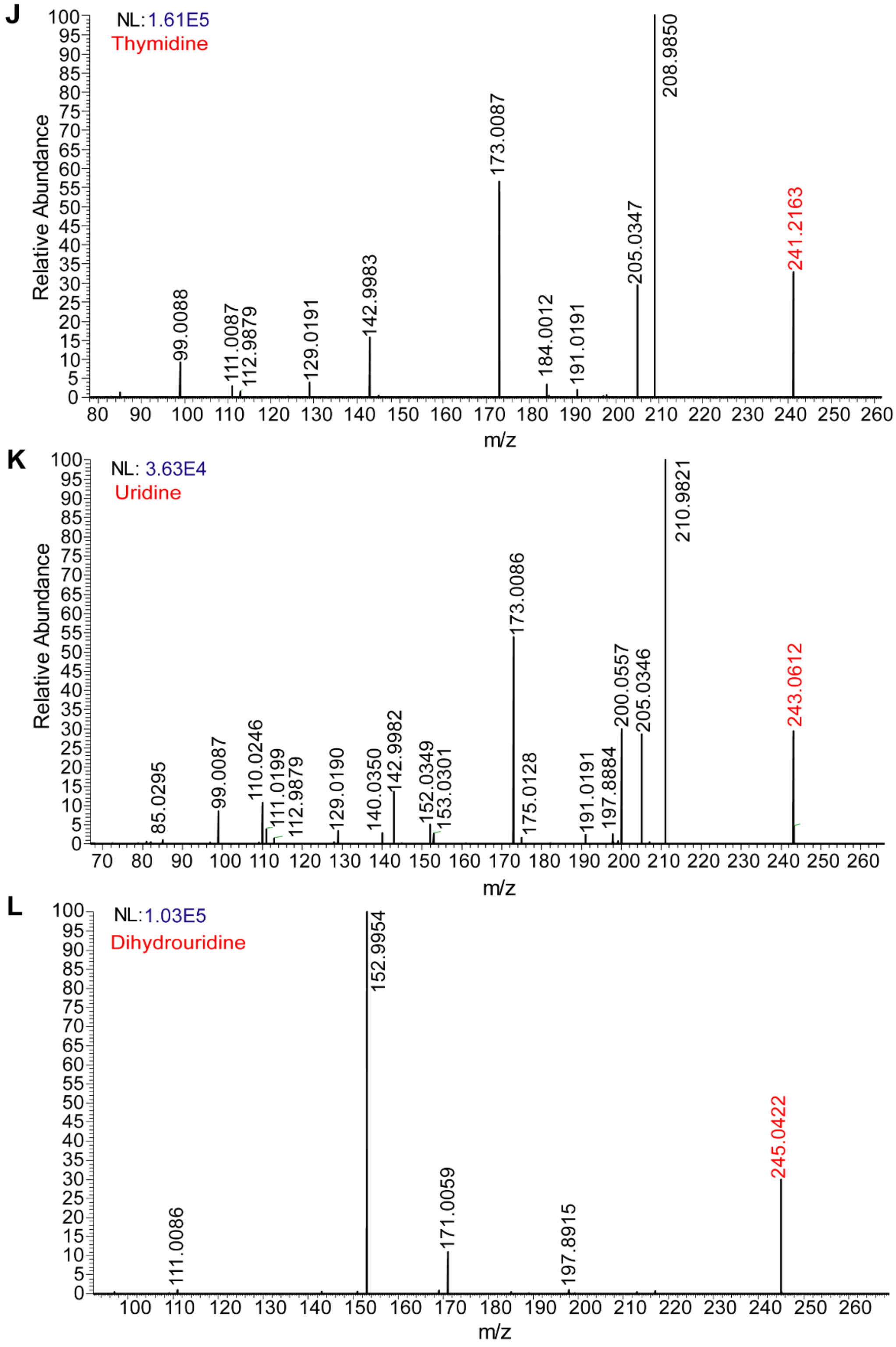

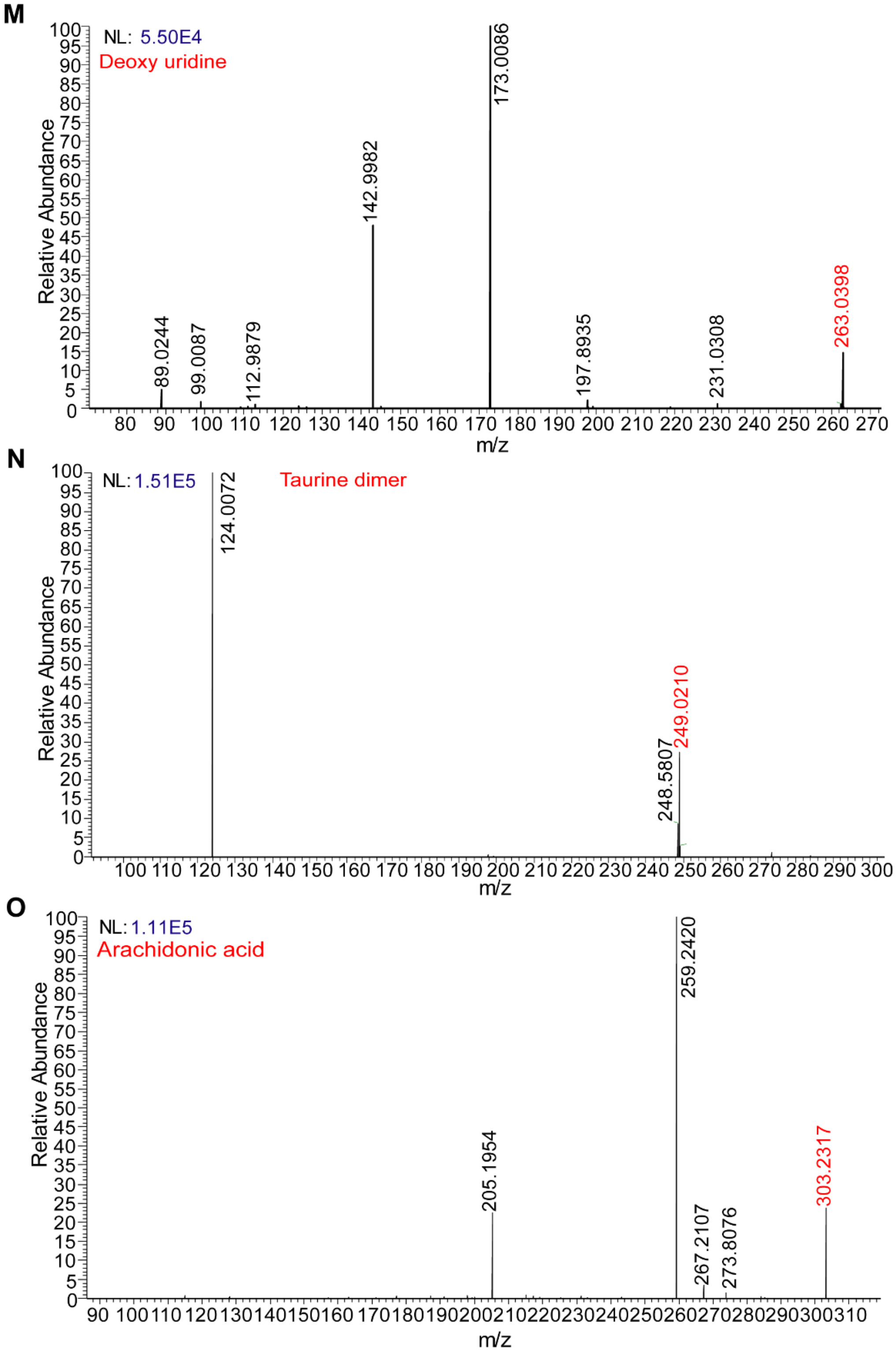

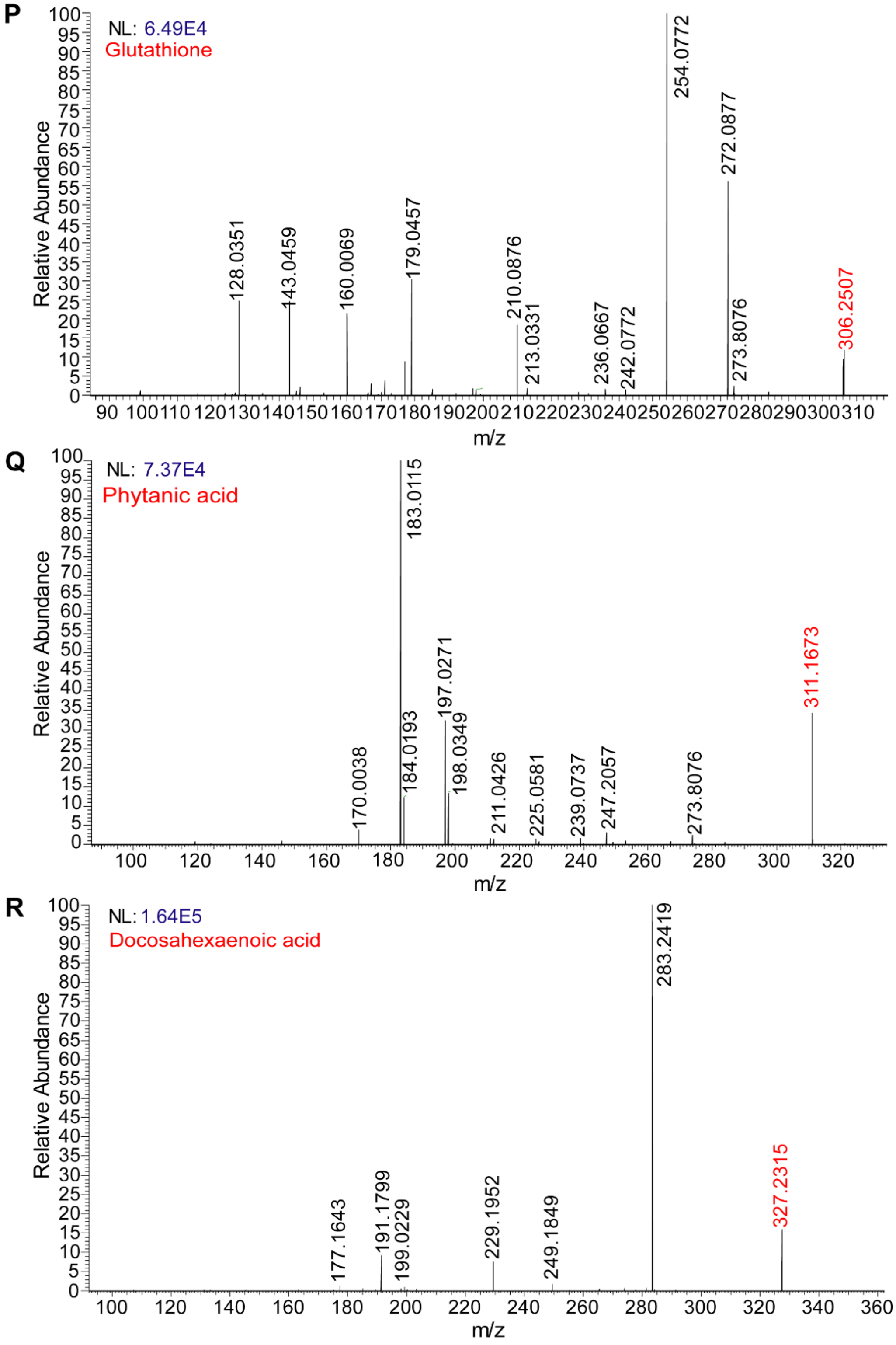

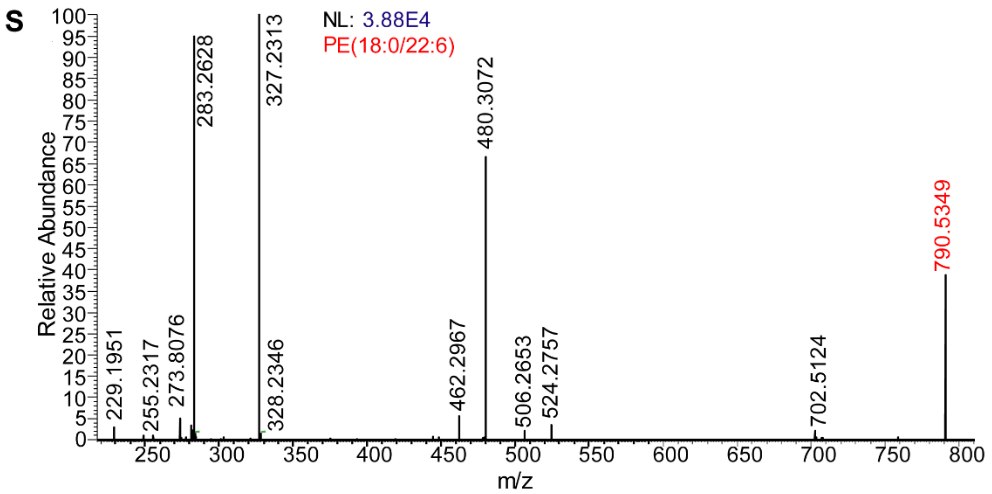

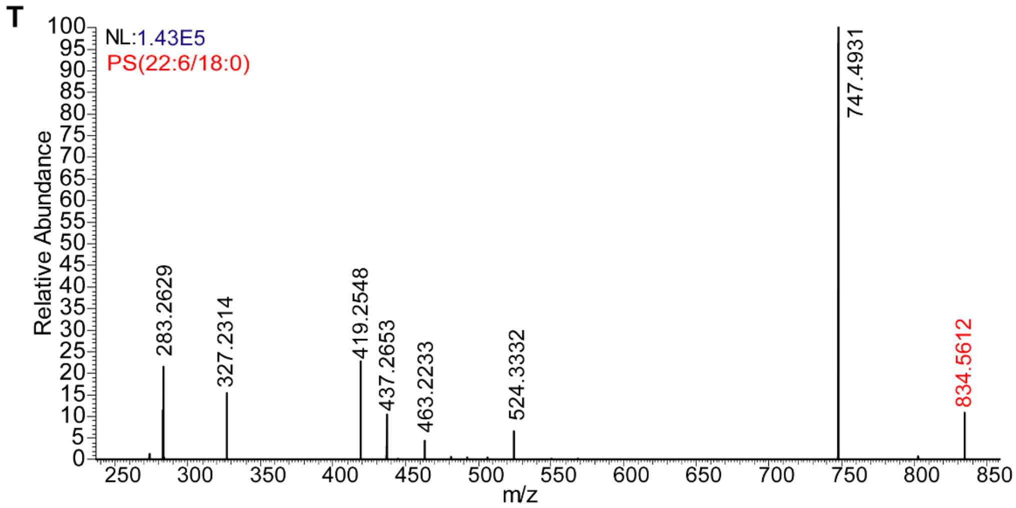

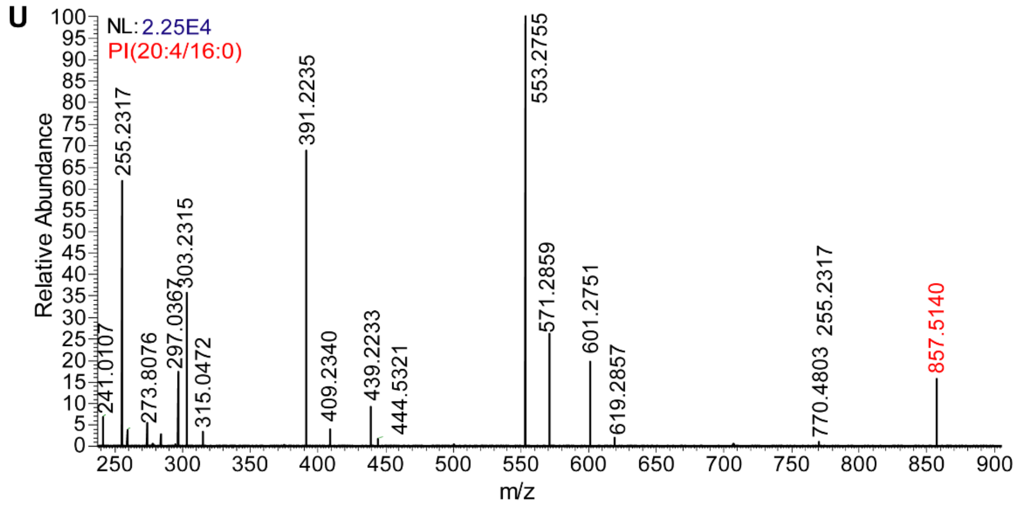

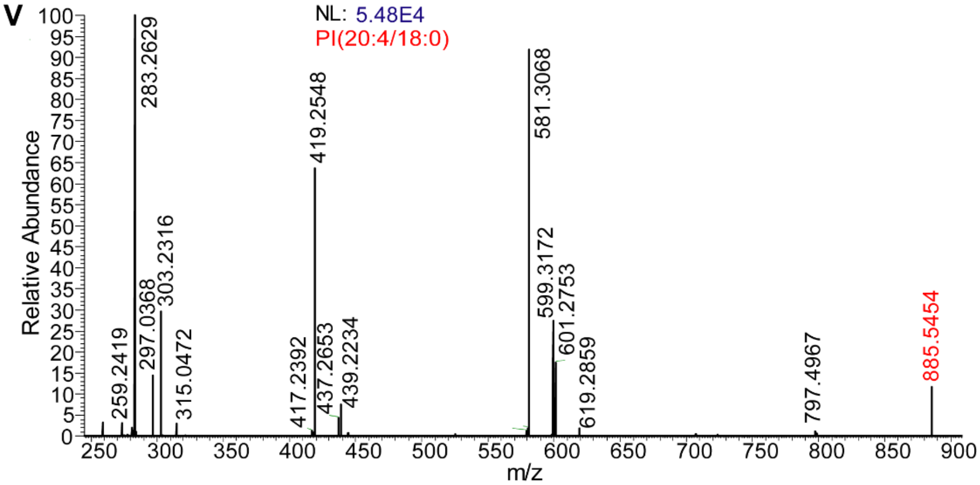

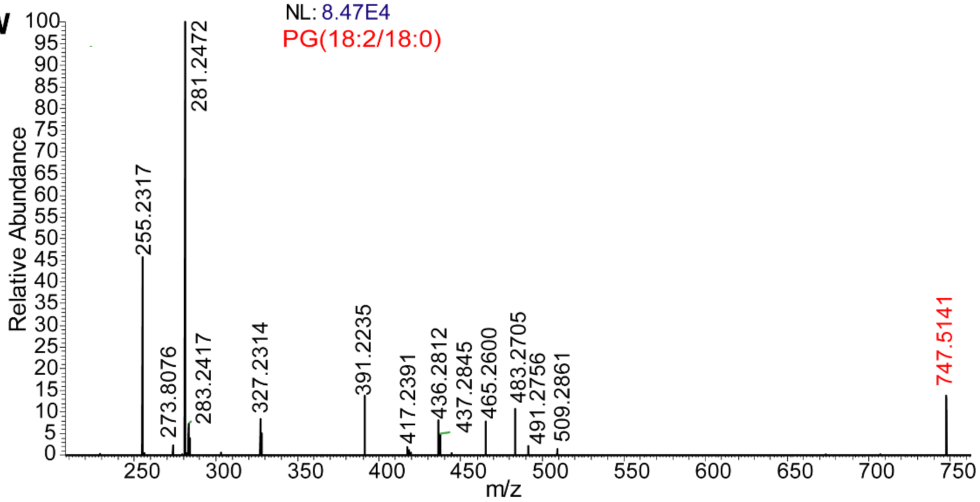

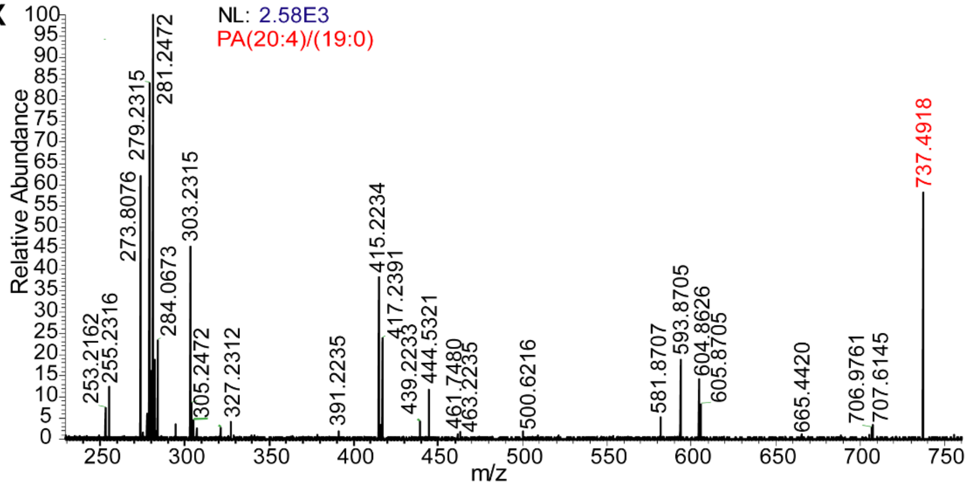

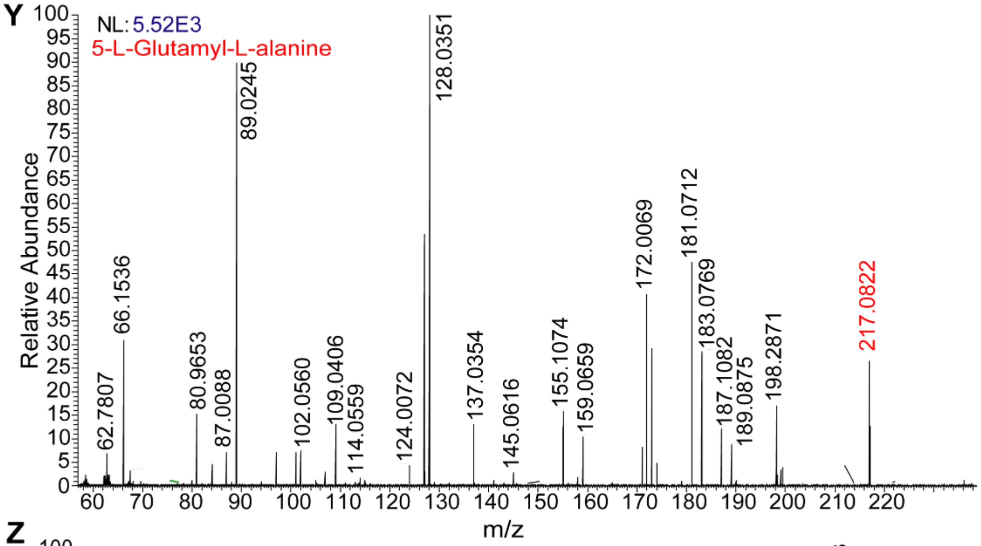

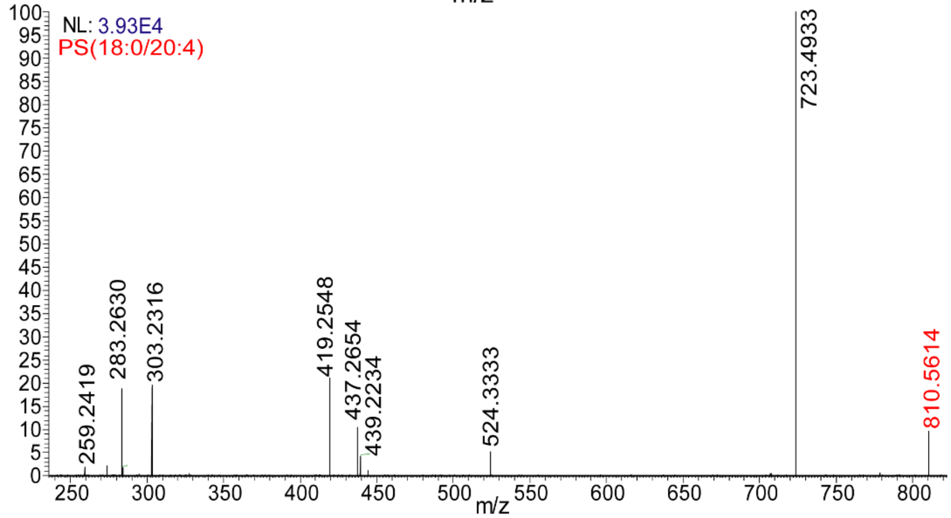

### Supplemental Figure 7

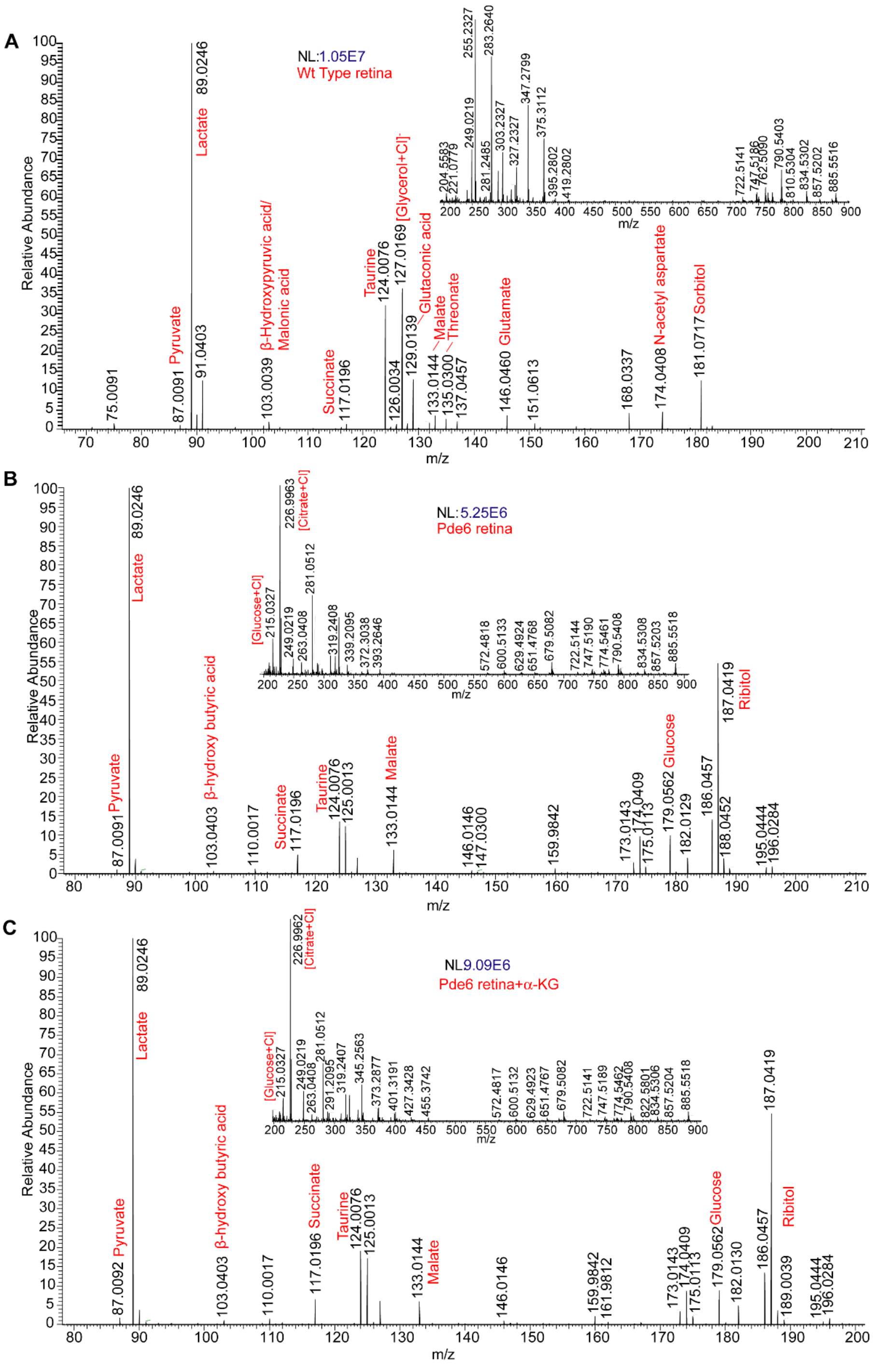

### Supplemental Figure 8

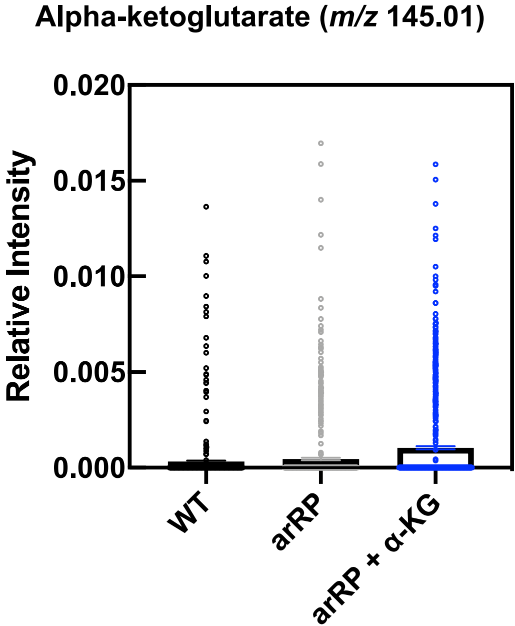
