## Supplemental Table 1 for "Metabolite therapy guided by liquid biopsy proteomics delays retinal neurodegeneration"

**Table S1.** **Vitreous biopsies from RP patients and unaffected controls:** RP, retinitis pigmentosa; ERM, epiretinal membrane.

| **Retinitis Pigmentosa Samples** | | | | | |
| --- | --- | --- | --- | --- | --- |
| **Patient** | **Sex** | **Age*** | **Eye** | **Surgical Indication** | **Diagnosis** |
| 1 | M | 38 | OD | Vitrectomy, membrane peel | RP/ERM |
| 2 | M | 41 | OS | Vitrectomy, membrane peel | RP/ERM |
| **Control Samples** | | | | | |
| **Patient** | **Sex** | **Age*** | **Eye** | **Surgical Indication** | **Diagnosis** |
| 3 | M | 81 | OD | Vitrectomy, membrane peel | ERM |
| 4 | F | 65 | OD | Vitrectomy, membrane peel | ERM |

** - At the time of surgery*
