## Supplemental Table 2 for "Metabolite therapy guided by liquid biopsy proteomics delays retinal neurodegeneration"

**Table S2.** **Proteomic content of human vitreous samples by LC-MS/MS analysis:** RP, retinitis pigmentosa; ERM, epiretinal membrane.

| **Patient** | **Condition** | **Peptide Hits** | **Peptides** | **Proteins** | **Clusters** | **Unique Proteins** |
| --- | --- | --- | --- | --- | --- | --- |
| 1 | RP/ERM | 68,329 | 1,862 | 3,255 | 841 | 875 |
| 2 | RP/ERM | 75,336 | 1,481 | 733 | 181 | 215 |
| 3 | ERM | 114,166 | 1,627 | 738 | 178 | 352 |
| 4 | ERM | 108,540 | 1,717 | 891 | 233 | 286 |
