## Supplemental Table 4 for "Metabolite therapy guided by liquid biopsy proteomics delays retinal neurodegeneration"

**Table S4.** Proteomic content of *Pde6ɑ^D670G^* retina and vitreous by LC-MS/MS analysis.

| **SAMPLE** | **MS File Name (Derived)** | **Protein Groups** | **Distinguishable Proteins** | **Distinct Peptides** | **SC (Weighted)** |
| --- | --- | --- | --- | --- | --- |
| s243_r1 | Mouse_Retina_Control_t90_s243_r1_f1 | 432 | 546 | 1331 | 8258.16 |
| s243_r1 | Mouse_Retina_Control_t90_s243_r1_f2 | 415 | 533 | 1252 | 8522.72 |
| s243_r1 | Mouse_Retina_Control_t90_s243_r1_f3 | 412 | 525 | 1194 | 7247.45 |
| s243_r1 | Mouse_Retina_Control_t90_s243_r1_f4 | 379 | 484 | 1105 | 6276.24 |
| s243_r1 | Mouse_Retina_Control_t90_s243_r1_f5 | 373 | 468 | 1061 | 5443.58 |
| s243_r1 | Mouse_Retina_Control_t90_s243_r1_f6 | 416 | 518 | 1039 | 5023.47 |
| s243_r2 | Mouse_Retina_Control_t90_s243_r2_f1 | 412 | 532 | 1285 | 8336.49 |
| s243_r2 | Mouse_Retina_Control_t90_s243_r2_f2 | 430 | 537 | 1264 | 8356.77 |
| s243_r2 | Mouse_Retina_Control_t90_s243_r2_f3 | 402 | 513 | 1201 | 7261.95 |
| s243_r2 | Mouse_Retina_Control_t90_s243_r2_f4 | 356 | 458 | 1082 | 6168.93 |
| s243_r2 | Mouse_Retina_Control_t90_s243_r2_f5 | 356 | 455 | 1044 | 5498.02 |
| s243_r2 | Mouse_Retina_Control_t90_s243_r2_f6 | 414 | 504 | 1039 | 5122.21 |
| s243_r3 | Mouse_Retina_Control_t90_s243_r3_f1 | 431 | 543 | 1307 | 8474.75 |
| s243_r3 | Mouse_Retina_Control_t90_s243_r3_f2 | 413 | 521 | 1245 | 8607.75 |
| s243_r3 | Mouse_Retina_Control_t90_s243_r3_f3 | 401 | 509 | 1159 | 7140.88 |
| s243_r3 | Mouse_Retina_Control_t90_s243_r3_f4 | 354 | 459 | 1030 | 5947.46 |
| s243_r3 | Mouse_Retina_Control_t90_s243_r3_f5 | 343 | 435 | 976 | 5337.57 |
| s243_r3 | Mouse_Retina_Control_t90_s243_r3_f6 | 383 | 469 | 984 | 5012.04 |
| s244_r1 | Mouse_Vitreous_Control_t90_s244_r1_f1 | 102 | 151 | 410 | 5788.03 |
| s244_r1 | Mouse_Vitreous_Control_t90_s244_r1_f2 | 89 | 128 | 399 | 4321.83 |
| s244_r1 | Mouse_Vitreous_Control_t90_s244_r1_f3 | 83 | 116 | 396 | 3982.39 |
| s244_r1 | Mouse_Vitreous_Control_t90_s244_r1_f4 | 84 | 114 | 352 | 3046.66 |
| s244_r1 | Mouse_Vitreous_Control_t90_s244_r1_f5 | 68 | 95 | 290 | 2445.29 |
| s244_r1 | Mouse_Vitreous_Control_t90_s244_r1_f6 | 85 | 122 | 318 | 2482.25 |
| s244_r2 | Mouse_Vitreous_Control_t90_s244_r2_f1 | 109 | 154 | 418 | 5774.15 |
| s244_r2 | Mouse_Vitreous_Control_t90_s244_r2_f2 | 85 | 124 | 392 | 4674.81 |
| s244_r2 | Mouse_Vitreous_Control_t90_s244_r2_f3 | 79 | 115 | 383 | 3801.49 |
| s244_r2 | Mouse_Vitreous_Control_t90_s244_r2_f4 | 74 | 97 | 348 | 3099.44 |
| s244_r2 | Mouse_Vitreous_Control_t90_s244_r2_f5 | 69 | 96 | 301 | 2548.21 |
| s244_r2 | Mouse_Vitreous_Control_t90_s244_r2_f6 | 77 | 108 | 298 | 2345.96 |
| s244_r3 | Mouse_Vitreous_Control_t90_s244_r3_f1 | 97 | 131 | 407 | 5741.26 |
| s244_r3 | Mouse_Vitreous_Control_t90_s244_r3_f2 | 91 | 130 | 383 | 4170.43 |
| s244_r3 | Mouse_Vitreous_Control_t90_s244_r3_f3 | 77 | 108 | 370 | 3515.83 |
| s244_r3 | Mouse_Vitreous_Control_t90_s244_r3_f4 | 69 | 97 | 317 | 2746.37 |
| s244_r3 | Mouse_Vitreous_Control_t90_s244_r3_f5 | 63 | 87 | 282 | 2430.59 |
| s244_r3 | Mouse_Vitreous_Control_t90_s244_r3_f6 | 80 | 107 | 293 | 2327.21 |
| s245_r1 | Mouse_Retina_PDE6_t90_s245_r1_f1 | 118 | 169 | 299 | 1345.07 |
| s245_r1 | Mouse_Retina_PDE6_t90_s245_r1_f2 | 191 | 267 | 555 | 8061.12 |
| s245_r1 | Mouse_Retina_PDE6_t90_s245_r1_f3 | 183 | 247 | 559 | 4788.87 |
| s245_r1 | Mouse_Retina_PDE6_t90_s245_r1_f4 | 165 | 224 | 526 | 4055.3 |
| s245_r1 | Mouse_Retina_PDE6_t90_s245_r1_f5 | 151 | 209 | 524 | 3813.88 |
| s245_r1 | Mouse_Retina_PDE6_t90_s245_r1_f6 | 47 | 82 | 169 | 1454.31 |
| s245_r2 | Mouse_Retina_PDE6_t90_s245_r2_f1 | 99 | 152 | 281 | 1308.81 |
| s245_r2 | Mouse_Retina_PDE6_t90_s245_r2_f2 | 203 | 281 | 585 | 7668.05 |
| s245_r2 | Mouse_Retina_PDE6_t90_s245_r2_f3 | 188 | 257 | 567 | 4826.06 |
| s245_r2 | Mouse_Retina_PDE6_t90_s245_r2_f4 | 155 | 211 | 510 | 4097.77 |
| s245_r2 | Mouse_Retina_PDE6_t90_s245_r2_f5 | 153 | 208 | 486 | 3651.86 |
| s245_r2 | Mouse_Retina_PDE6_t90_s245_r2_f6 | 162 | 215 | 466 | 3188.96 |
| s245_r3 | Mouse_Retina_PDE6_t90_s245_r3_f1 | 106 | 162 | 287 | 1387.03 |
| s245_r3 | Mouse_Retina_PDE6_t90_s245_r3_f2 | 202 | 279 | 589 | 8467.23 |
| s245_r3 | Mouse_Retina_PDE6_t90_s245_r3_f3 | 159 | 213 | 521 | 4187.68 |
| s245_r3 | Mouse_Retina_PDE6_t90_s245_r3_f4 | 152 | 212 | 491 | 3577.82 |
| s245_r3 | Mouse_Retina_PDE6_t90_s245_r3_f5 | 156 | 208 | 468 | 2887.93 |
| s245_r3 | Mouse_Retina_PDE6_t90_s245_r3_f6 | 251 | 330 | 863 | 8678.03 |
| s246_r1 | Mouse_Vitreous_PDE6_t90_s246_r1_f1 | 284 | 367 | 987 | 9717.2 |
| s246_r1 | Mouse_Vitreous_PDE6_t90_s246_r1_f2 | 198 | 270 | 790 | 8266.55 |
| s246_r1 | Mouse_Vitreous_PDE6_t90_s246_r1_f3 | 165 | 235 | 698 | 7140.18 |
| s246_r1 | Mouse_Vitreous_PDE6_t90_s246_r1_f4 | 153 | 221 | 664 | 6248.8 |
| s246_r1 | Mouse_Vitreous_PDE6_t90_s246_r1_f5 | 166 | 228 | 653 | 5875.04 |
| s246_r1 | Mouse_Vitreous_PDE6_t90_s246_r1_f6 | 224 | 300 | 733 | 5361.75 |
| s246_r2 | Mouse_Vitreous_PDE6_t90_s246_r2_f1 | 279 | 361 | 970 | 9321.81 |
| s246_r2 | Mouse_Vitreous_PDE6_t90_s246_r2_f2 | 183 | 255 | 775 | 7940.1 |
| s246_r2 | Mouse_Vitreous_PDE6_t90_s246_r2_f3 | 163 | 240 | 692 | 6947.22 |
| s246_r2 | Mouse_Vitreous_PDE6_t90_s246_r2_f4 | 161 | 230 | 663 | 6322.05 |
| s246_r2 | Mouse_Vitreous_PDE6_t90_s246_r2_f5 | 168 | 241 | 650 | 5777.32 |
| s246_r2 | Mouse_Vitreous_PDE6_t90_s246_r2_f6 | 213 | 285 | 699 | 5601.77 |
| s246_r3 | Mouse_Vitreous_PDE6_t90_s246_r3_f1 | 211 | 290 | 832 | 7829.12 |
| s246_r3 | Mouse_Vitreous_PDE6_t90_s246_r3_f2 | 168 | 240 | 679 | 6523.42 |
| s246_r3 | Mouse_Vitreous_PDE6_t90_s246_r3_f3 | 151 | 221 | 621 | 5874.75 |
| s246_r3 | Mouse_Vitreous_PDE6_t90_s246_r3_f4 | 167 | 229 | 643 | 5682.69 |
| s246_r3 | Mouse_Vitreous_PDE6_t90_s246_r3_f5 | 218 | 285 | 699 | 5360.5 |
| s246_r3 | Mouse_Vitreous_PDE6_t90_s246_r3_f6 | 42 | 74 | 221 | 2162.19 |
| s264_r1 | Mouse_Retina_PDE6_t28_s264_r1_f1 | 504 | 634 | 1487 | 4131.98 |
| s264_r1 | Mouse_Retina_PDE6_t28_s264_r1_f2 | 349 | 452 | 960 | 2504.43 |
| s264_r1 | Mouse_Retina_PDE6_t28_s264_r1_f3 | 241 | 321 | 590 | 1413.46 |
| s264_r1 | Mouse_Retina_PDE6_t28_s264_r1_f4 | 285 | 367 | 687 | 1787.06 |
| s264_r1 | Mouse_Retina_PDE6_t28_s264_r1_f5 | 144 | 205 | 313 | 666.04 |
| s264_r1 | Mouse_Retina_PDE6_t28_s264_r1_f6 | 301 | 378 | 710 | 1901.81 |
| s264_r2 | Mouse_Retina_PDE6_t28_s264_r3_f1 | 515 | 652 | 1560 | 4070.64 |
| s264_r2 | Mouse_Retina_PDE6_t28_s264_r3_f2 | 374 | 488 | 1092 | 2576.71 |
| s264_r2 | Mouse_Retina_PDE6_t28_s264_r3_f3 | 300 | 392 | 793 | 1849.55 |
| s264_r2 | Mouse_Retina_PDE6_t28_s264_r3_f4 | 297 | 385 | 749 | 1611.02 |
| s264_r2 | Mouse_Retina_PDE6_t28_s264_r3_f5 | 158 | 219 | 388 | 765.51 |
| s264_r2 | Mouse_Retina_PDE6_t28_s264_r3_f6 | 338 | 427 | 933 | 2261.34 |
| s264_r3 | Mouse_Retina_PDE6_t28_s264_r3_f1 | 515 | 652 | 1560 | 4070.64 |
| s264_r3 | Mouse_Retina_PDE6_t28_s264_r3_f2 | 374 | 488 | 1092 | 2576.71 |
| s264_r3 | Mouse_Retina_PDE6_t28_s264_r3_f3 | 300 | 392 | 793 | 1849.55 |
| s264_r3 | Mouse_Retina_PDE6_t28_s264_r3_f4 | 297 | 385 | 749 | 1611.02 |
| s264_r3 | Mouse_Retina_PDE6_t28_s264_r3_f5 | 158 | 219 | 388 | 765.51 |
| s264_r3 | Mouse_Retina_PDE6_t28_s264_r3_f6 | 338 | 427 | 933 | 2261.34 |
| s265_r1 | Mouse_Vitreous_PDE6_t28_s265_r1_f1 | 75 | 119 | 191 | 525.44 |
| s265_r1 | Mouse_Vitreous_PDE6_t28_s265_r1_f2 | 45 | 76 | 130 | 388.02 |
| s265_r1 | Mouse_Vitreous_PDE6_t28_s265_r1_f3 | 39 | 61 | 91 | 294.63 |
| s265_r1 | Mouse_Vitreous_PDE6_t28_s265_r1_f4 | 18 | 29 | 28 | 60.79 |
| s265_r1 | Mouse_Vitreous_PDE6_t28_s265_r1_f5 | 17 | 30 | 23 | 29.79 |
| s265_r1 | Mouse_Vitreous_PDE6_t28_s265_r1_f6 | 28 | 47 | 58 | 146.98 |
| s265_r2 | Mouse_Vitreous_PDE6_t28_s265_r2_f1 | 42 | 75 | 126 | 340.18 |
| s265_r2 | Mouse_Vitreous_PDE6_t28_s265_r2_f2 | 43 | 70 | 123 | 342.63 |
| s265_r2 | Mouse_Vitreous_PDE6_t28_s265_r2_f3 | 42 | 64 | 108 | 278.6 |
| s265_r2 | Mouse_Vitreous_PDE6_t28_s265_r2_f4 | 22 | 37 | 40 | 74.71 |
| s265_r2 | Mouse_Vitreous_PDE6_t28_s265_r2_f5 | 17 | 30 | 37 | 58.42 |
| s265_r2 | Mouse_Vitreous_PDE6_t28_s265_r2_f6 | 32 | 51 | 74 | 201 |
| s265_r3 | Mouse_Vitreous_PDE6_t28_s265_r3_f1 | 46 | 83 | 119 | 300.08 |
| s265_r3 | Mouse_Vitreous_PDE6_t28_s265_r3_f2 | 39 | 64 | 111 | 288.3 |
| s265_r3 | Mouse_Vitreous_PDE6_t28_s265_r3_f3 | 38 | 63 | 103 | 252.27 |
| s265_r3 | Mouse_Vitreous_PDE6_t28_s265_r3_f4 | 19 | 37 | 35 | 69.72 |
| s265_r3 | Mouse_Vitreous_PDE6_t28_s265_r3_f5 | 19 | 31 | 34 | 51.39 |
| s265_r3 | Mouse_Vitreous_PDE6_t28_s265_r3_f6 | 27 | 41 | 66 | 190.95 |
| s266_r1 | Mouse_Retina_PDE6_t15_s266_r1_f1 | 690 | 815 | 1716 | 10083.33 |
| s266_r1 | Mouse_Retina_PDE6_t15_s266_r1_f2 | 480 | 592 | 1457 | 7597.77 |
| s266_r1 | Mouse_Retina_PDE6_t15_s266_r1_f3 | 392 | 492 | 1167 | 5129.16 |
| s266_r1 | Mouse_Retina_PDE6_t15_s266_r1_f4 | 293 | 377 | 881 | 3480.44 |
| s266_r1 | Mouse_Retina_PDE6_t15_s266_r1_f5 | 264 | 342 | 710 | 2559.83 |
| s266_r1 | Mouse_Retina_PDE6_t15_s266_r1_f6 | 279 | 367 | 677 | 2232.58 |
| s266_r2 | Mouse_Retina_PDE6_t15_s266_r2_f1 | 739 | 860 | 1866 | 10507.15 |
| s266_r2 | Mouse_Retina_PDE6_t15_s266_r2_f2 | 545 | 665 | 1623 | 8107.64 |
| s266_r2 | Mouse_Retina_PDE6_t15_s266_r2_f3 | 454 | 557 | 1266 | 5316.71 |
| s266_r2 | Mouse_Retina_PDE6_t15_s266_r2_f4 | 340 | 435 | 972 | 3401.87 |
| s266_r2 | Mouse_Retina_PDE6_t15_s266_r2_f5 | 250 | 322 | 667 | 2282.58 |
| s266_r2 | Mouse_Retina_PDE6_t15_s266_r2_f6 | 300 | 389 | 687 | 2089.96 |
| s266_r3 | Mouse_Retina_PDE6_t15_s266_r3_f1 | 748 | 874 | 1867 | 10831.81 |
| s266_r3 | Mouse_Retina_PDE6_t15_s266_r3_f2 | 541 | 661 | 1600 | 8250.13 |
| s266_r3 | Mouse_Retina_PDE6_t15_s266_r3_f3 | 452 | 560 | 1211 | 5007.02 |
| s266_r3 | Mouse_Retina_PDE6_t15_s266_r3_f4 | 314 | 401 | 824 | 2780.83 |
| s266_r3 | Mouse_Retina_PDE6_t15_s266_r3_f5 | 231 | 302 | 588 | 1929.78 |
| s266_r3 | Mouse_Retina_PDE6_t15_s266_r3_f6 | 320 | 406 | 732 | 2173.16 |
| s267_r1 | Mouse_Vitreous_PDE6_t15_s267_r1_f1 | 230 | 298 | 737 | 4423.85 |
| s267_r1 | Mouse_Vitreous_PDE6_t15_s267_r1_f2 | 172 | 236 | 560 | 3148.55 |
| s267_r1 | Mouse_Vitreous_PDE6_t15_s267_r1_f3 | 128 | 178 | 408 | 1849.33 |
| s267_r1 | Mouse_Vitreous_PDE6_t15_s267_r1_f4 | 125 | 174 | 361 | 1563.12 |
| s267_r1 | Mouse_Vitreous_PDE6_t15_s267_r1_f5 | 81 | 119 | 275 | 1070.58 |
| s267_r1 | Mouse_Vitreous_PDE6_t15_s267_r1_f6 | 98 | 134 | 289 | 2964.05 |
| s267_r2 | Mouse_Vitreous_PDE6_t15_s267_r2_f1 | 230 | 302 | 724 | 4258.12 |
| s267_r2 | Mouse_Vitreous_PDE6_t15_s267_r2_f2 | 157 | 218 | 526 | 2852.52 |
| s267_r2 | Mouse_Vitreous_PDE6_t15_s267_r2_f3 | 118 | 167 | 381 | 1679.34 |
| s267_r2 | Mouse_Vitreous_PDE6_t15_s267_r2_f4 | 106 | 149 | 349 | 1428.54 |
| s267_r2 | Mouse_Vitreous_PDE6_t15_s267_r2_f5 | 67 | 96 | 237 | 781.67 |
| s267_r2 | Mouse_Vitreous_PDE6_t15_s267_r2_f6 | 96 | 125 | 269 | 2692.73 |
| s267_r3 | Mouse_Vitreous_PDE6_t15_s267_r3_f1 | 189 | 255 | 588 | 3220.8 |
| s267_r3 | Mouse_Vitreous_PDE6_t15_s267_r3_f2 | 116 | 169 | 371 | 1709.76 |
| s267_r3 | Mouse_Vitreous_PDE6_t15_s267_r3_f3 | 89 | 135 | 268 | 1023.79 |
| s267_r3 | Mouse_Vitreous_PDE6_t15_s267_r3_f4 | 83 | 119 | 255 | 916.74 |
| s267_r3 | Mouse_Vitreous_PDE6_t15_s267_r3_f5 | 73 | 105 | 244 | 860.75 |
| s267_r3 | Mouse_Vitreous_PDE6_t15_s267_r3_f6 | 91 | 122 | 258 | 2533.78 |
