## Supplemental Table 11 for "Metabolite therapy guided by liquid biopsy proteomics delays retinal neurodegeneration"

**Table S10.** Down-regulated proteins in the *Pde6ɑ^D670G^* retina.

| **UniProt** | **Symbol** | **Protein names** |
| --- | --- | --- |
| Q6ZPJ3 | UBE2O | (E3-independent) E2 ubiquitin-conjugating enzyme UBE2O (EC 2.3.2.24) (E2/E3 hybrid ubiquitin-protein ligase UBE2O) (Ubiquitin carrier protein O) (Ubiquitin-conjugating enzyme E2 O) (Ubiquitin-conjugating enzyme E2 of 230 kDa) (Ubiquitin-conjugating enzyme E2-230K) (Ubiquitin-protein ligase O) |
| Q9D517 | PLCC | 1-acyl-sn-glycerol-3-phosphate acyltransferase gamma (EC 2.3.1.51) (1-acylglycerol-3-phosphate O-acyltransferase 3) (1-AGP acyltransferase 3) (1-AGPAT 3) (Lysophosphatidic acid acyltransferase gamma) (LPAAT-gamma) |
| O08810 | U5S1 | 116 kDa U5 small nuclear ribonucleoprotein component (Elongation factor Tu GTP-binding domain-containing protein 2) (U5 snRNP-specific protein, 116 kDa) (U5-116 kDa) |
| P62259 | 1433E | 14-3-3 protein epsilon (14-3-3E) |
| Q60597 | ODO1 | 2-oxoglutarate dehydrogenase, mitochondrial (EC 1.2.4.2) (2-oxoglutarate dehydrogenase complex component E1) (OGDC-E1) (Alpha-ketoglutarate dehydrogenase) |
| Q9D8W5 | PSD12 | 26S proteasome non-ATPase regulatory subunit 12 (26S proteasome regulatory subunit RPN5) (26S proteasome regulatory subunit p55) |
| Q8VDM4 | PSMD2 | 26S proteasome non-ATPase regulatory subunit 2 (26S proteasome regulatory subunit RPN1) (26S proteasome regulatory subunit S2) (26S proteasome subunit p97) |
| A0ZNJ2 | A0ZNJ2 | 3-hydroxyisobutyrate dehydrogenase (HIBADH) (EC 1.1.1.31) |
| P61922 | GABT | 4-aminobutyrate aminotransferase, mitochondrial (EC 2.6.1.19) ((S)-3-amino-2-methylpropionate transaminase) (EC 2.6.1.22) (GABA aminotransferase) (GABA-AT) (Gamma-amino-N-butyrate transaminase) (GABA transaminase) (GABA-T) (L-AIBAT) |
| P63325 | RS10 | 40S ribosomal protein S10 |
| P14131 | RS16 | 40S ribosomal protein S16 |
| P62270 | RS18 | 40S ribosomal protein S18 (Ke-3) (Ke3) |
| Q9CZX8 | RS19 | 40S ribosomal protein S19 |
| P25444 | RS2 | 40S ribosomal protein S2 (40S ribosomal protein S4) (Protein LLRep3) |
| P62267 | RS23 | 40S ribosomal protein S23 |
| P62849 | RS24 | 40S ribosomal protein S24 |
| P62274 | RS29 | 40S ribosomal protein S29 |
| P62702 | RS4X | 40S ribosomal protein S4, X isoform |
| D3YYM6 | D3YYM6 | 40S ribosomal protein S5 (Fragment) |
| P62754 | RS6 | 40S ribosomal protein S6 (Phosphoprotein NP33) |
| P62082 | RS7 | 40S ribosomal protein S7 |
| Q6ZWN5 | RS9 | 40S ribosomal protein S9 |
| P14206 | RSSA | 40S ribosomal protein SA (37 kDa laminin receptor precursor) (37LRP) (37 kDa oncofetal antigen) (37/67 kDa laminin receptor) (LRP/LR) (67 kDa laminin receptor) (67LR) (Laminin receptor 1) (LamR) (Laminin-binding protein precursor p40) (LBP/p40) (OFA/iLRP) |
| P10852 | 4F2 | 4F2 cell-surface antigen heavy chain (4F2hc) (Solute carrier family 3 member 2) (CD antigen CD98) |
| Q9DCD0 | 6PGD | 6-phosphogluconate dehydrogenase, decarboxylating (EC 1.1.1.44) |
| Q9CQ60 | 6PGL | 6-phosphogluconolactonase (6PGL) (EC 3.1.1.31) |
| P63038 | CH60 | 60 kDa heat shock protein, mitochondrial (EC 5.6.1.7) (60 kDa chaperonin) (Chaperonin 60) (CPN60) (HSP-65) (Heat shock protein 60) (HSP-60) (Hsp60) (Mitochondrial matrix protein P1) |
| P14869 | RLA0 | 60S acidic ribosomal protein P0 (60S ribosomal protein L10E) |
| P47955 | RLA1 | 60S acidic ribosomal protein P1 |
| P99027 | RLA2 | 60S acidic ribosomal protein P2 |
| Q6ZWV3 | RL10 | 60S ribosomal protein L10 (Protein QM homolog) (Ribosomal protein L10) |
| Q9CXW4 | RL11 | 60S ribosomal protein L11 |
| P47963 | RL13 | 60S ribosomal protein L13 (A52) |
| Q9CR57 | RL14 | 60S ribosomal protein L14 |
| Q9CZM2 | RL15 | 60S ribosomal protein L15 |
| P35980 | RL18 | 60S ribosomal protein L18 |
| Q8BP67 | RL24 | 60S ribosomal protein L24 |
| P14115 | RL27A | 60S ribosomal protein L27a (L29) |
| Q9D8E6 | RL4 | 60S ribosomal protein L4 |
| P47962 | RL5 | 60S ribosomal protein L5 |
| P47911 | RL6 | 60S ribosomal protein L6 (TAX-responsive enhancer element-binding protein 107) (TAXREB107) |
| P14148 | RL7 | 60S ribosomal protein L7 |
| P12970 | RL7A | 60S ribosomal protein L7a (Surfeit locus protein 3) |
| P97822 | AN32E | Acidic leucine-rich nuclear phosphoprotein 32 family member E (Cerebellar postnatal development protein 1) (LANP-like protein) (LANP-L) |
| Q99KI0 | ACON | Aconitate hydratase, mitochondrial (Aconitase) (EC 4.2.1.3) (Citrate hydro-lyase) |
| P61161 | ARP2 | Actin-related protein 2 (Actin-like protein 2) |
| Q9JM76 | ARPC3 | Actin-related protein 2/3 complex subunit 3 (Arp2/3 complex 21 kDa subunit) (p21-ARC) |
| Q99JY9 | ARP3 | Actin-related protein 3 (Actin-like protein 3) |
| Q8JZN5 | ACAD9 | Acyl-CoA dehydrogenase family member 9, mitochondrial (ACAD-9) (EC 1.3.99.-) |
| P08030 | APT | Adenine phosphoribosyltransferase (APRT) (EC 2.4.2.7) |
| P50247 | SAHH | Adenosylhomocysteinase (AdoHcyase) (EC 3.3.1.1) (CUBP) (Liver copper-binding protein) (S-adenosyl-L-homocysteine hydrolase) |
| P54822 | PUR8 | Adenylosuccinate lyase (ASL) (EC 4.3.2.2) (Adenylosuccinase) (ASase) |
| P40124 | CAP1 | Adenylyl cyclase-associated protein 1 (CAP 1) |
| P48962 | ADT1 | ADP/ATP translocase 1 (ADP,ATP carrier protein 1) (ADP,ATP carrier protein, heart/skeletal muscle isoform T1) (Adenine nucleotide translocator 1) (ANT 1) (Solute carrier family 25 member 4) (mANC1) |
| Q8CG76 | ARK72 | Aflatoxin B1 aldehyde reductase member 2 (EC 1.1.1.n11) (Succinic semialdehyde reductase) (SSA reductase) |
| Q9JII6 | AK1A1 | Aldo-keto reductase family 1 member A1 (EC 1.1.1.2) (EC 1.1.1.33) (EC 1.1.1.372) (EC 1.1.1.54) (Alcohol dehydrogenase [NADP(+)]) (Aldehyde reductase) (Glucuronate reductase) (EC 1.1.1.19) (Glucuronolactone reductase) (EC 1.1.1.20) |
| Q9QYC0 | ADDA | Alpha-adducin (Erythrocyte adducin subunit alpha) |
| P61164 | ACTZ | Alpha-centractin (Centractin) (ARP1) (Actin-RPV) (Centrosome-associated actin homolog) |
| P24622 | CRYAA | Alpha-crystallin A chain |
| P23927 | CRYAB | Alpha-crystallin B chain (Alpha(B)-crystallin) (P23) |
| P17182 | ENOA | Alpha-enolase (EC 4.2.1.11) (2-phospho-D-glycerate hydro-lyase) (Enolase 1) (Non-neural enolase) (NNE) |
| Q9DB05 | SNAA | Alpha-soluble NSF attachment protein (SNAP-alpha) (N-ethylmaleimide-sensitive factor attachment protein alpha) |
| Q8BLS7 | SWAHA | Ankyrin repeat domain-containing protein SOWAHA (Ankyrin repeat domain-containing protein 43) (Protein sosondowah homolog A) |
| Q8C8R3 | ANK2 | Ankyrin-2 (ANK-2) (Ankyrin-B) (Brain ankyrin) |
| P07356 | ANXA2 | Annexin A2 (Annexin II) (Annexin-2) (Calpactin I heavy chain) (Calpactin-1 heavy chain) (Chromobindin-8) (Lipocortin II) (Placental anticoagulant protein IV) (PAP-IV) (Protein I) (p36) |
| O35643 | AP1B1 | AP-1 complex subunit beta-1 (Adaptor protein complex AP-1 subunit beta-1) (Adaptor-related protein complex 1 subunit beta-1) (Beta-1-adaptin) (Beta-adaptin 1) (Clathrin assembly protein complex 1 beta large chain) (Golgi adaptor HA1/AP1 adaptin beta subunit) |
| P17427 | AP2A2 | AP-2 complex subunit alpha-2 (100 kDa coated vesicle protein C) (Adaptor protein complex AP-2 subunit alpha-2) (Adaptor-related protein complex 2 subunit alpha-2) (Alpha-adaptin C) (Alpha2-adaptin) (Clathrin assembly protein complex 2 alpha-C large chain) (Plasma membrane adaptor HA2/AP2 adaptin alpha C subunit) |
| P16460 | ASSY | Argininosuccinate synthase (EC 6.3.4.5) (Citrulline--aspartate ligase) |
| P98203 | ARVC | Armadillo repeat protein deleted in velo-cardio-facial syndrome homolog |
| P05201 | AATC | Aspartate aminotransferase, cytoplasmic (cAspAT) (EC 2.6.1.1) (EC 2.6.1.3) (Cysteine aminotransferase, cytoplasmic) (Cysteine transaminase, cytoplasmic) (cCAT) (Glutamate oxaloacetate transaminase 1) (Transaminase A) |
| P05202 | AATM | Aspartate aminotransferase, mitochondrial (mAspAT) (EC 2.6.1.1) (EC 2.6.1.7) (Fatty acid-binding protein) (FABP-1) (Glutamate oxaloacetate transaminase 2) (Kynurenine aminotransferase 4) (Kynurenine aminotransferase IV) (Kynurenine--oxoglutarate transaminase 4) (Kynurenine--oxoglutarate transaminase IV) (Plasma membrane-associated fatty acid-binding protein) (FABPpm) (Transaminase A) |
| Q9Z2W0 | DNPEP | Aspartyl aminopeptidase (EC 3.4.11.21) |
| Q9CQQ7 | AT5F1 | ATP synthase F(0) complex subunit B1, mitochondrial (ATP synthase peripheral stalk-membrane subunit b) (ATP synthase subunit b) (ATPase subunit b) |
| Q78IK2 | ATPMD | ATP synthase membrane subunit DAPIT, mitochondrial (Diabetes-associated protein in insulin-sensitive tissues) (Up-regulated during skeletal muscle growth protein 5) |
| P03930 | ATP8 | ATP synthase protein 8 (A6L) (F-ATPase subunit 8) |
| Q03265 | ATPA | ATP synthase subunit alpha, mitochondrial (ATP synthase F1 subunit alpha) |
| P56480 | ATPB | ATP synthase subunit beta, mitochondrial (EC 7.1.2.2) (ATP synthase F1 subunit beta) |
| Q9DCX2 | ATP5H | ATP synthase subunit d, mitochondrial (ATPase subunit d) (ATP synthase peripheral stalk subunit d) |
| P56135 | ATPK | ATP synthase subunit f, mitochondrial (ATP synthase membrane subunit f) |
| Q91VR2 | ATPG | ATP synthase subunit gamma, mitochondrial (ATP synthase F1 subunit gamma) (F-ATPase gamma subunit) |
| Q9DB20 | ATPO | ATP synthase subunit O, mitochondrial (ATP synthase peripheral stalk subunit OSCP) (Oligomycin sensitivity conferral protein) (OSCP) |
| P55096 | ABCD3 | ATP-binding cassette sub-family D member 3 (68 kDa peroxisomal membrane protein) (PMP68) (70 kDa peroxisomal membrane protein) (PMP70) |
| P61222 | ABCE1 | ATP-binding cassette sub-family E member 1 (RNase L inhibitor) (Ribonuclease 4 inhibitor) (RNS4I) |
| Q91V92 | ACLY | ATP-citrate synthase (EC 2.3.3.8) (ATP-citrate (pro-S-)-lyase) (Citrate cleavage enzyme) |
| P12382 | PFKAL | ATP-dependent 6-phosphofructokinase, liver type (ATP-PFK) (PFK-L) (EC 2.7.1.11) (6-phosphofructokinase type B) (Phosphofructo-1-kinase isozyme B) (PFK-B) (Phosphohexokinase) |
| O70133 | DHX9 | ATP-dependent RNA helicase A (EC 3.6.4.13) (DEAH box protein 9) (mHEL-5) (Nuclear DNA helicase II) (NDH II) (RNA helicase A) (RHA) |
| Q62167 | DDX3X | ATP-dependent RNA helicase DDX3X (EC 3.6.4.13) (D1Pas1-related sequence 2) (DEAD box RNA helicase DEAD3) (mDEAD3) (DEAD box protein 3, X-chromosomal) (Embryonic RNA helicase) |
| O54984 | ASNA | ATPase Asna1 (EC 3.6.-.-) (Arsenical pump-driving ATPase) (Arsenite-stimulated ATPase) |
| Q9WV92 | E41L3 | Band 4.1-like protein 3 (4.1B) (Differentially expressed in adenocarcinoma of the lung protein 1) (DAL-1) (DAL1P) (mDAL-1) [Cleaved into: Band 4.1-like protein 3, N-terminally processed] |
| E9PZ16 | E9PZ16 | Basement membrane-specific heparan sulfate proteoglycan core protein |
| P18572 | BASI | Basigin (Basic immunoglobulin superfamily) (HT7 antigen) (Membrane glycoprotein gp42) (CD antigen CD147) |
| Q9QXC6 | Q9QXC6 | Beta-A3/A1 crystallin protein (Beta-crystallin A1) (Crystallin, beta A1) |
| Q8BFZ3 | ACTBL | Beta-actin-like protein 2 (Kappa-actin) |
| E9QAS6 | E9QAS6 | Beta-crystallin A4 |
| Q9WVJ5 | CRBB1 | Beta-crystallin B1 (Beta-B1 crystallin) [Cleaved into: Beta-crystallin B1B] |
| Q9JJU9 | CRBB3 | Beta-crystallin B3 (Beta-B3 crystallin) [Cleaved into: Beta-crystallin B3, N-terminally processed] |
| A8DUK4 | A8DUK4 | Beta-globin (Globin a1) (Hemoglobin, beta adult s chain) (Hemoglobin, beta adult t chain) |
| P28663 | SNAB | Beta-soluble NSF attachment protein (SNAP-beta) (Brain protein I47) (N-ethylmaleimide-sensitive factor attachment protein beta) |
| Q9CWJ9 | PUR9 | Bifunctional purine biosynthesis protein PURH [Includes: Phosphoribosylaminoimidazolecarboxamide formyltransferase (EC 2.1.2.3) (5-aminoimidazole-4-carboxamide ribonucleotide formyltransferase) (AICAR transformylase); IMP cyclohydrolase (EC 3.5.4.10) (ATIC) (IMP synthase) (Inosinicase)] |
| O88712 | CTBP1 | C-terminal-binding protein 1 (CtBP1) (EC 1.1.1.-) |
| P56546 | CTBP2 | C-terminal-binding protein 2 (CtBP2) |
| Q8BH59 | CMC1 | Calcium-binding mitochondrial carrier protein Aralar1 (Mitochondrial aspartate glutamate carrier 1) (Solute carrier family 25 member 12) |
| Q06138 | CAB39 | Calcium-binding protein 39 (MO25alpha) (Protein Mo25) |
| P11798 | KCC2A | Calcium/calmodulin-dependent protein kinase type II subunit alpha (CaM kinase II subunit alpha) (CaMK-II subunit alpha) (EC 2.7.11.17) |
| P35564 | CALX | Calnexin |
| O88456 | CPNS1 | Calpain small subunit 1 (CSS1) (Calcium-activated neutral proteinase small subunit) (CANP small subunit) (Calcium-dependent protease small subunit) (CDPS) (Calcium-dependent protease small subunit 1) (Calpain regulatory subunit) |
| P68181 | KAPCB | cAMP-dependent protein kinase catalytic subunit beta (PKA C-beta) (EC 2.7.11.11) |
| Q60737 | CSK21 | Casein kinase II subunit alpha (CK II alpha) (EC 2.7.11.1) |
| Q02248 | CTNB1 | Catenin beta-1 (Beta-catenin) |
| Q8R5M8 | CADM1 | Cell adhesion molecule 1 (Immunoglobulin superfamily member 4) (IgSF4) (Nectin-like protein 2) (NECL-2) (Spermatogenic immunoglobulin superfamily) (SgIgSF) (Synaptic cell adhesion molecule) (SynCAM) (Tumor suppressor in lung cancer 1) (TSLC-1) |
| Q8VDP4 | CCAR2 | Cell cycle and apoptosis regulator protein 2 (Cell division cycle and apoptosis regulator protein 2) |
| P60766 | CDC42 | Cell division control protein 42 homolog (G25K GTP-binding protein) |
| Q9CZU6 | CISY | Citrate synthase, mitochondrial (EC 2.3.3.1) (Citrate (Si)-synthase) |
| Q61548 | AP180 | Clathrin coat assembly protein AP180 (91 kDa synaptosomal-associated protein) (Clathrin coat-associated protein AP180) (Phosphoprotein F1-20) |
| Q68FD5 | CLH1 | Clathrin heavy chain 1 |
| Q8VBZ3 | CLPT1 | Cleft lip and palate transmembrane protein 1 homolog (Thymic epithelial cell surface antigen) |
| P02463 | CO4A1 | Collagen alpha-1(IV) chain [Cleaved into: Arresten] |
| P08122 | CO4A2 | Collagen alpha-2(IV) chain [Cleaved into: Canstatin] |
| O54751 | CRX | Cone-rod homeobox protein |
| O88543 | CSN3 | COP9 signalosome complex subunit 3 (SGN3) (Signalosome subunit 3) (JAB1-containing signalosome subunit 3) |
| Q9CZ04 | CSN7A | COP9 signalosome complex subunit 7a (SGN7a) (Signalosome subunit 7a) (JAB1-containing signalosome subunit 7a) |
| Q9QZQ8 | H2AY | Core histone macro-H2A.1 (Histone macroH2A1) (mH2A1) (H2A.y) (H2A/y) |
| Q04447 | KCRB | Creatine kinase B-type (EC 2.7.3.2) (B-CK) (Creatine kinase B chain) (Creatine phosphokinase B-type) (CPK-B) |
| P30275 | KCRU | Creatine kinase U-type, mitochondrial (EC 2.7.3.2) (Acidic-type mitochondrial creatine kinase) (Mia-CK) (Ubiquitous mitochondrial creatine kinase) (U-MtCK) |
| Q6ZQ38 | CAND1 | Cullin-associated NEDD8-dissociated protein 1 (Cullin-associated and neddylation-dissociated protein 1) (p120 CAND1) |
| E1AZ71 | E1AZ71 | Cyclic nucleotide-gated channel beta 1 (cGMP-gated cation channel beta subunit) |
| Q9CZ13 | QCR1 | Cytochrome b-c1 complex subunit 1, mitochondrial (Complex III subunit 1) (Core protein I) (Ubiquinol-cytochrome-c reductase complex core protein 1) |
| P00405 | COX2 | Cytochrome c oxidase subunit 2 (Cytochrome c oxidase polypeptide II) |
| P12787 | COX5A | Cytochrome c oxidase subunit 5A, mitochondrial (Cytochrome c oxidase polypeptide Va) |
| P62897 | CYC | Cytochrome c, somatic |
| Q9JHU4 | DYHC1 | Cytoplasmic dynein 1 heavy chain 1 (Cytoplasmic dynein heavy chain 1) (Dynein heavy chain, cytosolic) |
| Q8R1Q8 | DC1L1 | Cytoplasmic dynein 1 light intermediate chain 1 (Dynein light chain A) (DLC-A) (Dynein light intermediate chain 1, cytosolic) |
| Q8BMK4 | CKAP4 | Cytoskeleton-associated protein 4 (63-kDa cytoskeleton-linking membrane protein) (Climp-63) (p63) |
| Q9CPY7 | AMPL | Cytosol aminopeptidase (EC 3.4.11.1) (Leucine aminopeptidase 3) (LAP-3) (Leucyl aminopeptidase) (Proline aminopeptidase) (EC 3.4.11.5) (Prolyl aminopeptidase) |
| Q8R0Y6 | AL1L1 | Cytosolic 10-formyltetrahydrofolate dehydrogenase (10-FTHFDH) (FDH) (EC 1.5.1.6) (Aldehyde dehydrogenase family 1 member L1) |
| Q91V12 | BACH | Cytosolic acyl coenzyme A thioester hydrolase (EC 3.1.2.2) (Acyl-CoA thioesterase 7) (Brain acyl-CoA hydrolase) (BACH) (CTE-IIa) (CTE-II) (Long chain acyl-CoA thioester hydrolase) |
| Q61753 | SERA | D-3-phosphoglycerate dehydrogenase (3-PGDH) (EC 1.1.1.95) (A10) |
| O08749 | DLDH | Dihydrolipoyl dehydrogenase, mitochondrial (EC 1.8.1.4) (Dihydrolipoamide dehydrogenase) |
| Q9D2G2 | ODO2 | Dihydrolipoyllysine-residue succinyltransferase component of 2-oxoglutarate dehydrogenase complex, mitochondrial (EC 2.3.1.61) (2-oxoglutarate dehydrogenase complex component E2) (OGDC-E2) (Dihydrolipoamide succinyltransferase component of 2-oxoglutarate dehydrogenase complex) (E2K) |
| P97427 | DPYL1 | Dihydropyrimidinase-related protein 1 (DRP-1) (Collapsin response mediator protein 1) (CRMP-1) (Unc-33-like phosphoprotein 3) (ULIP-3) |
| O08553 | DPYL2 | Dihydropyrimidinase-related protein 2 (DRP-2) (Unc-33-like phosphoprotein 2) (ULIP-2) |
| Q62188 | DPYL3 | Dihydropyrimidinase-related protein 3 (DRP-3) (Unc-33-like phosphoprotein 1) (ULIP-1) |
| O35098 | DPYL4 | Dihydropyrimidinase-related protein 4 (DRP-4) (Collapsin response mediator protein 3) (CRMP-3) (UNC33-like phosphoprotein 4) (ULIP-4) |
| Q9Z218 | DPP6 | Dipeptidyl aminopeptidase-like protein 6 (DPPX) (Dipeptidyl aminopeptidase-related protein) (Dipeptidyl peptidase 6) (Dipeptidyl peptidase IV-like protein) (Dipeptidyl peptidase VI) (DPP VI) |
| Q64511 | TOP2B | DNA topoisomerase 2-beta (EC 5.6.2.3) (DNA topoisomerase II, beta isozyme) |
| P28352 | APEX1 | DNA-(apurinic or apyrimidinic site) lyase (EC 3.1.-.-) (EC 4.2.99.18) (APEX nuclease) (APEN) (Apurinic-apyrimidinic endonuclease 1) (AP endonuclease 1) (REF-1) (Redox factor-1) [Cleaved into: DNA-(apurinic or apyrimidinic site) lyase, mitochondrial] |
| O54734 | OST48 | Dolichyl-diphosphooligosaccharide--protein glycosyltransferase 48 kDa subunit (DDOST 48 kDa subunit) (Oligosaccharyl transferase 48 kDa subunit) |
| Q91YQ5 | RPN1 | Dolichyl-diphosphooligosaccharide--protein glycosyltransferase subunit 1 (Dolichyl-diphosphooligosaccharide--protein glycosyltransferase 67 kDa subunit) (Ribophorin I) (RPN-I) (Ribophorin-1) |
| Q3TDQ1 | STT3B | Dolichyl-diphosphooligosaccharide--protein glycosyltransferase subunit STT3B (Oligosaccharyl transferase subunit STT3B) (STT3-B) (EC 2.4.99.18) (B6dom1 antigen) (Source of immunodominant MHC-associated peptides) |
| P39053 | DYN1 | Dynamin-1 (EC 3.6.5.5) |
| Q8K1M6 | DNM1L | Dynamin-1-like protein (EC 3.6.5.5) (Dynamin family member proline-rich carboxyl-terminal domain less) (Dymple) (Dynamin-related protein 1) |
| Q91ZU6 | DYST | Dystonin (Bullous pemphigoid antigen 1) (BPA) (Dystonia musculorum protein) (Hemidesmosomal plaque protein) (Microtubule actin cross-linking factor 2) |
| P70372 | ELAV1 | ELAV-like protein 1 (Elav-like generic protein) (Hu-antigen R) (HuR) (MelG) |
| P10126 | EF1A1 | Elongation factor 1-alpha 1 (EF-1-alpha-1) (Elongation factor Tu) (EF-Tu) (Eukaryotic elongation factor 1 A-1) (eEF1A-1) |
| P57776 | EF1D | Elongation factor 1-delta (EF-1-delta) |
| Q9D8N0 | EF1G | Elongation factor 1-gamma (EF-1-gamma) (eEF-1B gamma) |
| P57759 | ERP29 | Endoplasmic reticulum resident protein 29 (ERp29) |
| P42125 | ECI1 | Enoyl-CoA delta isomerase 1, mitochondrial (EC 5.3.3.8) (3,2-trans-enoyl-CoA isomerase) (Delta(3),Delta(2)-enoyl-CoA isomerase) (D3,D2-enoyl-CoA isomerase) (Dodecenoyl-CoA isomerase) |
| Q8BFZ9 | ERLN2 | Erlin-2 (Endoplasmic reticulum lipid raft-associated protein 2) (Stomatin-prohibitin-flotillin-HflC/K domain-containing protein 2) (SPFH domain-containing protein 2) |
| P10630 | IF4A2 | Eukaryotic initiation factor 4A-II (eIF-4A-II) (eIF4A-II) (EC 3.6.4.13) (ATP-dependent RNA helicase eIF4A-2) |
| P60229 | EIF3E | Eukaryotic translation initiation factor 3 subunit E (eIF3e) (Eukaryotic translation initiation factor 3 subunit 6) (MMTV integration site 6) (Mammary tumor-associated protein INT-6) (Viral integration site protein INT-6) (eIF-3 p48) |
| Q8QZY1 | EIF3L | Eukaryotic translation initiation factor 3 subunit L (eIF3l) (66 kDa tyrosine-rich heat shock protein) (67 kDa polymerase-associated factor) (Eukaryotic translation initiation factor 3 subunit 6-interacting protein) (Eukaryotic translation initiation factor 3 subunit E-interacting protein) (HSP-66Y) (PAF67) |
| P56564 | EAA1 | Excitatory amino acid transporter 1 (Glial high affinity glutamate transporter) (High-affinity neuronal glutamate transporter) (GluT-1) (Sodium-dependent glutamate/aspartate transporter 1) (GLAST-1) (Solute carrier family 1 member 3) |
| Q6P5F9 | XPO1 | Exportin-1 (Exp1) (Chromosome region maintenance 1 protein homolog) |
| Q9ERK4 | XPO2 | Exportin-2 (Exp2) (Chromosome segregation 1-like protein) (Importin-alpha re-exporter) |
| P26040 | EZRI | Ezrin (Cytovillin) (Villin-2) (p81) |
| P47753 | CAZA1 | F-actin-capping protein subunit alpha-1 (CapZ alpha-1) |
| P47754 | CAZA2 | F-actin-capping protein subunit alpha-2 (CapZ alpha-2) |
| Q3U0V1 | FUBP2 | Far upstream element-binding protein 2 (FUSE-binding protein 2) (KH type-splicing regulatory protein) (KSRP) |
| Q920E5 | FPPS | Farnesyl pyrophosphate synthase (FPP synthase) (FPS) (EC 2.5.1.10) ((2E,6E)-farnesyl diphosphate synthase) (Cholesterol-regulated 39 kDa protein) (CR 39) (Dimethylallyltranstransferase) (EC 2.5.1.1) (Farnesyl diphosphate synthase) (Geranyltranstransferase) |
| P19096 | FAS | Fatty acid synthase (EC 2.3.1.85) [Includes: [Acyl-carrier-protein] S-acetyltransferase (EC 2.3.1.38); [Acyl-carrier-protein] S-malonyltransferase (EC 2.3.1.39); 3-oxoacyl-[acyl-carrier-protein] synthase (EC 2.3.1.41); 3-oxoacyl-[acyl-carrier-protein] reductase (EC 1.1.1.100); 3-hydroxyacyl-[acyl-carrier-protein] dehydratase (EC 4.2.1.59); Enoyl-[acyl-carrier-protein] reductase (EC 1.3.1.39); Oleoyl-[acyl-carrier-protein] hydrolase (EC 3.1.2.14)] |
| A3KGK3 | FR1L4 | Fer-1-like protein 4 |
| A2AMT1 | BFSP1 | Filensin (Beaded filament structural protein 1) (Lens fiber cell beaded-filament structural protein CP 95) (CP95) |
| P05064 | ALDOA | Fructose-bisphosphate aldolase A (EC 4.1.2.13) (Aldolase 1) (Muscle-type aldolase) |
| P97807 | FUMH | Fumarate hydratase, mitochondrial (Fumarase) (EC 4.2.1.2) (EF-3) |
| Q8R3R8 | GBRL1 | Gamma-aminobutyric acid receptor-associated protein-like 1 (GABA(A) receptor-associated protein-like 1) (Glandular epithelial cell protein 1) (GEC-1) |
| Q8VHL5 | CRGN | Gamma-crystallin N (Gamma-N-crystallin) |
| P28236 | CXA8 | Gap junction alpha-8 protein (Connexin-50) (Cx50) (Lens fiber protein MP70) |
| P13020 | GELS | Gelsolin (Actin-depolymerizing factor) (ADF) (Brevin) |
| Q00612 | G6PD1 | Glucose-6-phosphate 1-dehydrogenase X (G6PD) (EC 1.1.1.49) |
| P06745 | G6PI | Glucose-6-phosphate isomerase (GPI) (EC 5.3.1.9) (Autocrine motility factor) (AMF) (Neuroleukin) (NLK) (Phosphoglucose isomerase) (PGI) (Phosphohexose isomerase) (PHI) |
| P26443 | DHE3 | Glutamate dehydrogenase 1, mitochondrial (GDH 1) (EC 1.4.1.3) |
| P97494 | GSH1 | Glutamate--cysteine ligase catalytic subunit (EC 6.3.2.2) (GCS heavy chain) (Gamma-ECS) (Gamma-glutamylcysteine synthetase) |
| P15105 | GLNA | Glutamine synthetase (GS) (EC 6.3.1.2) (Glutamate--ammonia ligase) (Palmitoyltransferase GLUL) (EC 2.3.1.225) |
| Q9CQM9 | GLRX3 | Glutaredoxin-3 (PKC-interacting cousin of thioredoxin) (PICOT) (PKC-theta-interacting protein) (PKCq-interacting protein) (Thioredoxin-like protein 2) |
| P48774 | GSTM5 | Glutathione S-transferase Mu 5 (EC 2.5.1.18) (Fibrous sheath component 2) (Fsc2) (GST class-mu 5) |
| P51855 | GSHB | Glutathione synthetase (GSH synthetase) (GSH-S) (EC 6.3.2.3) (Glutathione synthase) |
| Q3ULJ0 | GPD1L | Glycerol-3-phosphate dehydrogenase 1-like protein (EC 1.1.1.8) |
| Q9CZD3 | GARS | Glycine--tRNA ligase (EC 3.6.1.17) (EC 6.1.1.14) (Diadenosine tetraphosphate synthetase) (AP-4-A synthetase) (Glycyl-tRNA synthetase) (GlyRS) |
| Q9ET01 | PYGL | Glycogen phosphorylase, liver form (EC 2.4.1.1) |
| Q61543 | GSLG1 | Golgi apparatus protein 1 (E-selectin ligand 1) (ESL-1) (Selel) (Golgi sialoglycoprotein MG-160) |
| P62874 | GBB1 | Guanine nucleotide-binding protein G(I)/G(S)/G(T) subunit beta-1 (Transducin beta chain 1) |
| Q61011 | GBB3 | Guanine nucleotide-binding protein G(I)/G(S)/G(T) subunit beta-3 (Transducin beta chain 3) |
| P21279 | GNAQ | Guanine nucleotide-binding protein G(q) subunit alpha (Guanine nucleotide-binding protein alpha-q) |
| P20612 | GNAT1 | Guanine nucleotide-binding protein G(t) subunit alpha-1 (Transducin alpha-1 chain) |
| P30677 | GNA14 | Guanine nucleotide-binding protein subunit alpha-14 (G alpha-14) (G-protein subunit alpha-14) |
| Q8K0U4 | HS12A | Heat shock 70 kDa protein 12A |
| P17879 | HS71B | Heat shock 70 kDa protein 1B (Heat shock 70 kDa protein 1) (HSP70.1) |
| Q3U2G2 | Q3U2G2 | Heat shock 70 kDa protein 4 (Heat shock protein 4, isoform CRA_a) |
| P48722 | HS74L | Heat shock 70 kDa protein 4L (Heat shock 70-related protein APG-1) (Osmotic stress protein 94) |
| Q61699 | HS105 | Heat shock protein 105 kDa (42 degrees C-HSP) (Heat shock 110 kDa protein) (Heat shock-related 100 kDa protein E7I) (HSP-E7I) |
| Q9CQN1 | TRAP1 | Heat shock protein 75 kDa, mitochondrial (HSP 75) (TNFR-associated protein 1) (Tumor necrosis factor type 1 receptor-associated protein) (TRAP-1) |
| P11499 | HS90B | Heat shock protein HSP 90-beta (Heat shock 84 kDa) (HSP 84) (HSP84) (Tumor-specific transplantation 84 kDa antigen) (TSTA) |
| Q3TEA8 | HP1B3 | Heterochromatin protein 1-binding protein 3 |
| Q99020 | ROAA | Heterogeneous nuclear ribonucleoprotein A/B (hnRNP A/B) (CArG-binding factor-A) (CBF-A) |
| P49312 | ROA1 | Heterogeneous nuclear ribonucleoprotein A1 (hnRNP A1) (HDP-1) (Helix-destabilizing protein) (Single-strand-binding protein) (Topoisomerase-inhibitor suppressed) (hnRNP core protein A1) [Cleaved into: Heterogeneous nuclear ribonucleoprotein A1, N-terminally processed] |
| Q8BG05 | ROA3 | Heterogeneous nuclear ribonucleoprotein A3 (hnRNP A3) |
| Q9Z130 | HNRDL | Heterogeneous nuclear ribonucleoprotein D-like (hnRNP D-like) (hnRNP DL) (JKT41-binding protein) |
| Q60668 | HNRPD | Heterogeneous nuclear ribonucleoprotein D0 (hnRNP D0) (AU-rich element RNA-binding protein 1) |
| O35737 | HNRH1 | Heterogeneous nuclear ribonucleoprotein H (hnRNP H) [Cleaved into: Heterogeneous nuclear ribonucleoprotein H, N-terminally processed] |
| P70333 | HNRH2 | Heterogeneous nuclear ribonucleoprotein H2 (hnRNP H2) (Heterogeneous nuclear ribonucleoprotein H') (hnRNP H') [Cleaved into: Heterogeneous nuclear ribonucleoprotein H2, N-terminally processed] |
| P61979 | HNRPK | Heterogeneous nuclear ribonucleoprotein K (hnRNP K) |
| Q8R081 | HNRPL | Heterogeneous nuclear ribonucleoprotein L (hnRNP L) |
| Q921F4 | HNRLL | Heterogeneous nuclear ribonucleoprotein L-like |
| Q9D0E1 | HNRPM | Heterogeneous nuclear ribonucleoprotein M (hnRNP M) |
| Q00PI9 | HNRL2 | Heterogeneous nuclear ribonucleoprotein U-like protein 2 (MLF1-associated nuclear protein) |
| O88569 | ROA2 | Heterogeneous nuclear ribonucleoproteins A2/B1 (hnRNP A2/B1) |
| Q9Z204 | HNRPC | Heterogeneous nuclear ribonucleoproteins C1/C2 (hnRNP C1/C2) |
| O08528 | HXK2 | Hexokinase-2 (EC 2.7.1.1) (Hexokinase type II) (HK II) |
| P63158 | HMGB1 | High mobility group protein B1 (High mobility group protein 1) (HMG-1) |
| P70288 | HDAC2 | Histone deacetylase 2 (HD2) (EC 3.5.1.98) (YY1 transcription factor-binding protein) |
| P10922 | H10 | Histone H1.0 (Histone H1') (Histone H1(0)) (MyD196) [Cleaved into: Histone H1.0, N-terminally processed] |
| P43275 | H11 | Histone H1.1 (H1 VAR.3) (Histone H1a) (H1a) |
| P15864 | H12 | Histone H1.2 (H1 VAR.1) (H1c) |
| Q64522 | H2A2B | Histone H2A type 2-B (H2a-613A) |
| P0C0S6 | H2AZ | Histone H2A.Z (H2A/z) |
| P10854 | H2B1M | Histone H2B type 1-M (H2B 291B) |
| P84228 | H32 | Histone H3.2 |
| P62806 | H4 | Histone H4 |
| Q99L47 | F10A1 | Hsc70-interacting protein (Hip) (Protein FAM10A1) (Protein ST13 homolog) |
| P00493 | HPRT | Hypoxanthine-guanine phosphoribosyltransferase (HGPRT) (HGPRTase) (EC 2.4.2.8) (HPRT B) |
| O35343 | IMA3 | Importin subunit alpha-3 (Importin alpha Q1) (Qip1) (Karyopherin subunit alpha-4) |
| P70168 | IMB1 | Importin subunit beta-1 (Karyopherin subunit beta-1) (Nuclear factor p97) (Pore targeting complex 97 kDa subunit) (PTAC97) (SCG) |
| Q8BKC5 | IPO5 | Importin-5 (Imp5) (Importin subunit beta-3) (Karyopherin beta-3) (Ran-binding protein 5) (RanBP5) |
| Q9CXY6 | ILF2 | Interleukin enhancer-binding factor 2 (Nuclear factor of activated T-cells 45 kDa) |
| P85094 | ISC2A | Isochorismatase domain-containing protein 2A |
| Q9D6R2 | IDH3A | Isocitrate dehydrogenase [NAD] subunit alpha, mitochondrial (EC 1.1.1.41) (Isocitric dehydrogenase subunit alpha) (NAD(+)-specific ICDH subunit alpha) |
| Q91VA7 | Q91VA7 | Isocitrate dehydrogenase [NAD] subunit, mitochondrial |
| O88844 | IDHC | Isocitrate dehydrogenase [NADP] cytoplasmic (IDH) (EC 1.1.1.42) (Cytosolic NADP-isocitrate dehydrogenase) (IDP) (NADP(+)-specific ICDH) (Oxalosuccinate decarboxylase) |
| O54983 | CRYM | Ketimine reductase mu-crystallin (EC 1.5.1.25) (NADP-regulated thyroid-hormone-binding protein) |
| Q3THB4 | Q3THB4 | L-lactate dehydrogenase (EC 1.1.1.27) |
| P16125 | LDHB | L-lactate dehydrogenase B chain (LDH-B) (EC 1.1.1.27) (LDH heart subunit) (LDH-H) |
| Q66VB7 | Q66VB7 | Lacrein (Predicted gene 1553) |
| Q9CPU0 | LGUL | Lactoylglutathione lyase (EC 4.4.1.5) (Aldoketomutase) (Glyoxalase I) (Glx I) (Ketone-aldehyde mutase) (Methylglyoxalase) (S-D-lactoylglutathione methylglyoxal lyase) |
| P14733 | LMNB1 | Lamin-B1 |
| P51180 | MIP | Lens fiber major intrinsic protein (Aquaporin-0) (MIP26) (MP26) |
| Q61735 | CD47 | Leukocyte surface antigen CD47 (Integrin-associated protein) (IAP) (CD antigen CD47) |
| P24527 | LKHA4 | Leukotriene A-4 hydrolase (LTA-4 hydrolase) (EC 3.3.2.6) (Leukotriene A(4) hydrolase) |
| P32067 | LA | Lupus La protein homolog (La autoantigen homolog) (La ribonucleoprotein) |
| P14152 | MDHC | Malate dehydrogenase, cytoplasmic (EC 1.1.1.37) (Cytosolic malate dehydrogenase) |
| P08249 | MDHM | Malate dehydrogenase, mitochondrial (EC 1.1.1.37) |
| Q8K310 | MATR3 | Matrin-3 |
| A2A4X6 | A2A4X6 | MCG21910 (Predicted gene 12355) |
| P45952 | ACADM | Medium-chain specific acyl-CoA dehydrogenase, mitochondrial (MCAD) (EC 1.3.8.7) |
| O55022 | PGRC1 | Membrane-associated progesterone receptor component 1 (mPR) |
| P14873 | MAP1B | Microtubule-associated protein 1B (MAP-1B) (MAP1(X)) (MAP1.2) [Cleaved into: MAP1B heavy chain; MAP1 light chain LC1] |
| P20357 | MTAP2 | Microtubule-associated protein 2 (MAP-2) |
| P10637 | TAU | Microtubule-associated protein tau (Neurofibrillary tangle protein) (Paired helical filament-tau) (PHF-tau) |
| Q9CR62 | M2OM | Mitochondrial 2-oxoglutarate/malate carrier protein (OGCP) (Solute carrier family 25 member 11) |
| Q791V5 | MTCH2 | Mitochondrial carrier homolog 2 |
| Q9D023 | MPC2 | Mitochondrial pyruvate carrier 2 (Brain protein 44) |
| Q6A0D0 | Q6A0D0 | MKIAA0106 protein (Fragment) |
| Q9DCL9 | PUR6 | Multifunctional protein ADE2 [Includes: Phosphoribosylaminoimidazole-succinocarboxamide synthase (EC 6.3.2.6) (SAICAR synthetase); Phosphoribosylaminoimidazole carboxylase (EC 4.1.1.21) (AIR carboxylase) (AIRC)] |
| Q8C854 | MYEF2 | Myelin expression factor 2 (MEF-2) (MyEF-2) |
| Q60605 | MYL6 | Myosin light polypeptide 6 (17 kDa myosin light chain) (LC17) (Myosin light chain 3) (MLC-3) (Myosin light chain alkali 3) (Myosin light chain A3) (Smooth muscle and nonmuscle myosin light chain alkali 6) |
| Q8VDD5 | MYH9 | Myosin-9 (Cellular myosin heavy chain, type A) (Myosin heavy chain 9) (Myosin heavy chain, non-muscle IIa) (Non-muscle myosin heavy chain A) (NMMHC-A) (Non-muscle myosin heavy chain IIa) (NMMHC II-a) (NMMHC-IIA) |
| B1AR69 | B1AR69 | Myosin, heavy polypeptide 13, skeletal muscle |
| Q9DCJ9 | NPL | N-acetylneuraminate lyase (NALase) (EC 4.1.3.3) (N-acetylneuraminate pyruvate-lyase) (N-acetylneuraminic acid aldolase) (Sialate lyase) (Sialate-pyruvate lyase) (Sialic acid aldolase) (Sialic acid lyase) |
| Q99KK2 | NEUA | N-acylneuraminate cytidylyltransferase (EC 2.7.7.43) (CMP-N-acetylneuraminic acid synthase) (CMP-NeuNAc synthase) |
| O35683 | NDUA1 | NADH dehydrogenase [ubiquinone] 1 alpha subcomplex subunit 1 (Complex I-MWFE) (CI-MWFE) (NADH-ubiquinone oxidoreductase MWFE subunit) |
| Q99LC3 | NDUAA | NADH dehydrogenase [ubiquinone] 1 alpha subcomplex subunit 10, mitochondrial (Complex I-42kD) (CI-42kD) (NADH-ubiquinone oxidoreductase 42 kDa subunit) |
| Q9DC69 | NDUA9 | NADH dehydrogenase [ubiquinone] 1 alpha subcomplex subunit 9, mitochondrial (Complex I-39kD) (CI-39kD) (NADH-ubiquinone oxidoreductase 39 kDa subunit) |
| Q9CQ54 | NDUC2 | NADH dehydrogenase [ubiquinone] 1 subunit C2 (Complex I-B14.5b) (CI-B14.5b) (NADH-ubiquinone oxidoreductase subunit B14.5b) |
| Q9DCT2 | NDUS3 | NADH dehydrogenase [ubiquinone] iron-sulfur protein 3, mitochondrial (EC 1.6.99.3) (EC 7.1.1.2) (Complex I-30kD) (CI-30kD) (NADH-ubiquinone oxidoreductase 30 kDa subunit) |
| Q8K3J1 | NDUS8 | NADH dehydrogenase [ubiquinone] iron-sulfur protein 8, mitochondrial (EC 1.6.99.3) (EC 7.1.1.2) (Complex I-23kD) (CI-23kD) (NADH-ubiquinone oxidoreductase 23 kDa subunit) |
| P03921 | NU5M | NADH-ubiquinone oxidoreductase chain 5 (EC 7.1.1.2) (NADH dehydrogenase subunit 5) |
| P13595 | NCAM1 | Neural cell adhesion molecule 1 (N-CAM-1) (NCAM-1) (CD antigen CD56) |
| Q810U3 | NFASC | Neurofascin |
| Q810U4 | NRCAM | Neuronal cell adhesion molecule (Nr-CAM) (Neuronal surface protein Bravo) (mBravo) (NgCAM-related cell adhesion molecule) (Ng-CAM-related) |
| Q8BHN3 | GANAB | Neutral alpha-glucosidase AB (EC 3.2.1.84) (Alpha-glucosidase 2) (Glucosidase II subunit alpha) |
| P10493 | NID1 | Nidogen-1 (NID-1) (Entactin) |
| Q99K48 | NONO | Non-POU domain-containing octamer-binding protein (NonO protein) |
| P09405 | NUCL | Nucleolin (Protein C23) |
| Q61937 | NPM | Nucleophosmin (NPM) (Nucleolar phosphoprotein B23) (Nucleolar protein NO38) (Numatrin) |
| P15532 | NDKA | Nucleoside diphosphate kinase A (NDK A) (NDP kinase A) (EC 2.7.4.6) (Metastasis inhibition factor NM23) (NDPK-A) (Tumor metastatic process-associated protein) (nm23-M1) |
| P28656 | NP1L1 | Nucleosome assembly protein 1-like 1 (Brain protein DN38) (NAP-1-related protein) |
| Q78ZA7 | NP1L4 | Nucleosome assembly protein 1-like 4 |
| Q9CZ30 | OLA1 | Obg-like ATPase 1 (GTP-binding protein 9) |
| P29758 | OAT | Ornithine aminotransferase, mitochondrial (EC 2.6.1.13) (Ornithine--oxo-acid aminotransferase) |
| Q99JF8 | PSIP1 | PC4 and SFRS1-interacting protein (Lens epithelium-derived growth factor) (mLEDGF) |
| P15499 | PRPH2 | Peripherin-2 (Retinal degeneration slow protein) |
| P35700 | PRDX1 | Peroxiredoxin-1 (EC 1.11.1.15) (Macrophage 23 kDa stress protein) (Osteoblast-specific factor 3) (OSF-3) (Thioredoxin peroxidase 2) (Thioredoxin-dependent peroxide reductase 2) |
| Q61171 | PRDX2 | Peroxiredoxin-2 (EC 1.11.1.15) (Thiol-specific antioxidant protein) (TSA) (Thioredoxin peroxidase 1) (Thioredoxin-dependent peroxide reductase 1) |
| Q6NVD9 | BFSP2 | Phakinin (49 kDa cytoskeletal protein) (Beaded filament structural protein 2) (Lens fiber cell beaded filament protein CP 49) (CP49) |
| Q9WUA2 | SYFB | Phenylalanine--tRNA ligase beta subunit (EC 6.1.1.20) (Phenylalanyl-tRNA synthetase beta subunit) (PheRS) |
| Q9QW08 | PHOS | Phosducin (PHD) (33 kDa phototransducing protein) (Rod photoreceptor 1) (RPR-1) |
| O70172 | PI42A | Phosphatidylinositol 5-phosphate 4-kinase type-2 alpha (EC 2.7.1.149) (1-phosphatidylinositol 5-phosphate 4-kinase 2-alpha) (Diphosphoinositide kinase 2-alpha) (Phosphatidylinositol 5-phosphate 4-kinase type II alpha) (PI(5)P 4-kinase type II alpha) (PIP4KII-alpha) (PtdIns(5)P-4-kinase isoform 2-alpha) |
| Q7M6Y3 | PICAL | Phosphatidylinositol-binding clathrin assembly protein (Clathrin assembly lymphoid myeloid leukemia) (CALM) |
| Q9D0F9 | PGM1 | Phosphoglucomutase-1 (PGM 1) (EC 5.4.2.2) (Glucose phosphomutase 1) (Phosphoglucomutase-2) |
| Q8BZF8 | PGM5 | Phosphoglucomutase-like protein 5 |
| P09411 | PGK1 | Phosphoglycerate kinase 1 (EC 2.7.2.3) |
| Q9DBJ1 | PGAM1 | Phosphoglycerate mutase 1 (EC 5.4.2.11) (EC 5.4.2.4) (BPG-dependent PGAM 1) (Phosphoglycerate mutase isozyme B) (PGAM-B) |
| Q80X80 | C2C2L | Phospholipid transfer protein C2CD2L (C2 domain-containing protein 2-like) (Transmembrane protein 24) |
| G5E829 | AT2B1 | Plasma membrane calcium-transporting ATPase 1 (PMCA1) (EC 7.2.2.10) (Plasma membrane calcium ATPase isoform 1) (Plasma membrane calcium pump isoform 1) |
| P63005 | LIS1 | Platelet-activating factor acetylhydrolase IB subunit alpha (Lissencephaly-1 protein) (LIS-1) (PAF acetylhydrolase 45 kDa subunit) (PAF-AH 45 kDa subunit) (PAF-AH alpha) (PAFAH alpha) |
| Q61206 | PA1B2 | Platelet-activating factor acetylhydrolase IB subunit beta (EC 3.1.1.47) (PAF acetylhydrolase 30 kDa subunit) (PAF-AH 30 kDa subunit) (PAF-AH subunit beta) (PAFAH subunit beta) |
| Q61205 | PA1B3 | Platelet-activating factor acetylhydrolase IB subunit gamma (EC 3.1.1.47) (PAF acetylhydrolase 29 kDa subunit) (PAF-AH 29 kDa subunit) (PAF-AH subunit gamma) (PAFAH subunit gamma) |
| Q61990 | PCBP2 | Poly(rC)-binding protein 2 (Alpha-CP2) (CTBP) (CBP) (Putative heterogeneous nuclear ribonucleoprotein X) (hnRNP X) |
| P29341 | PABP1 | Polyadenylate-binding protein 1 (PABP-1) (Poly(A)-binding protein 1) |
| Q8CCS6 | PABP2 | Polyadenylate-binding protein 2 (PABP-2) (Poly(A)-binding protein 2) (Nuclear poly(A)-binding protein 1) (Poly(A)-binding protein II) (PABII) (Polyadenylate-binding nuclear protein 1) |
| E9Q035 | E9Q035 | Predicted gene 20425 |
| Q61656 | DDX5 | Probable ATP-dependent RNA helicase DDX5 (EC 3.6.4.13) (DEAD box RNA helicase DEAD1) (mDEAD1) (DEAD box protein 5) (RNA helicase p68) |
| P67778 | PHB | Prohibitin (B-cell receptor-associated protein 32) (BAP 32) |
| O35129 | PHB2 | Prohibitin-2 (B-cell receptor-associated protein BAP37) (Repressor of estrogen receptor activity) |
| Q9R1P0 | PSA4 | Proteasome subunit alpha type-4 (EC 3.4.25.1) (Macropain subunit C9) (Multicatalytic endopeptidase complex subunit C9) (Proteasome component C9) (Proteasome subunit L) |
| Q9QUM9 | PSA6 | Proteasome subunit alpha type-6 (EC 3.4.25.1) (Macropain iota chain) (Multicatalytic endopeptidase complex iota chain) (Proteasome iota chain) |
| O09061 | PSB1 | Proteasome subunit beta type-1 (EC 3.4.25.1) (Macropain subunit C5) (Multicatalytic endopeptidase complex subunit C5) (Proteasome component C5) (Proteasome gamma chain) |
| Q9R1P3 | PSB2 | Proteasome subunit beta type-2 (EC 3.4.25.1) (Macropain subunit C7-I) (Multicatalytic endopeptidase complex subunit C7-I) (Proteasome component C7-I) |
| P27773 | PDIA3 | Protein disulfide-isomerase A3 (EC 5.3.4.1) (58 kDa glucose-regulated protein) (58 kDa microsomal protein) (p58) (Disulfide isomerase ER-60) (Endoplasmic reticulum resident protein 57) (ER protein 57) (ERp57) (Endoplasmic reticulum resident protein 60) (ER protein 60) (ERp60) |
| Q99LT0 | DPY30 | Protein dpy-30 homolog (Dpy-30-like protein) (Dpy-30L) |
| Q4VA93 | Q4VA93 | Protein kinase C (EC 2.7.11.13) |
| Q61644 | PACN1 | Protein kinase C and casein kinase substrate in neurons protein 1 (Syndapin-1) |
| Q8K1M3 | Q8K1M3 | Protein kinase, cAMP dependent regulatory, type II alpha (Protein kinase, cAMP dependent regulatory, type II alpha, isoform CRA_b) (cAMP-dependent protein kinase type II-alpha regulatory chain) (cAMP-dependent protein kinase type II-alpha regulatory subunit) |
| Q62433 | NDRG1 | Protein NDRG1 (N-myc downstream-regulated gene 1 protein) (Protein Ndr1) |
| Q9QYG0 | NDRG2 | Protein NDRG2 (N-myc downstream-regulated gene 2 protein) (Protein Ndr2) |
| O55126 | NIPS2 | Protein NipSnap homolog 2 (NipSnap2) (Glioblastoma-amplified sequence) |
| Q3UM45 | PP1R7 | Protein phosphatase 1 regulatory subunit 7 (Protein phosphatase 1 regulatory subunit 22) |
| Q80TL0 | PPM1E | Protein phosphatase 1E (EC 3.1.3.16) (Ca(2+)/calmodulin-dependent protein kinase phosphatase N) (CaMKP-N) (CaMKP-nucleus) (CaMKN) (Partner of PIX 1) (Partner of PIX-alpha) (Partner of PIXA) |
| Q8BVQ5 | PPME1 | Protein phosphatase methylesterase 1 (PME-1) (EC 3.1.1.89) |
| Q9D394 | RUFY3 | Protein RUFY3 (Rap2-interacting protein x) (RIPx) (Single axon-regulated protein 1) (Singar1) |
| Q9EQU5 | SET | Protein SET (Phosphatase 2A inhibitor I2PP2A) (I-2PP2A) (Template-activating factor I) (TAF-I) |
| Q99LX0 | PARK7 | Protein/nucleic acid deglycase DJ-1 (EC 3.1.2.-) (EC 3.5.1.-) (EC 3.5.1.124) (Maillard deglycase) (Parkinson disease protein 7 homolog) (Parkinsonism-associated deglycase) (Protein DJ-1) (DJ-1) |
| Q11011 | PSA | Puromycin-sensitive aminopeptidase (PSA) (EC 3.4.11.14) (Cytosol alanyl aminopeptidase) (AAP-S) |
| P35486 | ODPA | Pyruvate dehydrogenase E1 component subunit alpha, somatic form, mitochondrial (EC 1.2.4.1) (PDHE1-A type I) |
| Q9D051 | ODPB | Pyruvate dehydrogenase E1 component subunit beta, mitochondrial (PDHE1-B) (EC 1.2.4.1) |
| P52480 | KPYM | Pyruvate kinase PKM (EC 2.7.1.40) (Pyruvate kinase muscle isozyme) |
| P50396 | GDIA | Rab GDP dissociation inhibitor alpha (Rab GDI alpha) (Guanosine diphosphate dissociation inhibitor 1) (GDI-1) |
| E9Q6Q4 | E9Q6Q4 | RAP1, GTP-GDP dissociation stimulator 1 |
| P53994 | RAB2A | Ras-related protein Rab-2A |
| P63011 | RAB3A | Ras-related protein Rab-3A |
| P68040 | RACK1 | Receptor of activated protein C kinase 1 (12-3) (Guanine nucleotide-binding protein subunit beta-2-like 1) (Receptor for activated C kinase) (Receptor of activated protein kinase C 1) (p205) [Cleaved into: Receptor of activated protein C kinase 1, N-terminally processed (Guanine nucleotide-binding protein subunit beta-2-like 1, N-terminally processed)] |
| P34057 | RECO | Recoverin (23 kDa photoreceptor cell-specific protein) (Cancer-associated retinopathy protein) (Protein CAR) |
| Q9EPU0 | RENT1 | Regulator of nonsense transcripts 1 (EC 3.6.4.-) (ATP-dependent helicase RENT1) (Nonsense mRNA reducing factor 1) (NORF1) (Up-frameshift suppressor 1 homolog) (mUpf1) |
| P24549 | AL1A1 | Retinal dehydrogenase 1 (RALDH 1) (RalDH1) (EC 1.2.1.-) (EC 1.2.1.36) (ALDH-E1) (ALHDII) (Aldehyde dehydrogenase family 1 member A1) (Aldehyde dehydrogenase, cytosolic) |
| O35600 | ABCA4 | Retinal-specific ATP-binding cassette transporter (ATP-binding cassette sub-family A member 4) (RIM ABC transporter) (RIM protein) (RmP) |
| Q9Z275 | RLBP1 | Retinaldehyde-binding protein 1 (Cellular retinaldehyde-binding protein) |
| Q8BYK4 | RDH12 | Retinol dehydrogenase 12 (EC 1.1.1.300) |
| P49194 | RET3 | Retinol-binding protein 3 (Interphotoreceptor retinoid-binding protein) (IRBP) (Interstitial retinol-binding protein) |
| Q9Z1L4 | XLRS1 | Retinoschisin (X-linked juvenile retinoschisis protein homolog) |
| Q99PT1 | GDIR1 | Rho GDP-dissociation inhibitor 1 (Rho GDI 1) (GDI-1) (Rho-GDI alpha) |
| P15409 | OPSD | Rhodopsin |
| Q9WVL4 | RK | Rhodopsin kinase (RK) (EC 2.7.11.14) (G protein-coupled receptor kinase 1) |
| Q91VM5 | RMXL1 | RNA binding motif protein, X-linked-like-1 (Heterogeneous nuclear ribonucleoprotein G-like 1) (RNA binding motif protein, X chromosome retrogene) |
| Q9CQE8 | RTRAF | RNA transcription, translation and transport factor protein |
| Q8VH51 | RBM39 | RNA-binding protein 39 (Coactivator of activating protein 1 and estrogen receptors) (Coactivator of AP-1 and ERs) (RNA-binding motif protein 39) (RNA-binding region-containing protein 2) (Transcription coactivator CAPER) |
| P32958 | ROM1 | Rod outer segment membrane protein 1 (ROSP1) |
| P20443 | ARRS | S-arrestin (48 kDa protein) (Retinal S-antigen) (S-AG) (Rod photoreceptor arrestin) |
| Q9R0P3 | ESTD | S-formylglutathione hydrolase (FGH) (EC 3.1.2.12) (Esterase 10) (Esterase D) (Sid 478) |
| Q91WD9 | SEGN | Secretagogin |
| O55131 | 7-Sep | Septin-7 (CDC10 protein homolog) |
| P26638 | SYSC | Serine--tRNA ligase, cytoplasmic (EC 6.1.1.11) (Seryl-tRNA synthetase) (SerRS) (Seryl-tRNA(Ser/Sec) synthetase) |
| Q6PDM2 | SRSF1 | Serine/arginine-rich splicing factor 1 (ASF/SF2) (Pre-mRNA-splicing factor SRp30a) (Splicing factor, arginine/serine-rich 1) |
| Q9R0U0 | SRS10 | Serine/arginine-rich splicing factor 10 (FUS-interacting serine-arginine-rich protein 1) (Neural-salient serine/arginine-rich protein) (Neural-specific SR protein) (Splicing factor, arginine/serine-rich 13A) (TLS-associated protein with Ser-Arg repeats) (TASR) (TLS-associated protein with SR repeats) (TLS-associated serine-arginine protein) (TLS-associated SR protein) |
| Q76MZ3 | 2AAA | Serine/threonine-protein phosphatase 2A 65 kDa regulatory subunit A alpha isoform (PP2A subunit A isoform PR65-alpha) (PP2A subunit A isoform R1-alpha) |
| P63328 | PP2BA | Serine/threonine-protein phosphatase 2B catalytic subunit alpha isoform (EC 3.1.3.16) (CAM-PRP catalytic subunit) (Calmodulin-dependent calcineurin A subunit alpha isoform) (CNA alpha) |
| P62141 | PP1B | Serine/threonine-protein phosphatase PP1-beta catalytic subunit (PP-1B) (EC 3.1.3.16) (EC 3.1.3.53) |
| Q91V61 | SFXN3 | Sideroflexin-3 |
| P62305 | RUXE | Small nuclear ribonucleoprotein E (snRNP-E) (Sm protein E) (Sm-E) (SmE) |
| P62320 | SMD3 | Small nuclear ribonucleoprotein Sm D3 (Sm-D3) (snRNP core protein D3) |
| P31648 | SC6A1 | Sodium- and chloride-dependent GABA transporter 1 (GAT-1) (Solute carrier family 6 member 1) |
| Q6PIC6 | AT1A3 | Sodium/potassium-transporting ATPase subunit alpha-3 (Na(+)/K(+) ATPase alpha-3 subunit) (EC 7.2.2.13) (Na(+)/K(+) ATPase alpha(III) subunit) (Sodium pump subunit alpha-3) |
| P14094 | AT1B1 | Sodium/potassium-transporting ATPase subunit beta-1 (Sodium/potassium-dependent ATPase subunit beta-1) |
| P14231 | AT1B2 | Sodium/potassium-transporting ATPase subunit beta-2 (Adhesion molecule in glia) (AMOG) (Sodium/potassium-dependent ATPase subunit beta-2) |
| P97370 | AT1B3 | Sodium/potassium-transporting ATPase subunit beta-3 (Sodium/potassium-dependent ATPase subunit beta-3) (ATPB-3) (CD antigen CD298) |
| Q5XG69 | F169A | Soluble lamin-associated protein of 75 kDa (SLAP75) (Protein FAM169A) |
| Q91V14 | S12A5 | Solute carrier family 12 member 5 (Electroneutral potassium-chloride cotransporter 2) (K-Cl cotransporter 2) (mKCC2) (Neuronal K-Cl cotransporter) |
| Q924N4 | S12A6 | Solute carrier family 12 member 6 (Electroneutral potassium-chloride cotransporter 3) (K-Cl cotransporter 3) |
| P17809 | GTR1 | Solute carrier family 2, facilitated glucose transporter member 1 (Glucose transporter type 1, erythrocyte/brain) (GLUT-1) (GT1) |
| Q6P069 | SORCN | Sorcin |
| O70492 | SNX3 | Sorting nexin-3 (SDP3 protein) |
| A3KGU7 | A3KGU7 | Spectrin alpha chain, non-erythrocytic 1 |
| Q62261 | SPTB2 | Spectrin beta chain, non-erythrocytic 1 (Beta-II spectrin) (Embryonic liver fodrin) (Fodrin beta chain) |
| Q9Z1N5 | DX39B | Spliceosome RNA helicase Ddx39b (EC 3.6.4.13) (56 kDa U2AF65-associated protein) (DEAD box protein UAP56) (HLA-B-associated transcript 1 protein) |
| Q99NB9 | SF3B1 | Splicing factor 3B subunit 1 (Pre-mRNA-splicing factor SF3b 155 kDa subunit) (SF3b155) (Spliceosome-associated protein 155) (SAP 155) |
| P26369 | U2AF2 | Splicing factor U2AF 65 kDa subunit (U2 auxiliary factor 65 kDa subunit) (U2 snRNP auxiliary factor large subunit) |
| Q8VIJ6 | SFPQ | Splicing factor, proline- and glutamine-rich (DNA-binding p52/p100 complex, 100 kDa subunit) (Polypyrimidine tract-binding protein-associated-splicing factor) (PSF) (PTB-associated-splicing factor) |
| P38647 | GRP75 | Stress-70 protein, mitochondrial (75 kDa glucose-regulated protein) (GRP-75) (Heat shock 70 kDa protein 9) (Mortalin) (Peptide-binding protein 74) (PBP74) (p66 MOT) |
| Q60864 | STIP1 | Stress-induced-phosphoprotein 1 (STI1) (mSTI1) (Hsc70/Hsp90-organizing protein) (Hop) |
| Q8K2B3 | SDHA | Succinate dehydrogenase [ubiquinone] flavoprotein subunit, mitochondrial (EC 1.3.5.1) (Flavoprotein subunit of complex II) (Fp) |
| Q9Z2I9 | SUCB1 | Succinate--CoA ligase [ADP-forming] subunit beta, mitochondrial (EC 6.2.1.5) (ATP-specific succinyl-CoA synthetase subunit beta) (A-SCS) (Succinyl-CoA synthetase beta-A chain) (SCS-betaA) |
| Q8BWF0 | SSDH | Succinate-semialdehyde dehydrogenase, mitochondrial (EC 1.2.1.24) (Aldehyde dehydrogenase family 5 member A1) (NAD(+)-dependent succinic semialdehyde dehydrogenase) |
| Q9D0K2 | SCOT1 | Succinyl-CoA:3-ketoacid coenzyme A transferase 1, mitochondrial (EC 2.8.3.5) (3-oxoacid CoA-transferase 1) (Somatic-type succinyl-CoA:3-oxoacid CoA-transferase) (SCOT-s) |
| P09671 | SODM | Superoxide dismutase [Mn], mitochondrial (EC 1.15.1.1) |
| Q9JIS5 | SV2A | Synaptic vesicle glycoprotein 2A (Synaptic vesicle protein 2) (Synaptic vesicle protein 2A) (Calcium regulator SV2A) |
| Q62465 | VAT1 | Synaptic vesicle membrane protein VAT-1 homolog (EC 1.-.-.-) |
| Q80TB8 | VAT1L | Synaptic vesicle membrane protein VAT-1 homolog-like (EC 1.-.-.-) |
| P46096 | SYT1 | Synaptotagmin-1 (Synaptotagmin I) (SytI) (p65) |
| O08599 | STXB1 | Syntaxin-binding protein 1 (Protein unc-18 homolog 1) (Unc18-1) (Protein unc-18 homolog A) (Unc-18A) |
| P80314 | TCPB | T-complex protein 1 subunit beta (TCP-1-beta) (CCT-beta) |
| P80315 | TCPD | T-complex protein 1 subunit delta (TCP-1-delta) (A45) (CCT-delta) |
| P80316 | TCPE | T-complex protein 1 subunit epsilon (TCP-1-epsilon) (CCT-epsilon) |
| P80318 | TCPG | T-complex protein 1 subunit gamma (TCP-1-gamma) (CCT-gamma) (Matricin) (mTRiC-P5) |
| P42932 | TCPQ | T-complex protein 1 subunit theta (TCP-1-theta) (CCT-theta) |
| P80317 | TCPZ | T-complex protein 1 subunit zeta (TCP-1-zeta) (CCT-zeta-1) |
| Q921F2 | TADBP | TAR DNA-binding protein 43 (TDP-43) |
| O88746 | TOM1 | Target of Myb protein 1 |
| Q91W90 | TXND5 | Thioredoxin domain-containing protein 5 (Endoplasmic reticulum resident protein 46) (ER protein 46) (ERp46) (Plasma cell-specific thioredoxin-related protein) (PC-TRP) (Thioredoxin-like protein p46) |
| P20108 | PRDX3 | Thioredoxin-dependent peroxide reductase, mitochondrial (EC 1.11.1.15) (Antioxidant protein 1) (AOP-1) (PRX III) (Perioredoxin-3) (Protein MER5) |
| O08583 | THOC4 | THO complex subunit 4 (Tho4) (Ally of AML-1 and LEF-1) (Aly/REF export factor) (REF1-I) (RNA and export factor-binding protein 1) (Transcriptional coactivator Aly/REF) |
| Q9D0R2 | SYTC | Threonine--tRNA ligase, cytoplasmic (EC 6.1.1.3) (Threonyl-tRNA synthetase) (ThrRS) |
| Q5SRX1 | TM1L2 | TOM1-like protein 2 (Target of Myb-like protein 2) |
| Q921T2 | TOIP1 | Torsin-1A-interacting protein 1 (Lamina-associated polypeptide 1B) (LAP1B) |
| Q62318 | TIF1B | Transcription intermediary factor 1-beta (TIF1-beta) (E3 SUMO-protein ligase TRIM28) (EC 2.3.2.27) (KRAB-A-interacting protein) (KRIP-1) (RING-type E3 ubiquitin transferase TIF1-beta) (Tripartite motif-containing protein 28) |
| P42669 | PURA | Transcriptional activator protein Pur-alpha (Purine-rich single-stranded DNA-binding protein alpha) |
| O35295 | PURB | Transcriptional activator protein Pur-beta (Purine-rich element-binding protein B) (Vascular actin single-stranded DNA-binding factor 2 p44 component) |
| Q6PFR5 | TRA2A | Transformer-2 protein homolog alpha (TRA-2 alpha) (TRA2-alpha) (Transformer-2 protein homolog A) |
| Q9QUI0 | RHOA | Transforming protein RhoA |
| Q01853 | TERA | Transitional endoplasmic reticulum ATPase (TER ATPase) (EC 3.6.4.6) (15S Mg(2+)-ATPase p97 subunit) (Valosin-containing protein) (VCP) |
| P40142 | TKT | Transketolase (TK) (EC 2.2.1.1) (P68) |
| Q62186 | SSRD | Translocon-associated protein subunit delta (TRAP-delta) (Signal sequence receptor subunit delta) (SSR-delta) |
| P17751 | TPIS | Triosephosphate isomerase (TIM) (EC 5.3.1.1) (Methylglyoxal synthase) (EC 4.2.3.3) (Triose-phosphate isomerase) |
| P68368 | TBA4A | Tubulin alpha-4A chain (Alpha-tubulin 4) (Alpha-tubulin isotype M-alpha-4) (Tubulin alpha-4 chain) |
| Q9D6F9 | TBB4A | Tubulin beta-4A chain (Tubulin beta-4 chain) |
| Q62376 | RU17 | U1 small nuclear ribonucleoprotein 70 kDa (U1 snRNP 70 kDa) (U1-70K) (snRNP70) |
| Q6P4T2 | U520 | U5 small nuclear ribonucleoprotein 200 kDa helicase (EC 3.6.4.13) (BRR2 homolog) (U5 snRNP-specific 200 kDa protein) (U5-200KD) |
| P56399 | UBP5 | Ubiquitin carboxyl-terminal hydrolase 5 (EC 3.4.19.12) (Deubiquitinating enzyme 5) (Isopeptidase T) (Ubiquitin thioesterase 5) (Ubiquitin-specific-processing protease 5) |
| Q9D2M8 | UB2V2 | Ubiquitin-conjugating enzyme E2 variant 2 (Ubc-like protein MMS2) |
| Q02053 | UBA1 | Ubiquitin-like modifier-activating enzyme 1 (EC 6.2.1.45) (Ubiquitin-activating enzyme E1) (Ubiquitin-activating enzyme E1 X) (Ubiquitin-like modifier-activating enzyme 1 X) |
| Q3TJL8 | Q3TJL8 | Uncharacterized protein |
| Q8BPF4 | Q8BPF4 | Uncharacterized protein |
| Q3U5M8 | Q3U5M8 | Uncharacterized protein |
| Q3U8W9 | Q3U8W9 | Uncharacterized protein |
| Q3U995 | Q3U995 | Uncharacterized protein |
| Q3TFD9 | Q3TFD9 | Uncharacterized protein |
| Q9Z1G4 | VPP1 | V-type proton ATPase 116 kDa subunit a isoform 1 (V-ATPase 116 kDa isoform a1) (Clathrin-coated vesicle/synaptic vesicle proton pump 116 kDa subunit) (Vacuolar adenosine triphosphatase subunit Ac116) (Vacuolar proton pump subunit 1) (Vacuolar proton translocating ATPase 116 kDa subunit a isoform 1) |
| P50516 | VATA | V-type proton ATPase catalytic subunit A (V-ATPase subunit A) (EC 7.1.2.2) (V-ATPase 69 kDa subunit) (Vacuolar proton pump subunit alpha) |
| P62814 | VATB2 | V-type proton ATPase subunit B, brain isoform (V-ATPase subunit B 2) (Endomembrane proton pump 58 kDa subunit) (Vacuolar proton pump subunit B 2) |
| P50518 | VATE1 | V-type proton ATPase subunit E 1 (V-ATPase subunit E 1) (V-ATPase 31 kDa subunit) (p31) (Vacuolar proton pump subunit E 1) |
| Q9CR51 | VATG1 | V-type proton ATPase subunit G 1 (V-ATPase subunit G 1) (V-ATPase 13 kDa subunit 1) (Vacuolar proton pump subunit G 1) |
| Q9Z1Q9 | SYVC | Valine--tRNA ligase (EC 6.1.1.9) (Protein G7a) (Valyl-tRNA synthetase) (ValRS) |
| O70503 | DHB12 | Very-long-chain 3-oxoacyl-CoA reductase (EC 1.1.1.330) (17-beta-hydroxysteroid dehydrogenase 12) (17-beta-HSD 12) (3-ketoacyl-CoA reductase) (KAR) (Estradiol 17-beta-dehydrogenase 12) (EC 1.1.1.62) (KIK-I) |
| P63044 | VAMP2 | Vesicle-associated membrane protein 2 (VAMP-2) (Synaptobrevin-2) |
| P46460 | NSF | Vesicle-fusing ATPase (EC 3.6.4.6) (N-ethylmaleimide-sensitive fusion protein) (NEM-sensitive fusion protein) (Suppressor of K(+) transport growth defect 2) (Protein SKD2) (Vesicular-fusion protein NSF) |
| O35633 | VIAAT | Vesicular inhibitory amino acid transporter (GABA and glycine transporter) (Solute carrier family 32 member 1) (Vesicular GABA transporter) (mVGAT) (mVIAAT) |
| Q60932 | VDAC1 | Voltage-dependent anion-selective channel protein 1 (VDAC-1) (mVDAC1) (Outer mitochondrial membrane protein porin 1) (Plasmalemmal porin) (Voltage-dependent anion-selective channel protein 5) (VDAC-5) (mVDAC5) |
| Q60930 | VDAC2 | Voltage-dependent anion-selective channel protein 2 (VDAC-2) (mVDAC2) (Outer mitochondrial membrane protein porin 2) (Voltage-dependent anion-selective channel protein 6) (VDAC-6) (mVDAC6) |
