## Supplemental Table 12 for "Metabolite therapy guided by liquid biopsy proteomics delays retinal neurodegeneration"

**Table S12.** Pathway representation of down-regulated proteins in the *Pde6ɑ^D670G^* retina.

| **Pathway** | **Fold Enrichment** | **p-value** | **FDR** |
| --- | --- | --- | --- |
| Aminobutyrate degradation | 62.28 | 1.46E-03 | 1.60E-02 |
| Glutamine glutamate conversion | 37.37 | 2.07E-04 | 3.39E-03 |
| ATP synthesis | 35.59 | 1.85E-05 | 6.08E-04 |
| Pentose phosphate pathway | 31.14 | 2.75E-05 | 7.51E-04 |
| Asparagine and aspartate biosynthesis | 31.14 | 3.58E-03 | 3.26E-02 |
| Pyruvate metabolism | 25.95 | 5.07E-06 | 2.08E-04 |
| TCA cycle | 22.65 | 7.29E-05 | 1.50E-03 |
| Glycolysis | 22.42 | 2.06E-09 | 1.69E-07 |
| Gamma-aminobutyric acid synthesis | 20.76 | 6.55E-03 | 4.47E-02 |
| Pyrimidine Metabolism | 18.68 | 9.95E-04 | 1.26E-02 |
| Rod outer segment phototransduction | 13.47 | 4.66E-07 | 2.55E-05 |
| Axon guidance mediated by semaphorins | 10.83 | 8.07E-04 | 1.10E-02 |
| De novo purine biosynthesis | 8.9 | 2.77E-05 | 6.48E-04 |
| Synaptic vesicle trafficking | 8.9 | 1.55E-03 | 1.59E-02 |
| Cortocotropin releasing factor receptor signaling pathway | 8.65 | 4.50E-04 | 6.71E-03 |
| Angiotensin II-stimulated signaling through G proteins and beta-arrestin | 6.56 | 4.28E-03 | 3.51E-02 |
| Histamine H1 receptor mediated signaling pathway | 5.66 | 6.90E-03 | 4.53E-02 |
| Oxytocin receptor mediated signaling pathway | 5.37 | 3.18E-03 | 3.07E-02 |
| Muscarinic acetylcholine receptor 1 and 3 signaling pathway | 5.19 | 3.64E-03 | 3.15E-02 |
| Thyrotropin-releasing hormone receptor signaling pathway | 4.72 | 5.32E-03 | 3.97E-02 |
| 5HT2 type receptor mediated signaling pathway | 4.65 | 5.65E-03 | 4.03E-02 |
| Parkinson disease | 4.49 | 1.33E-03 | 1.56E-02 |
| Huntington disease | 4.13 | 1.24E-04 | 2.27E-03 |
| Inflammation mediated by chemokine and cytokine signaling pathway | 2.58 | 4.81E-03 | 3.75E-02 |
