## Supplemental Table 13 for "Metabolite therapy guided by liquid biopsy proteomics delays retinal neurodegeneration"

**Table S13.** Down-regulated proteins in the *Pde6ɑ^D670G^* retina involved in oxidative phosphorylation.

| **Symbol** | **Protein Name** | **p-value** | **Fold Change** |
| --- | --- | --- | --- |
| ATP5A1 | ATP synthase, mitochondrial F1 complex, alpha subunit 1 | 2.06E-13 | -128372 |
| ATP5B | ATP synthase, mitochondrial F1 complex, beta polypeptide | 3.97E-07 | -26.908 |
| ATP5C1 | ATP synthase, mitochondrial F1 complex, gamma polypeptide | 1.80E-02 | -329.308 |
| ATP5F1 | ATP synthase, mitochondrial Fo complex subunit B1 | 2.29E-03 | -6952.05 |
| ATP5H | ATP synthase, mitochondrial Fo complex subunit D | 2.27E-11 | -15810.3 |
| ATP5J2 | ATP synthase, mitochondrial Fo complex subunit F2 | 3.63E-12 | -18504.3 |
| ATP5L | ATP synthase, mitochondrial Fo complex subunit G | 1.02E-09 | -1803.99 |
| ATP5O | ATP synthase, mitochondrial F1 complex, O subunit | 1.88E-04 | -38887.2 |
| COX4I1 | cytochrome c oxidase subunit 4I1 | 1.39E-11 | -9966.55 |
| COX5A | cytochrome c oxidase subunit 5A | 4.42E-10 | -4717.69 |
| COX6B1 | cytochrome c oxidase subunit 6B1 | 1.61E-11 | -8962.81 |
| CYC1 | cytochrome c1 | 4.84E-03 | -1000 |
| CYCS | cytochrome c, somatic | 7.26E-13 | -3914.87 |
| MT-CO2 | cytochrome c oxidase subunit II | 3.11E-13 | -23633.3 |
| MT-ND5 | NADH dehydrogenase, subunit 5 (complex I) | 2.40E-03 | -1247.32 |
| NDUFA1 | NADH:ubiquinone oxidoreductase subunit A1 | 1.01E-03 | -3634.24 |
| NDUFA4 | NDUFA4, mitochondrial complex associated | 5.23E-04 | -14833.7 |
| NDUFA5 | NADH:ubiquinone oxidoreductase subunit A5 | 1.46E-03 | -2289.43 |
| NDUFA9 | NADH:ubiquinone oxidoreductase subunit A9 | 7.51E-04 | -5517.85 |
| NDUFA10 | NADH:ubiquinone oxidoreductase subunit A10 | 2.72E-09 | -6952.05 |
| NDUFA11 | NADH:ubiquinone oxidoreductase subunit A11 | 1.84E-03 | -1817.12 |
| NDUFA13 | NADH:ubiquinone oxidoreductase subunit A13 | 3.72E-04 | -23944.3 |
| NDUFB5 | NADH:ubiquinone oxidoreductase subunit B5 | 2.83E-03 | -1000 |
| NDUFB6 | NADH:ubiquinone oxidoreductase subunit B6 | 7.69E-04 | -8276.77 |
| NDUFB10 | NADH:ubiquinone oxidoreductase subunit B10 | 7.98E-04 | -5277.63 |
| NDUFS1 | NADH:ubiquinone oxidoreductase core subunit S1 | 1.03E-11 | -5517.85 |
| NDUFS2 | NADH:ubiquinone oxidoreductase core subunit S2 | 3.05E-13 | -4578.86 |
| NDUFS3 | NADH:ubiquinone oxidoreductase core subunit S3 | 2.19E-10 | -9252.13 |
| NDUFS4 | NADH:ubiquinone oxidoreductase subunit S4 | 2.22E-02 | -100 |
| NDUFS7 | NADH:ubiquinone oxidoreductase core subunit S7 | 1.48E-09 | -2620.74 |
| NDUFS8 | NADH:ubiquinone oxidoreductase core subunit S8 | 1.40E-14 | -6603.85 |
| NDUFV1 | NADH:ubiquinone oxidoreductase core subunit V1 | 9.64E-04 | -3914.87 |
| SDHA | succinate dehydrogenase complex flavoprotein subunit A | 4.48E-04 | -13276.1 |
| SDHB | succinate dehydrogenase complex iron sulfur subunit B | 9.64E-04 | -3914.87 |
| SDHC | succinate dehydrogenase complex subunit C | 1.35E-07 | -3419.95 |
| UQCR10 | ubiquinol-cytochrome c reductase, complex III subunit X | 5.49E-09 | -6257.32 |
| UQCRB | ubiquinol-cytochrome c reductase binding protein | 3.97E-11 | -2519.84 |
| UQCRC1 | ubiquinol-cytochrome c reductase core protein I | 3.66E-13 | -24449.8 |
| UQCRC2 | ubiquinol-cytochrome c reductase core protein II | 1.12E-10 | -4578.86 |
| UQCRFS1 | ubiquinol-cytochrome c reductase, Rieske iron-sulfur polypeptide 1 | 3.56E-11 | -1000 |
