## Supplemental Table 14 for "Metabolite therapy guided by liquid biopsy proteomics delays retinal neurodegeneration"

**Table S14.** Down-regulated proteins in the *Pde6ɑ^D670G^* retina involved in the tricarboxylic acid (TCA) cycle.

| **Symbol** | **Protein Name** | **p-value** | **Fold Change** |
| --- | --- | --- | --- |
| ACO2 | aconitase 2 | 1.27E-14 | -22740.6 |
| CS | citrate synthase | 3.52E-04 | -15323 |
| DLD | dihydrolipoamide dehydrogenase | 6.22E-04 | -7268.48 |
| DLST | dihydrolipoamide S-succinyltransferase | 9.98E-03 | -2289.43 |
| FH | fumarate hydratase | 5.27E-13 | -2620.74 |
| IDH3A | isocitrate dehydrogenase 3 (NAD+) alpha | 1.84E-13 | -33970 |
| IDH3B | isocitrate dehydrogenase 3 (NAD+) beta | 6.53E-11 | -6952.05 |
| IDH3G | isocitrate dehydrogenase 3 (NAD+) gamma | 1.99E-03 | -1587.4 |
| MDH2 | malate dehydrogenase 2 | 1.00E-19 | -133663 |
| OGDH | oxoglutarate dehydrogenase | 8.63E-04 | -12295.9 |
| SDHA | succinate dehydrogenase complex flavoprotein subunit A | 4.48E-04 | -13276.1 |
| SDHB | succinate dehydrogenase complex iron sulfur subunit B | 9.64E-04 | -3914.87 |
| SDHC | succinate dehydrogenase complex subunit C | 1.35E-07 | -3419.95 |
| SUCLA2 | succinate-CoA ligase ADP-forming beta subunit | 7.74E-04 | -5616.1 |
