## Supplemental Table 15 for "Metabolite therapy guided by liquid biopsy proteomics delays retinal neurodegeneration"

**Table S15.** Down-regulated proteins in the *Pde6ɑ^D670G^* retina involved in rod outer segment phototransduction.

| **UniProt** | **Symbol** | **Protein names** |
| --- | --- | --- |
| P62874 | GBB1 | Guanine nucleotide-binding protein G(I)/G(S)/G(T) subunit beta-1 (Transducin beta chain 1) |
| P15409 | OPSD | Rhodopsin |
| P20612 | GNAT1 | Guanine nucleotide-binding protein G(t) subunit alpha-1 (Transducin alpha-1 chain) |
| Q9QW08 | PHOS | Phosducin (PHD) (33 kDa phototransducing protein) (Rod photoreceptor 1) (RPR-1) |
| E1AZ71 | E1AZ71 | Cyclic nucleotide-gated channel beta 1 (cGMP-gated cation channel beta subunit) |
| Q9WVL4 | RK | Rhodopsin kinase (RK) (EC 2.7.11.14) (G protein-coupled receptor kinase 1) |
| P34057 | RECO | Recoverin (23 kDa photoreceptor cell-specific protein) (Cancer-associated retinopathy protein) (Protein CAR) |
| P20443 | ARRS | S-arrestin (48 kDa protein) (Retinal S-antigen) (S-AG) (Rod photoreceptor arrestin) |
