## Supplemental Table 16 for "Metabolite therapy guided by liquid biopsy proteomics delays retinal neurodegeneration"

**Table S16.** Tandem-MS analysis of extracted metabolites from the retinal tissue. # - Tentative assignments are based on high mass accuracy, isotopic distribution, and tandem mass spectrometry (using the collision induced dissociation technique). FA, fatty acid; PA, phosphatidic acid; PG, glycerophosphoglycerol; PS, glycerophosphoserine; PI, glycerophosphoinositol. (A:B/X:Y) denotes the number of carbons (A and X) and the number of double bonds (B and Y) in each of the two fatty-acid chains.

| **Sl. No.** | **Measured m/z*** | **Theoretical m/z^†^** | **Molecular formula** | **Attribution^#^** |
| --- | --- | --- | --- | --- |
| 1 | 87.0085 | 87.0088 | C_3_H_3_O_3_ | Pyruvic acid |
| 2 | 89.0243 | 89.0244 | C_3_H_5_O_3_ | Lactate |
| 3 | 103.0034 | 103.0037 | C_3_H_4_O_4_ | β-Hydroxypyruvic acid/  Malonic acid |
| 4 | 103.0403 | 103.0401 | C_4_H_8_O_3_ | β-Hydroxybutyric acid |
| 5 | 115.0032 | 115.0037 | C_4_H_3_O_4_ | Fumaric acid |
| 6 | 117.0188 | 117.0193 | C_4_H_5_O_4_ | Succinic acid |
| 7 | 124.0072 | 124.0074 | C_2_H_6_NO_3_S | Taurine |
| 8 | 127.0161 | 127.0162 | C_3_H_8_O_3_ | [Glycerol+Cl]^‑^ |
| 9 | 129.0386 | 129.0193 | C_5_H_6_O_4_ | Glutaconic acid |
| 10 | 133.0140 | 133.0142 | C_4_H_5_O_5_ | Malate |
| 11 | 146.0456 | 146.0459 | C_5_H_8_NO_4_ | Glutamate |
| 12 | 173.0088 | 173.0092 | C_6_H_5_O_6_ | Aconitic acid |
| 13 | 174.0403 | 174.0408 | C_6_H_9_NO_5_ | N-acetyl aspartate |
| 14 | 181.0713 | 181.0717 | C_6_H_14_O_6_ | Sorbitol |
| 15 | 217.0469 | 217.0479 | C_6_H_14_O_6_ | [Sorbitol+Cl]^‑^ |
| 16 | 241.0112 | 241.0830 | C_10_H_14_N_2_O_5_ | Thymidine |
| 17 | 243.0612 | 243.0623 | C_9_H_12_N_2_O_6_ | Uridine |
| 18 | 245.0422 | 245.0779 | C_9_H_14_N_2_O_6_ | Dihydrouridine |
| 19 | 249.0210 | 249.0221 | C_4_H_14_N_2_O_6_S | Taurine dimer |
| 20 | 255.2311 | 255.2329 | C_16_H_31_O_2_ | Palmitic acid/FA(16:0) |
| 21 | 281.2466 | 281.2486 | C_18_H_33_O_2_ | Oleic acid/FA(18:1) |
| 22 | 283.2622 | 283.2643 | C_18_H_36_O_2_ | Stearic acid |
| 23 | 303.2307 | 303.2329 | C_20_H_31_O_2_ | Arachidonic acid/FA(20:4) |
| 24 | 311.1673 | 311.2956 | C_20_H_40_O_2_ | Phytanic acid |
| 25 | 327.2305 | 327.2329 | C_22_H_31_O_2_ | Docosahexaenoic acid/FA(22:6) |
| 26 | 339.2071 | 339.2091 | C_20_H_31_O_2_ | [Arachidonic acid+Cl]^-^ |
| 27 | 349.2357 | 349.2384 | C_21_H_34_O_4_ | 11-deoxy-11-methylene-PGD2/9-deoxy-9-methylene-PGE2 |
| 28 | 363.2068 | 363.2091 | C_22_H_31_O_2_ | [Docosahexaenoic acid+Cl]^-^ |
| 29 | 395.2772 | C_23_H_40_O_5_ | C_23_H_40_O_5_ | PGF2α isopropyl ester |
| 30 | 437.3085 | 437.3009 | C_27_H_46_S | [Thiocholesterol+Cl]^-^ |
| 31 | 465.3398 | 465.3499 | C_29_H_50_O_2_ | DL-α-Tocopherol/  Cholesterol sulfate |
| 32 | 511.4692 | 511.4732 | C_32_H_63_O_4_ | Palmitic acid dimer |
| 33 | 537.4846 | 537.4888 | C_34_H_65_O_4_ | [Oleic acid + palmitic acid] |
| 34 | 539.5004 | 539.5045 | C_34_H_68_O_4_ | [Stearic acid + palmitic acid] |
| 35 | 563.5001 | 563.5044 | C_36_H_67_O_4_ | Oleic acid dimer |
| 36 | 585.4843 | 585.4883 | C_38_H_65_O_4_ | [Oleic acid + arachidonic acid] |
| 37 | 587.4999 | 587.5045 | C_38_H_68_O_4_ | [Stearic acid + arachidonic acid] |
| 38 | 607.4684 | 607.4732 | C_38_H_64_NaO_4_ | Arachidonic acid dimer |
| 39 | 631.4684 | 631.4708 | C_35_H_67_O_8_P | PA(16:0/16:1) |
| 40 | 737.4918 | 737.5127 | C_42_H_74_O_8_P | PA(20:4)/(19:0) |
| 41 | 747.5124 | 773.5338 | C_42_H_78_O_10_P | PG(18:2/18:0) |
| 42 | 774.5384 | 774.5079 | C_44_H_73_NO_8_P | PE(17:1/22:6) |
| 43 | 766.5354 | 766.5392 | C_43_H_77_NO_8_P | PE(20:4/18:0) |
| 44 | 790.5329 | 790.5392 | C_45_H_77_NO_8_P | PE(18:0/22:6) |
| 45 | 810.5228 | 810.5291 | C_44_H_77_NO_10_P | PS(18:0/20:4) |
| 46 | 834.5225 | 834.5291 | C_46_H_77_NO_10_P | PS(22:6/18:0) |
| 47 | 857.5116 | 857.5186 | C_45_H_78_O_13_P | PI(20:4/16:0) |
| 48 | 865.4959 | 865.4873 | C_46_H_74_O_13_P | PI(15:1/22:6) |
| 49 | 885.5429 | 885.5499 | C_47_H_82_O_13_P | PI(20:4/18:0) |
